# KinasePhos 3.0: Redesign and Expansion of the Prediction on Kinase-specific Phosphorylation Sites

**DOI:** 10.1101/2021.11.02.467032

**Authors:** Renfei Ma, Shangfu Li, Wenshuo Li, Lantian Yao, Hsien-Da Huang, Tzong-Yi Lee

## Abstract

The purpose of this work is to enhance KinasePhos, a machine-learning-based kinase-specific phosphorylation site prediction tool. Experimentally verified kinase-specific phosphorylation data were collected from PhosphoSitePlus, UniProt, GPS 5.0, and Phospho.ELM. In total, 41,421 experimentally verified kinase-specific phosphorylation sites were identified. A total of 1380 unique kinases were identified, including 753 with existing classification information from KinBase and the remaining 627 annotated by building a phylogenetic tree. Based on this kinase classification, a total of 771 predictive models were built at the individual, family, and group levels, using at least 15 experimentally verified substrate sites in positive training datasets. The improved models were observed to be more effective than other prediction tools. For example, the prediction of sites phosphorylated by the Akt, CKT, and PKA families had accuracies of 94.5%, 92.5%, and 90.0%, respectively. The average prediction accuracy for all 771 models was 87.2%. For enhancing interpretability, the Shapley additive explanations (SHAP) method was employed to assess feature importance. The web interface of KinasePhos 3.0 has been redesigned with the goal of providing comprehensive annotations of kinase-specific phosphorylation sites on multiple proteins. Additionally, considering the large scale of phosphoproteomic data, a downloadable prediction tool is available at https://awi.cuhk.edu.cn/KinasePhos/index.html or https://github.com/tom-209/KinasePhos-3.0-executable-file.

## Introduction

Protein phosphorylation is an important eukaryotic post-translational modification [1]. It involves the transfer of a phosphate group from ATP to specific amino-acid residues in the substrate. Phosphorylation is catalyzed by a number of protein kinases, which regulate a variety of signaling pathways and biological functions important in DNA repair, transcriptional regulation, apoptosis, immune response, signaling, metabolism, proliferation, and differentiation [2-7]. Dysregulation of intracellular phosphorylation networks contributes to the occurrence and development of multiple multifactorial diseases, including cancer, cardiovascular disease, obesity, and others [8-10]. Therefore, regulating phosphorylation networks by mediating kinase activity has become an attractive therapeutic strategy [11] with kinases being one of the most important drug targets [12,13]. Thus, linking dysregulated phosphorylation sites to candidate kinase targets is critical, both for the study of disease mechanisms and the development of therapeutic kinase inhibitors [14,15].

The number of experimentally detected phosphorylated sites has increased dramatically in recent years because of advances in mass spectrometry and new enrichment methods for phosphorylated proteins and peptides [16]. For example, deep phosphoproteome analysis of *Schistosoma mansoni* detected 15,844 unique phosphopeptides mapping to 3,176 proteins [17]. Phosphoproteomics can provide important information about protein phosphorylation sites, but the responsible kinases cannot be directly derived from such data. In fact, the kinases for a vast majority of phosphorylation sites are still unknown due to a lack of adequate evidence [18]. To address this problem, many tools have been developed to predict kinase-specific phosphorylation sites in proteins. For example, PhosphoPredict was developed to predict kinase-specific substrates and their associated phosphorylation sites for 12 human kinases and their families by combining protein sequences and functional features [19]. Neural networks were applied by NetPhos 3.1 to predict phosphorylation sites in eukaryotic proteins for 17 kinases [20]. Quokka was introduced to predict kinase family-specific phosphorylation sites at the proteomic scale in a high-throughput and cost-effective manner [21]. Musite provided a unique method that trained models with a bootstrap aggregating procedure, as well as integrated sequence cluster information around phosphorylation sites, protein-disorder scores, and amino-acid frequencies to predict general and kinase-specific phosphorylation sites [22]. The Group-Based Prediction System (GPS) 5.0 tool employed two novel methods, position-weight determination (PWD) and scoring-matrix optimization (SMO), to replace the motif-length selection (MLS) method for refining the prediction of kinase-specific phosphorylation sites [23]. In addition, the conditional random field (CRF) model (CRPhos) [24] and support vector machines (PredPhospho) have been employed to predict the phosphorylation sites [25]. These tools have made outstanding progress in protein phosphorylation studies.

In 2005, our group developed KinasePhos 1.0 to identify protein kinase-specific phosphorylation sites [26]. This tool constructed models from kinase-specific groups of phosphorylation sites based on the profile hidden Markov model (HMM). Subsequently, support vector machines (SVM) with the protein-sequence profiles and protein-coupling patterns were applied to update the tool to version 2.0 [27]. The datasets available for training are constantly expanding owing to the rapid development of phosphorylation-related research. Therefore, in this study, we introduce KinasePhos 3.0, with improved kinase-specific phosphorylation site prediction. We collected experimental identifications of kinase-specific phosphorylation sites from the PhosphoSitePlus [28], UniProt [29], GPS 5.0 [23], and Phospho.ELM [30] databases. Redundant data were removed after translating the kinase and substrate names into unique UniProt IDs. Finally, 41,421 empirically determined, kinase-specific phosphorylation sites were obtained for use as the training data set, which was a great improvement from the training of version 2.0, which involved 16,543 kinase-specific phosphorylation sites. We also assigned kinases to groups, families, or subfamilies according to sequence similarity and the classification method of KinBase [31]. Then, according to these classifications, we used both SVM and eXtreme Gradient Boosting (XGBoost) algorithms to construct 771 prediction models at the kinase group, kinase family, and individual kinase levels, in contrast to 60 predictive models at the individual kinase level in version 2.0. Using these models, specific phosphorylation sites for ten groups, 81 families, and 302 kinases were identified. We also plotted the Shapley additive explanations (SHAP) values of feature groups for each prediction result, which makes the tool more interpretable than version 2.0, as well as other tools in this field. Using SHAP values, users can subdivide the prediction to show the impact of each feature group—that is, features related to specific residues in this study—on the results. Additionally, a standalone version of KinasePhos 3.0, was developed, making it more convenient for users with larger phosphoproteomic datasets than KinasePhos 2.0.

## Method

### Schematic of the proposed KinasePhos 3.0

Figure 1 depicts a schematic of this study that includes kinase-specific phosphorylation site data collection, kinase group and family classifications, feature extractions, machine learning-based kinase-specific phosphorylation site prediction model development, and presentation of results. The novelties of this study are:

1. To our knowledge, the experimentally verified kinase-specific phosphorylation-site data used in this study are, to date, the most comprehensive compared to all existing kinase-specific phosphorylation site prediction tools, such as GPS 5.0 and Kinasephos 2.0.
2. We obtained 771 prediction models, with at least 15 kinase-specific phosphorylation sites considered in each. Thus, the minimum number of positive sites for a single model was greater than that of some other tools. For example, GPS 5.0 includes prediction models for clusters with no less than three positive sites.
3. To increase the feature interpretability of these prediction models, SHAP was integrated into KinasePhos 3.0.

**Figure 1.**
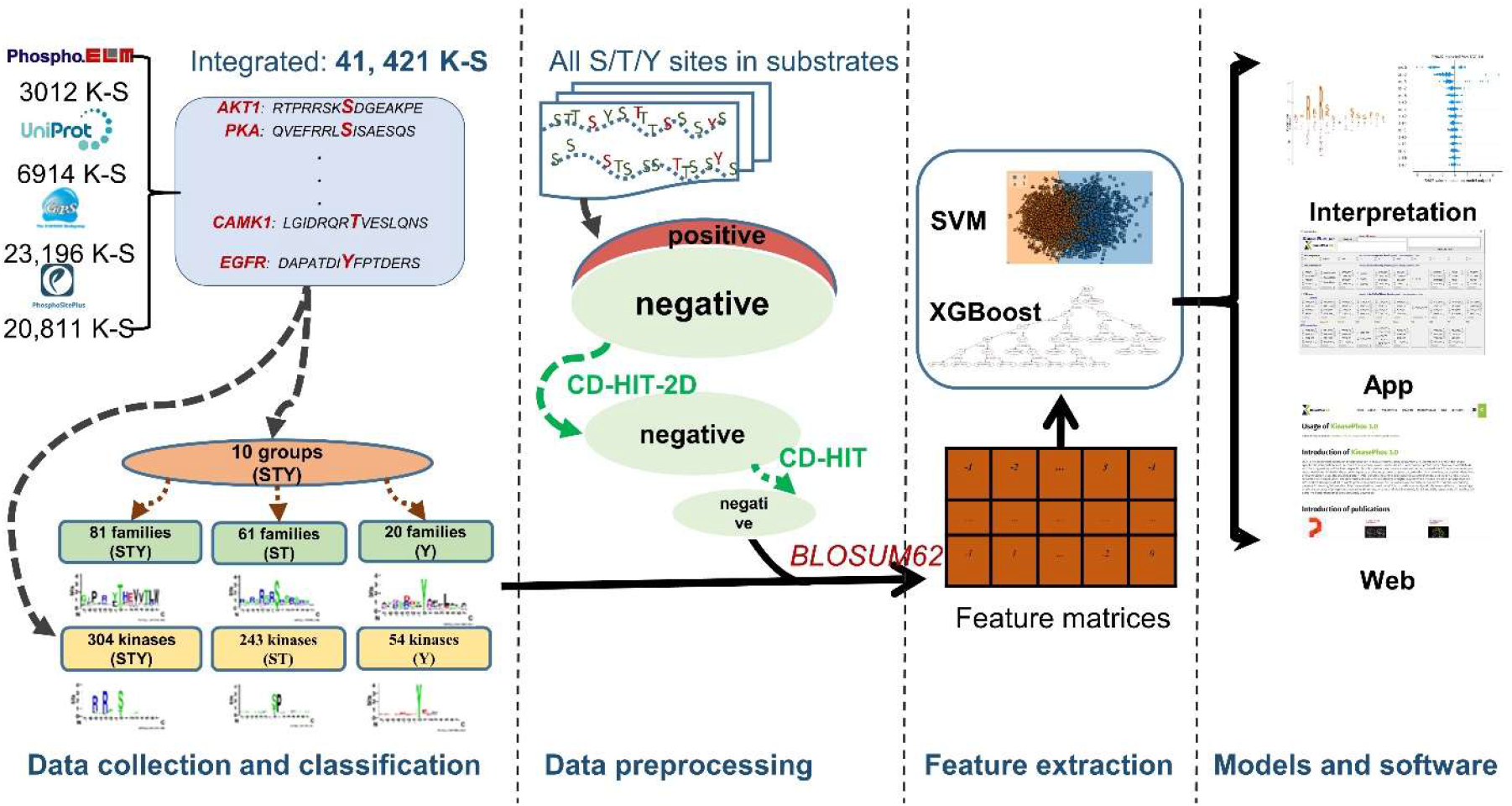
Schematic of KinasePhos 3.0 development. The procedures include data collection, processing, modeling, and website functions development.

### Kinase-substrate data collection

The experimentally verified kinase-specific phosphorylation sites used in this study were collected from four phosphorylation-associated resources: GPS 5.0 [23], Phospho.ELM [30], PhosphoSitePlus [28], and UniProt [29]. Although GPS 5.0, Phospho.ELM and PhosphoSitePlus provided downloadable, experimentally verified, and kinase-specific phosphorylation sites, their data is not frequently updated to reflect the increase in experimentally verified phosphorylated sites. In contrast, UniProt has a standard 8-week release cycle [29]. Therefore, we additionally curated experimentally verified, kinase-specific phosphorylation sites from UniProt with the aim of assembling the most comprehensive database. As depicted in Figure 1, 23,196, 3,012, 20,811, and 6,914 experimentally verified kinase-specific phosphorylation sites were retrieved from GPS 5.0, Phospho.ELM, PhosphoSitePlus, and UniProt, respectively. After eliminating redundancies, 41,421 sites remained, of which, the kinases for 28,369 had UniProt IDs. In contrast, the kinases for the remaining 13,052 sites lack UniProt IDs, primarily because only their kinase family types, instead of kinase names, are provided.

We converted all the kinase names in our substrate dataset into UniProt entry names. Then, we used the classification annotations of kinomes and their sequence information from KinBase as the annotated dataset [31]. By searching the UniProt database, gene names were converted to UniProt IDs. The collected and annotated human kinome datasets were merged and converted to FASTA format. Multiple sequence alignments were performed using the MAFFT program [32]. FastTree was then employed to infer kinetic-maximal-likelihood phylogenetic trees from the kinase sequence alignments [33]. We assumed that homologous proteins have consistent domains represented by closer distances in the phylogenetic tree. Therefore, based on the classification data from KinBase and the generated tree, kinases could be annotated to different clusters at the group, family, and subfamily levels [34]. In addition, we obtained kinase domain data from the PFAM and SMART databases to confirm the results of our classification annotation [35, 36]. TreeGraph 2 and the Interactive Tree Of Life (iTOL) were used to visualize the annotations [34, 37].

### Model development

The classical BLOSUM62 substitute matrix has been widely employed to encode sequence data [23, 27, 38, 39] and was used in this study. For GPS 5.0, the support vector machine (SVM) showed higher performance in kinase-specific phosphorylation site predictive models compared to the random forest (RF) and k-nearest neighbor (KNN) [23] methods. Additionally, eXtreme Gradient Boosting (XGBoost) [40], an efficient implementation of gradient boosted decision trees, is suitable for web server applications for a faster response owing to its model performance and execution speed. Therefore, SVM and XGBoost were used to train the prediction models. The development, testing, and validation of these algorithms were implemented using Python 3.8.

The performance of the kinase-specific, phosphorylation-site prediction models was assessed via classification accuracy and two other metrics, precision and recall, as indicators of reliability. The F1_score, a more comprehensive quantifier of model reliability and the area under the receiver operating characteristic (ROC) curve (AUC) were also computed. These performance measures are defined as:

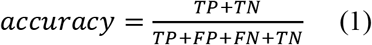

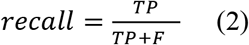

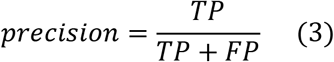

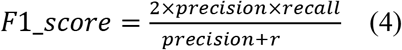

where TP, TN, FP, and FN represent true positives, true negatives, false positives, and false negatives, respectively. Weighted accuracy, weighted recall, weighted precision, and weighted F1_score are the weighted mean of accuracy, recall, precision, and F1_score with weights equal to the class probability, respectively.

### Feature interpretation with SHAP

Because explainable machine learning offers the potential to provide more insights into model behavior, the interpretability of machine-learning models has received significant attention, along with the popularity of machine-learning algorithms. Several feature-importance methods have been developed, including permutation feature importance, which is based on the decrease in model performance and SHAP values [41], which are based on the magnitudes of feature attributions. To increase the interpretability of our prediction models, SHAP was employed to integrate feature importance. SHAP is a game-theory approach and a local explanation to depict the feature’s importance. It has been adopted in some studies [42-44] to interpret machine-learning models. The explanation model can be illustrated by the following equation [41]:

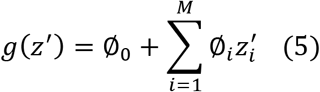

where *z*’ ∈ {0,1}^M^, with 0 and 1 indicating the absence and presence of a feature, respectively. M represents the number of simplified input features. The Shapley value *Ø_i_*, namely the contribution of feature *i*, is calculated as:

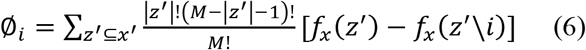

where |*z*’| is the number of non-zeros in *z*’, *z*’\*i* means *z*’ without feature *i, f_x_* is the output of the model, and *x*’ represents simplified inputs.

The SHAP typically evaluates each feature individually; however, in some cases, quantifying the effect of a group of features may be more informative. As mentioned above, the data are 15-mer sequences. In the feature extraction process, the residue at each position was encoded by a 20-dimensional BLOSUM log-odds vector [45]. After being encoded by the BLOSUM62 substitution matrix, the sequences were converted into 300-dimensional (15 × 20) vectors, with each element in a vector representing a feature. Because the amino-acid residue at each position was encoded by a 20-dimensional vector, representing 20 features, these features were clustered as a feature group when performing SHAP analysis, representing a group of features related to specific residues. As a result, 15 feature groups were obtained, corresponding to each position of a 15-mer sequence.

## Results

### Classifying kinases at group and family levels

In total, we obtained 1,380 unique kinases from the kinase-substrate dataset. Of these, 753 were included in the KinBase database, which includes classification information. In contrast, the remaining kinases needed to be annotated by other classification methods. Merging these kinases with the annotated dataset of the human kinome and classifying them by building an evolutionary phylogenetic tree (Figure S1) showed that proteins that are homologous or with consistent domains clustered tightly in smaller branches (Figure 2A), such as *STK10_BOVIN, STK10_HUAMN, STK10_MOUSE*, and *STK10_RAT*. Since *STK10_HUAMN* belongs to the SLK subfamily of the STE20 family of the STE group, we inferred that the other three kinases also belong to that subfamily. Different subfamilies of kinases can form different clusters. For example, for *TAO_DROME, TAOK3_HUMAN, TAOK2_HUAMN, TAOK2_MOUSE, TAOK1_HUAMN, TAOK1_RAT*, and *TAOK1_MOUSE*, although they also belong to the STE20 family of the STE group, the difference in the domain amino acid sequence from the SLK subfamily placed them on another branch belonging to the TAO subfamily. Based on this process, we annotated the collected kinases to groups, families, and subfamilies. Figure 2B shows a kinome tree for several major groups. Analysis of each group separately showed that kinases in the same group contained similar domains (Figure 2C). This indicated that our annotation of the collected kinases was reliable.

**Figure 2.**
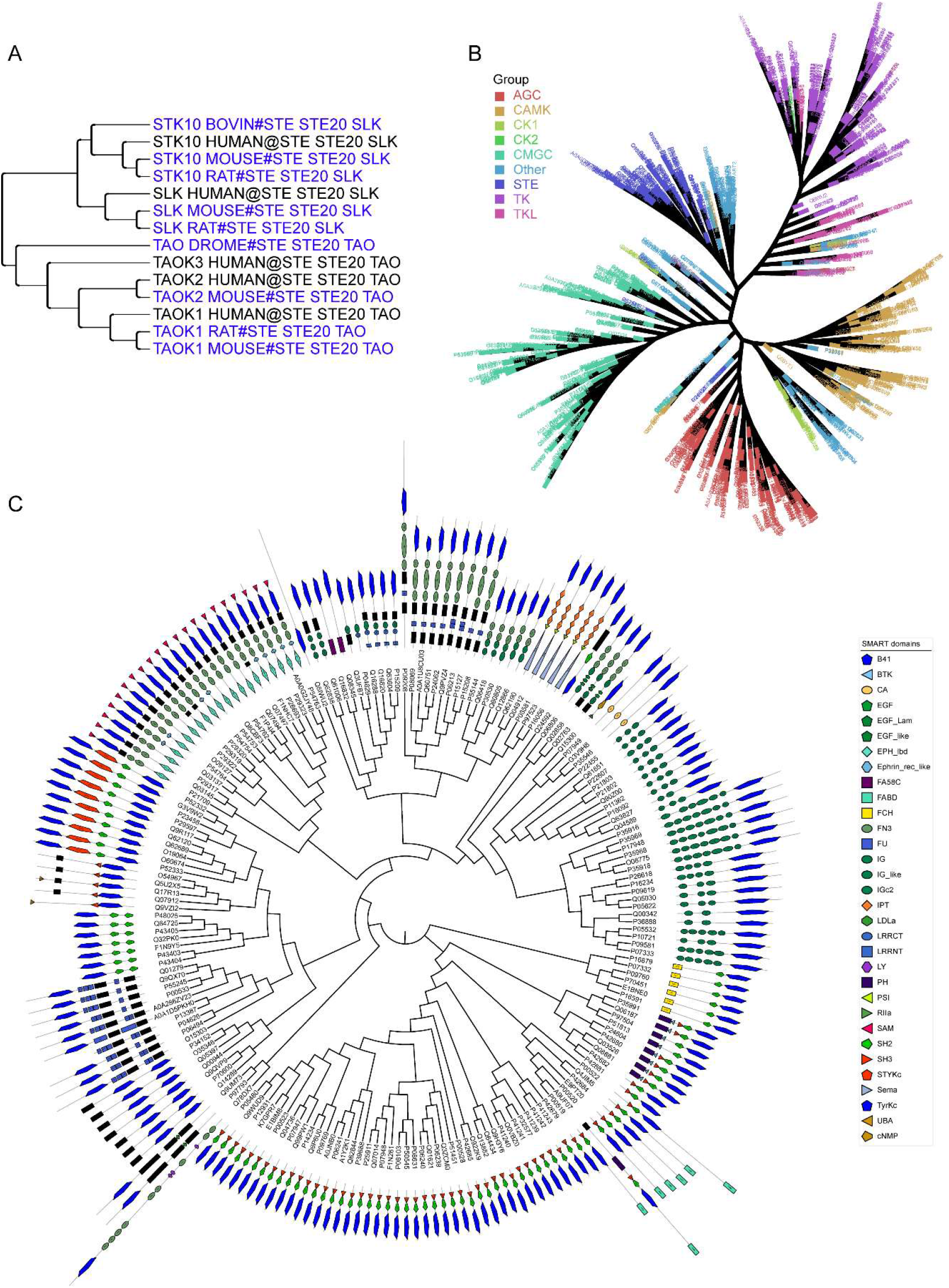
Kinase classification. **A**. Phylogenetic tree of the kinases of the SLK and TAO subfamilies in the STE20 family of the STE group. **B**. Kinome tree composed of several major groups. **C**. Domain annotation of TK group kinases.

Finally, these kinases were classified into 12 kinase groups and 116 kinase families. When we developed our predictive models, only groups or family clusters with at least 15 experimental phosphorylation sites were considered. As a result, ten groups and 81 families were retained. As serine/threonine (S/T) and tyrosine (Y) kinases modify different residues, we developed prediction models for both types separately in family clusters. Similarly, only group or family clusters with at least 15 related sites were considered. Since most substrate residues in the TK group were Y, while most substrate residues in the other nine groups were S/T, they were not separately considered when creating group prediction models. Moreover, we developed prediction models at the individual kinase level for clusters with more than 15 phosphorylation sites, with 11 types of organisms retained. While the majority are human, mice, and rat, others include mouse-ear cress (arath), bovine, chicken, pig, sumatran orangutan (ponab), fission yeast (schpo), African clawed frog (xenla), and yeast. Again, phosphoserine/phosphothreonine and phosphotyrosine sites were considered separately if their number in substrates of a particular kinase was no less than 15. In practice, 15-residue sequences (−7 to +7) surrounding kinase-specific phosphorylation sites were extracted as positive data. After removing redundant sites within each cluster, numbers of clusters and the number ranges for the positive data in each are summarized in **Table 1**. We obtained ten models for the ten group clusters; 81, 61, and 20 models were built for family clusters considering S/T and Y sites, S/T sites, and Y sites, respectively; 302, 243, and 54 models were developed for kinase-specific clusters considering S/T and Y sites, S/T sites, and Y sites, respectively. A total of 771 prediction models were created. In the group clusters, the numbers of positive sites ranged from 204 to 5,737. In the family clusters, the numbers ranged from 15 to 2,050, and the numbers of kinase clusters ranged from 15 to 930. Although clusters with positive sites less than 15 were not considered when developing models, the data for these clusters are included in supplementary files (Table S1) for those who might be interested in them.

**Table 1.** Summary of numbers of prediction models and ranges of positive sites for predictive models.

| Clusters | Model number | Number range of positive data |
| --- | --- | --- |
| group | 10 | 204 – 5737 |
| family_all | 81 | 15 – 2050 |
| family_ST | 61 | 15 – 2046 |
| family_Y | 20 | 18 – 1310 |
| kinase_all | 302 | 15 – 930 |
| kinase_ST | 243 | 15 – 929 |
| kinase_Y | 54 | 15 – 652 |

*Note:* all indicates the S/T and Y sites, ST means S/T sites, and Y refers to Y sites.

In each cluster, all the same types of residues in the phosphorylated substrate proteins, except those known to be positive phosphorylation sites, were regarded as negative data. For example, in family clusters considering all phosphorylated residues (model type family_all listed in Table 1), all S/T and Y sites in all substrate proteins in a cluster were obtained. After eliminating the positive data (*i.e*., experimentally verified phosphorylation sites), the remaining sites were taken as the negative data of that cluster. Similarly, in family clusters considering S/T residues (model type family_ST in Table 1), the negative data are all S and T sites except those sites in the positive data for that cluster. CD-HIT [46] has been widely used to reduce sequence similarity in the literature [19, 47]. Because the number of negative sites obtained via this method is much greater than the number of positive sites, for balance we first used the CD-HIT-2D [46] to reduce the similarity of negative data to positive data with a similarity threshold of 0.4, the minimum threshold in the CD-HIT-2D suite. Furthermore, CD-HIT [46] was employed to further reduce the similarity between the negative data in each cluster. After experimentally applying different threshold values, we found that the number of negative sites is sometimes much greater than the number of positive sites, even though the minimum threshold of 0.4 in the CD-HIT suite was adopted. Suppose the number of negative sites is more than five times greater than the number of positive sites after applying CD-HIT-2D and CD-HIT. In this case, we applied the random undersampling technique from the imbalanced-learn library in Python to keep the number difference within five-fold to reduce the imbalance between positive and negative data when developing the predictive models.

To investigate the characteristics of amino-acid composition in the aforementioned positive 15-mer sequences and provide a graphical representation, we obtained sequence logos of positive sequence clusters for all models using the WebLogo tool (https://weblogo.berkeley.edu/). Some representative logos are shown in Figure 3, which correspond to the ten groups (left two columns) and to some representative families (right two columns). In the common kinase family protein kinase A (PKA), protein kinase C (PKC), protein kinase D (PKD), casein kinase 2 (CK2), cyclin-dependent kinase (CDK), and mitogen-activated protein kinase (MAPK), the majority of phosphorylated sites are S/T residues, as shown in Figure 3. Kinases of some families, such as the focal adhesion kinase (FAK) and serine/threonine-protein kinase STE7 (STE7) families, can phosphorylate both S/T and Y residues. The Abelson kinase (Abl) family and tyrosine kinase (Tec) family clusters mainly correspond to Y sites. More sequence logos of these families and individual kinases are provided in Supplementary Table S2.

**Figure 3.**
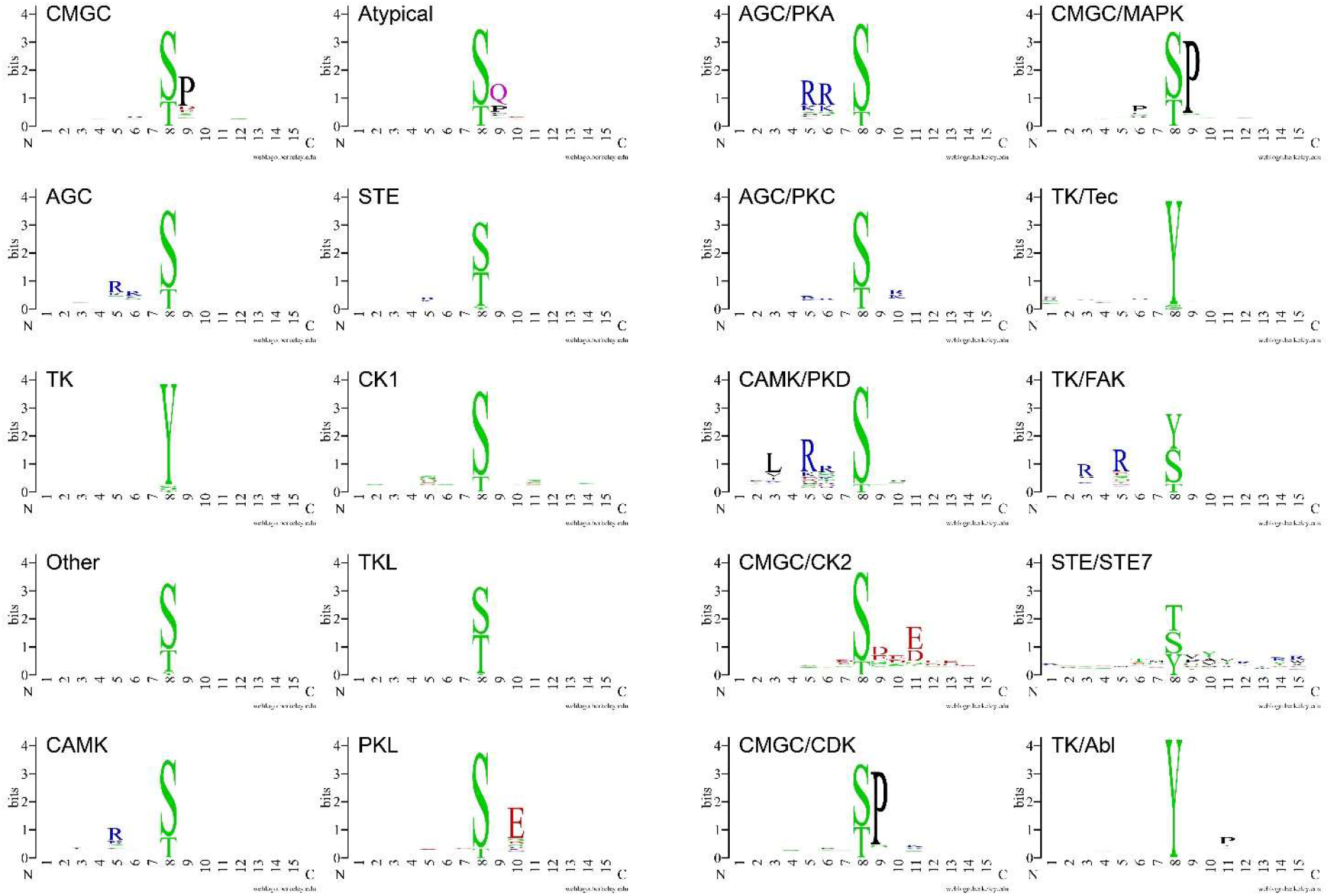
Sequence logos of site clusters of different kinase groups and family clusters. Sequence logos of substrate site clusters phosphorylated by kinases from the CMGC group, AGC group, TK group, Other group, CAMK group, Atypical group, STE group, CK1 group, TKL group, and PKL group, are shown in the left two columns. Those phosphorylated by kinases of the PKA, PKC, PKD, CK2, CDK, MAPK, Tec, FAK, STE7, and Abl families are shown in the right two columns.

### Performance of KinasePhos 3.0 and comparison with other tools

As there are a total of 771 prediction models, to conveniently present their overall performance, average values of accuracy, weighted F1 score, weighted precision, weighted recall, and ROC-AUC for models in each of the seven types of clusters (that is, groups, family_all, family _ST, family_Y, kinase_all, kinase_ST, and kinase_Y, as presented in Table 1) were calculated (**Table 2**). The performance of each model is shown in Supplementary Table S2. It is worth noting that the accuracy, weighted F1 score, weighted precision, weighted recall, and ROC-AUC were generally slightly higher with XGBoost than with SVM. Thus, the XGBoost algorithm was adopted for training our models to develop the website prediction function.

**Table 2.**
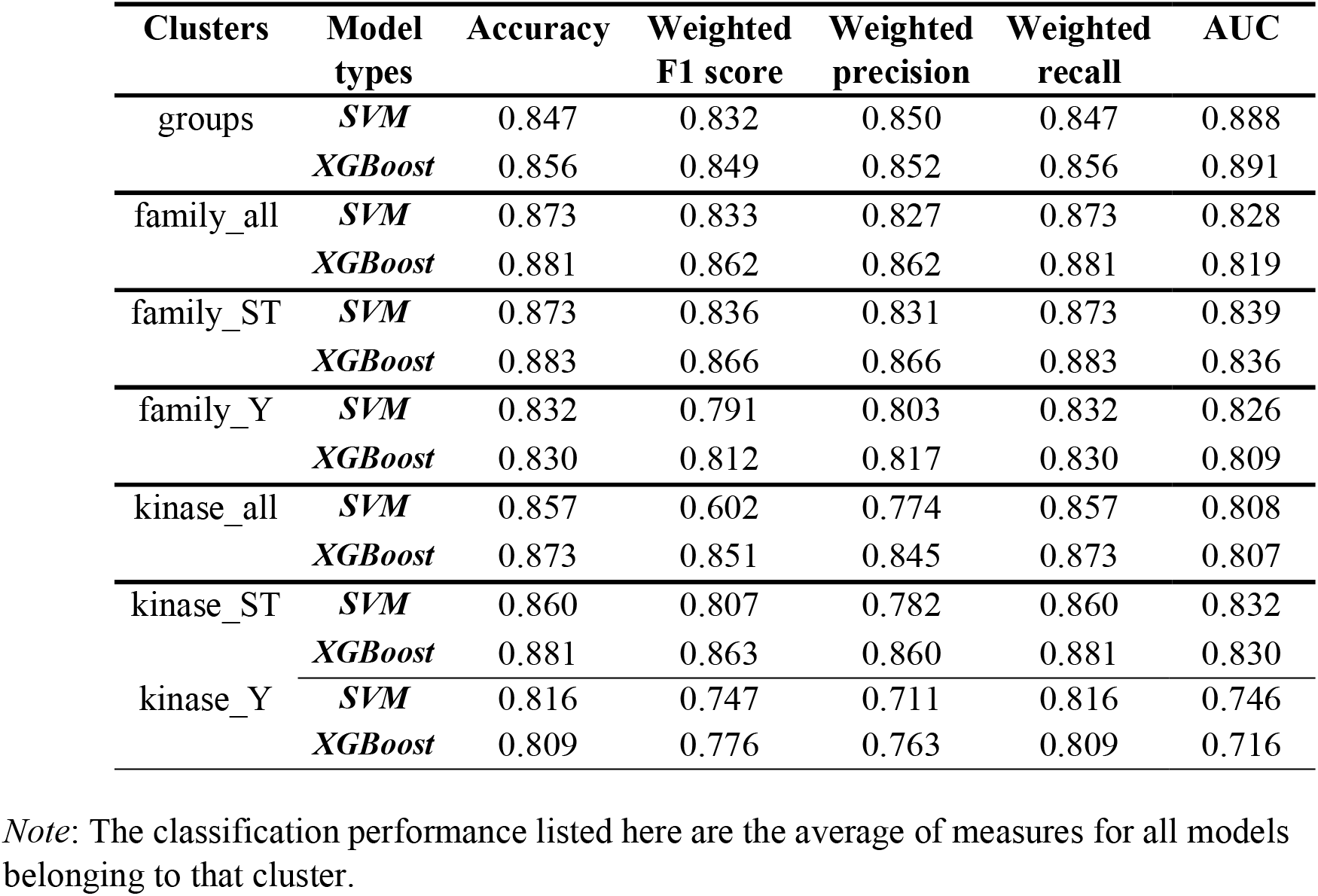
Selected KinasePhos 3.0 performance comparisons with support vector machines (SVM) and eXtreme Gradient Boosting (XGBoost) algorithms.

| Clusters | Model types | Accuracy | Weighted F1 score | Weighted precision | Weighted recall | AUC |
| --- | --- | --- | --- | --- | --- | --- |
| groups | *SVM* | 0.847 | 0.832 | 0.850 | 0.847 | 0.888 |
|  | *XGBoost* | 0.856 | 0.849 | 0.852 | 0.856 | 0.891 |
| family_all | *SVM* | 0.873 | 0.833 | 0.827 | 0.873 | 0.828 |
|  | *XGBoost* | 0.881 | 0.862 | 0.862 | 0.881 | 0.819 |
| family_ST | *SVM* | 0.873 | 0.836 | 0.831 | 0.873 | 0.839 |
|  | *XGBoost* | 0.883 | 0.866 | 0.866 | 0.883 | 0.836 |
| family_Y | *SVM* | 0.832 | 0.791 | 0.803 | 0.832 | 0.826 |
|  | *XGBoost* | 0.830 | 0.812 | 0.817 | 0.830 | 0.809 |
| kinase_all | *SVM* | 0.857 | 0.602 | 0.774 | 0.857 | 0.808 |
|  | *XGBoost* | 0.873 | 0.851 | 0.845 | 0.873 | 0.807 |
| kinase_ST | *SVM* | 0.860 | 0.807 | 0.782 | 0.860 | 0.832 |
|  | *XGBoost* | 0.881 | 0.863 | 0.860 | 0.881 | 0.830 |
| kinase_Y | *SVM* | 0.816 | 0.747 | 0.711 | 0.816 | 0.746 |
|  | *XGBoost* | 0.809 | 0.776 | 0.763 | 0.809 | 0.716 |

Note: The classification performance listed here are the average of measures for all models belonging to that cluster.

To examine these models in more detail, the classification performance of the kinase group models and the numbers of positive and negative sites used to train them are presented in **Table 3.**

**Table 3.** Table 3 Performance of kinase group eXtreme Gradient Boosting (XGBoost) models with 10-fold cross-validation.

| Kinase groups | No. of positive sites | No. of negative sites | Accuracy | Weighted F1 score | Weighted precision | Weighted recall | AUC |
| --- | --- | --- | --- | --- | --- | --- | --- |
| **CMGC** | 5737 | 1470 | 0.943 | 0.943 | 0.944 | 0.943 | 0.982 |
| **AGC** | 4602 | 1632 | 0.901 | 0.901 | 0.901 | 0.901 | 0.958 |
| **TK** | 2680 | 1688 | 0.808 | 0.807 | 0.808 | 0.808 | 0.884 |
| **Other** | 2068 | 1523 | 0.79 | 0.79 | 0.792 | 0.79 | 0.875 |
| **CAMK** | 1892 | 1920 | 0.852 | 0.852 | 0.854 | 0.852 | 0.928 |
| **Atypical** | 1037 | 2004 | 0.886 | 0.88 | 0.888 | 0.886 | 0.935 |
| **STE** | 625 | 1595 | 0.837 | 0.826 | 0.833 | 0.837 | 0.851 |
| **CK1** | 508 | 1402 | 0.857 | 0.849 | 0.854 | 0.857 | 0.888 |
| **TKL** | 360 | 1222 | 0.802 | 0.769 | 0.775 | 0.802 | 0.744 |
| **PKL** | 204 | 1400 | 0.882 | 0.875 | 0.876 | 0.882 | 0.862 |

The new KinasePhos 3.0 was compared with other predictive models, namely KinasePhos 1.0 [26], KinasePhos 2.0 [27], GPS 5.0 [23], ScanSite 4.0 [48], and Net-Phos3.1 [20], using four typical kinase families (CDK, CK2, PKA, and MAPK), selected and compared using GPS 5.0. We found that KinasePhos 3.0 is competitive (**Figure 4**). ROC curves produced by 10-fold cross-validation of KinasePhos 3.0 are presented, with the sensitivity (Sn) and 1-Specificity (Sp) values for the other tools shown as dots with different colors in the plots.

**Figure 4.**
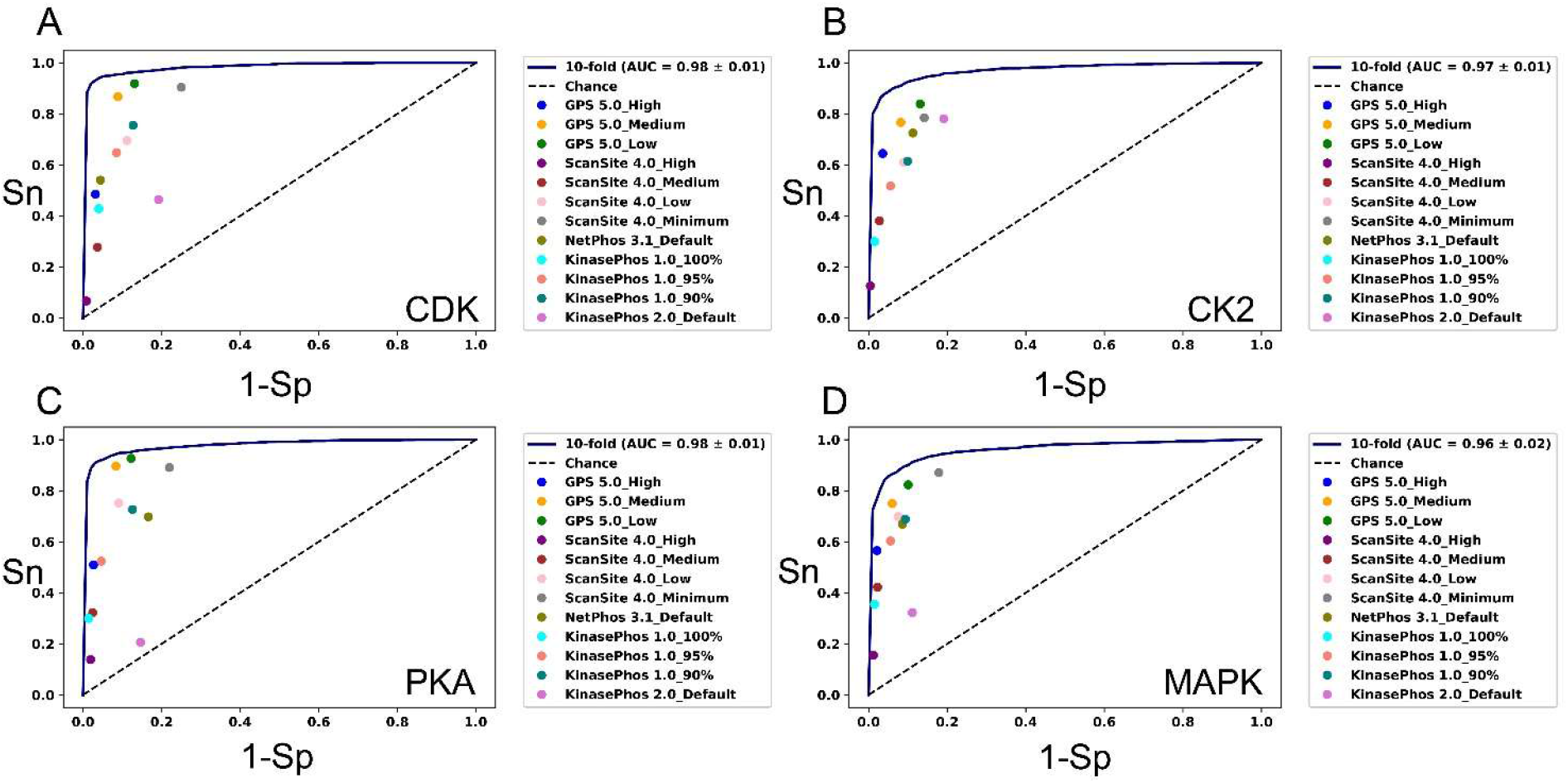
Performance comparisons between KinasePhos 3.0 and existing tools. Existing tools include GPS 5.0 (blue, orange, and green dots), ScanSite 4.0 (purple, brown, pink, and grey dots), NetPhos 3.1 (olive dot), KinasePhos 1.0 (cyan, salmon, and teal dots), and KinasePhos 2.0 (orchid dot). Models include those for the (**A**) CDK, (**B**) CK2, (**C**) PKA, and (**D**) MAPK families.

When k-fold cross-validation was applied, an optimization investigation of k for cross-validation with k=4, 6, 8, and 10 was performed (**Figure 5**), which includes the CDK, CK2, PKD, and Tec families and compares the performance with ROC curves and AUC values. We found that the selection of k did not have a significant impact on performance; thus, the commonly used 10-fold cross-validation was adopted for presenting performance.

**Figure 5.**
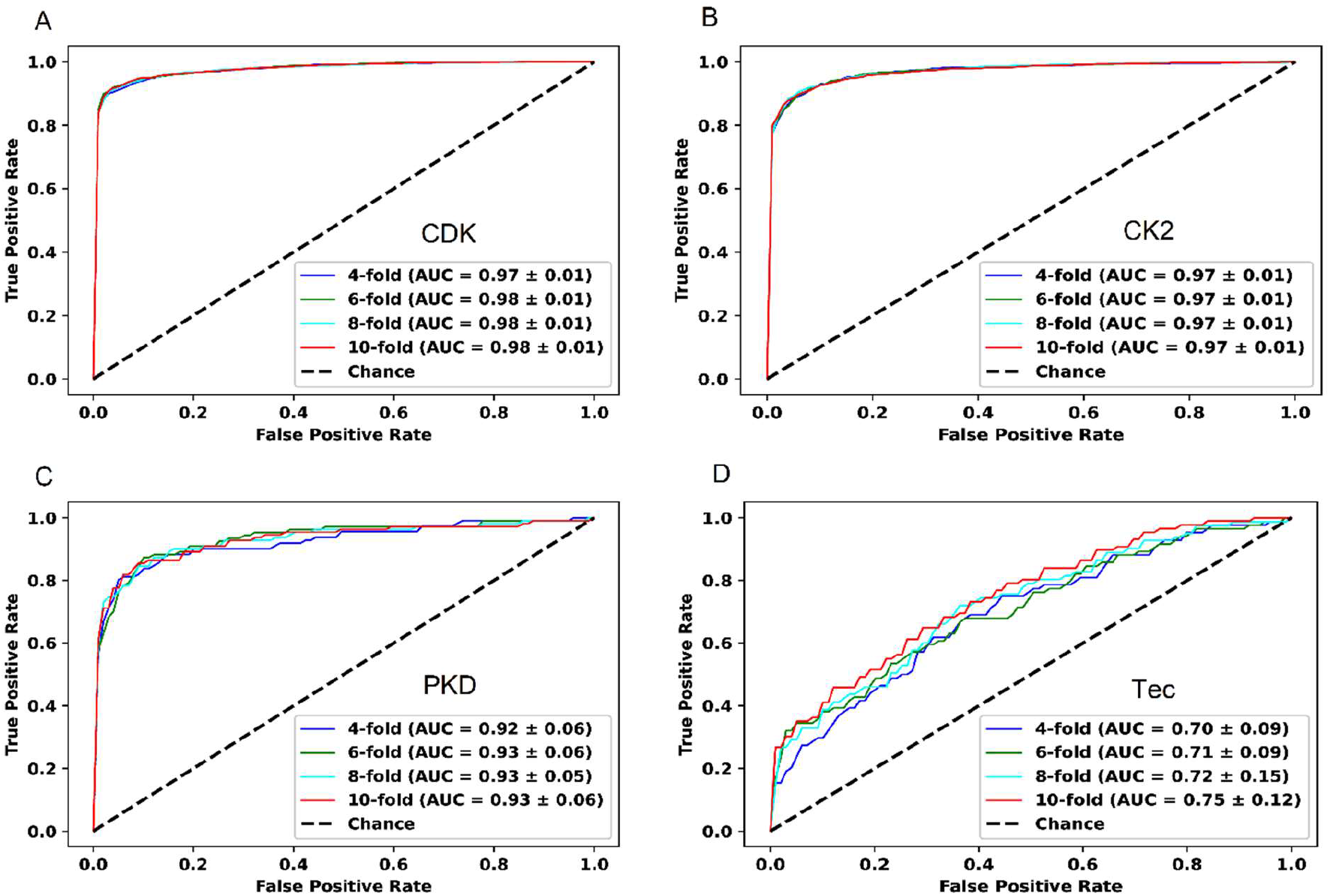
Performance comparisons between KinasePhos 3.0 at different levels of cross-validation. The results presented are from the (**A**) CDK, (**B**) CK2, (**C**) PKD, and (**D**) Tec family models, with 4-, 6-, 8-, and 10-fold cross-validations.

Human Beclin-1 (UniProt ID: Q14457) has been used as a test protein in GPS 5.0 to predict kinase-specific phosphorylation sites. For comparison, we used it to investigate the predictions made by KinasePhos 3.0. AGC family models were selected as representative models. GPS 5.0 predicted 38, 49, and 56 phosphorylation sites with high, medium, and low thresholds, respectively, while KinasePhos 3.0 obtained 33 phosphorylation sites. It should be noted that all these 33 phosphorylation sites lie within the 56 phosphorylation sites predicted by GPS 5.0 with a low threshold. Of these 33 phosphorylation sites, 30 belong to the 49 phosphorylation sites predicted by GPS 5.0, with a medium threshold, and 25 of these 33 phosphorylation sites fall among the 38 phosphorylation sites predicted by GPS 5.0 with a high threshold. Therefore, the prediction results from KinasePhos 3.0 are reasonably consistent with GPS 5.0.

### Results of feature interpretation with SHAP

We used mitogen-activated protein kinase 1 (*MAPK1*, UniProt ID P28482), of *Homo sapiens* to test the Akt family prediction model. *MAPK1* is a serine/threonine kinase that plays an essential role in the MAPK signal transduction pathway. Notably, residues 29, 185, 187, 190, 246, 248, and 284, in *MAPK1* can be phosphorylated [29]. To further investigate the importance of feature groups to amino acid characteristics of these 15-mer sequences, iceLogo [49] (https://iomics.ugent.be/icelogoserver/create), which is a web-based service capable of visualizing conserved patterns in protein and nucleotide sequences with probability theory, was used to obtain sequence logos to compare the difference between positive and negative data belonging to the same clusters.

**Figure 6**A and 6C represent the impact of feature groups on model output, while Figure 6B shows the iceLgo of the positive phosphorylation sites of the Akt family in contrast to the negative data. Figure 6D shows a heat map of the mean absolute SHAP values to show the impact of the features on the model output magnitude. It can be observed that the third position (pos-3) and fifth position (pos-5) before the phosphorylated sites have a relatively significant negative impact on the model prediction results. The results computed from SHAP are consistent with the iceLogo sequence and also with the position weight values computed for the Akt family at positions −5 and −3 in GPS 5.0, which were 0.85 and 1.00, respectively [22].

**Figure 6.**
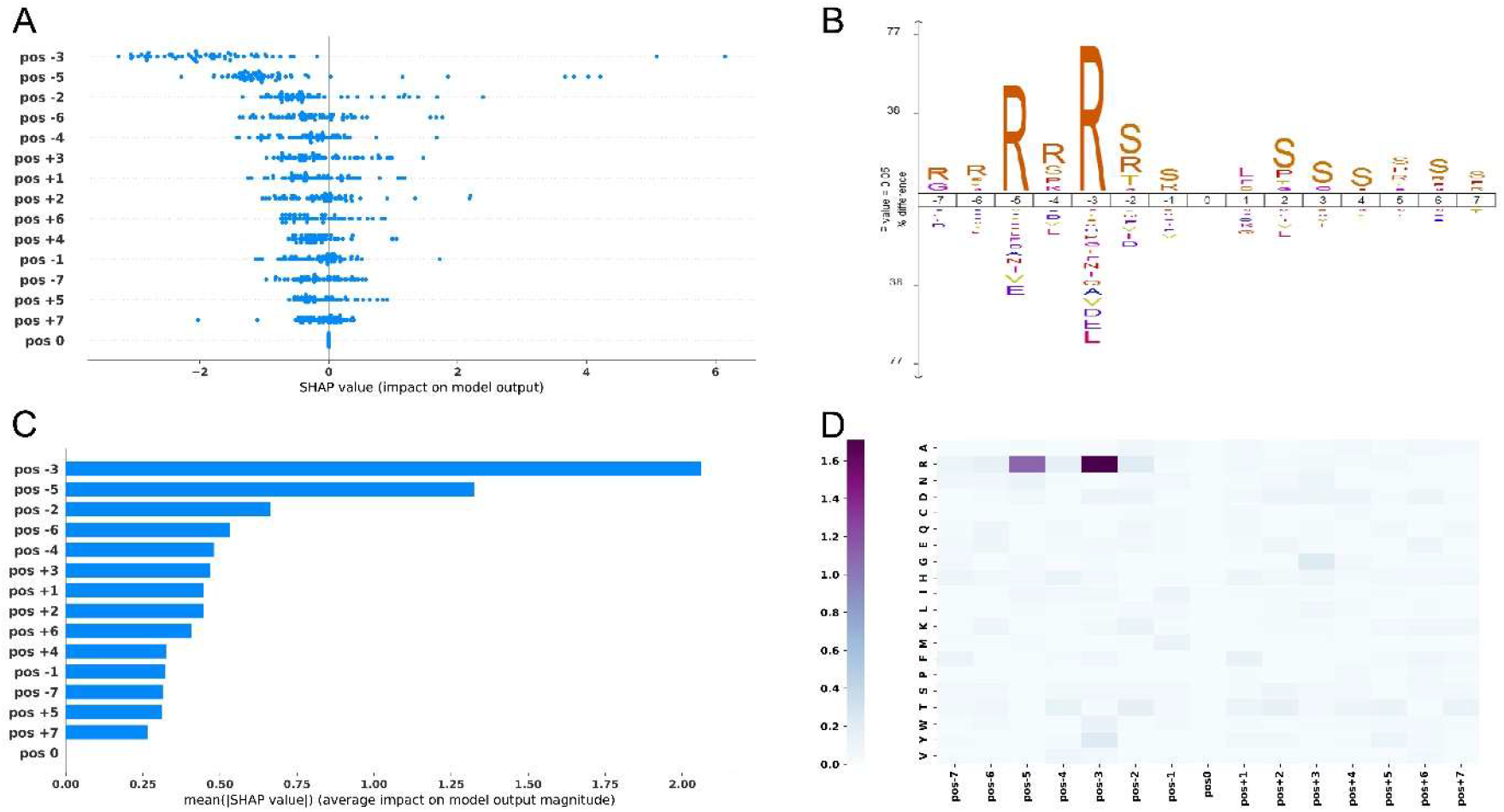
Feature explained by SHAP values. **A.** SHAP values showing the impacts of feature groups on model output. **B**. iceLogo of Akt family positive phosphorylation sites contrasted with its negative sites. **C**. Mean absolute SHAP values demonstrating the average impact of feature groups on model output magnitude. **D**. Heat map of mean absolute SHAP values. (**A**), (**C**) and (**D**) are derived from using mitogen-activated protein kinase 1 protein to test the Akt family prediction model.

### Web interface and downloadable prediction tool of the KinasePhos 3.0

The KinasePhos3.0 prediction service can be accessed via a web interface and a standalone prediction tool, the usages of which are presented in this section. *MAPK1* (UniProt ID: P28482) and human *Beclin-1* (UniProt ID: Q14457) were selected to illustrate the prediction of kinase-specific phosphorylation sites.

### Web interface

The core parts of the web interface allowing users to upload data, choose kinase models, and view predictions are presented in Figure S2A and S2B. The navigation bar “WEB SERVICES” (1 in Figure S2A) allows users to choose models of a specific type for the seven cluster types: group clusters considering S/T and Y sites; family clusters considering S/T and Y sites, S/T sites, and Y sites; and kinase-specific clusters considering S/T and Y sites, S/T sites, and Y sites. After choosing the model type, users can upload their FASTA format sequence data by clicking the “Choose File” button (2 in Figure S2A). Alternatively, users can enter protein UniProt IDs in the box (2 in Figure S2A) separated by a semicolon. Users can then choose models by ticking checkboxes (3 in Figure S2A). It should be noted that these models are sub-clustered into ten kinase groups: “AGC,” “Atypical,” “CAMK,” “CK1,” “CMGC,” “Other,” “PKL,” “STE,” “TK” and “TKL,” so users can click a specific kinase group first and then check the models belonging to that group. Subsequently, the users can click the “START KINASEPHOS” button (4 in Figure S2A) to run the prediction, following which, the result page, shown in Figure S2B, will finally appear.

As shown in section 1 of Figure S2B, users can download predictions in TXT format by clicking the download button. Section 2 summarizes the proteins uploaded or entered by users, along with the numbers of predicted phosphorylation sites for each protein and the models chosen by users. If a UniProt ID entered by users does not match any IDs in the UniProt database, it will be ignored. Section 3 lists the predicted sites. Similar to GPS 5.0, a column called “Source” is used to indicate whether the phosphorylation site has been experimentally verified (Exp.) or merely predicted (Pred.). To view the details for a specific protein, users can click the UniProt ID in column “Input ID” of section 2, and the page shown in **Figure 7** will appear.

**Figure 7.**
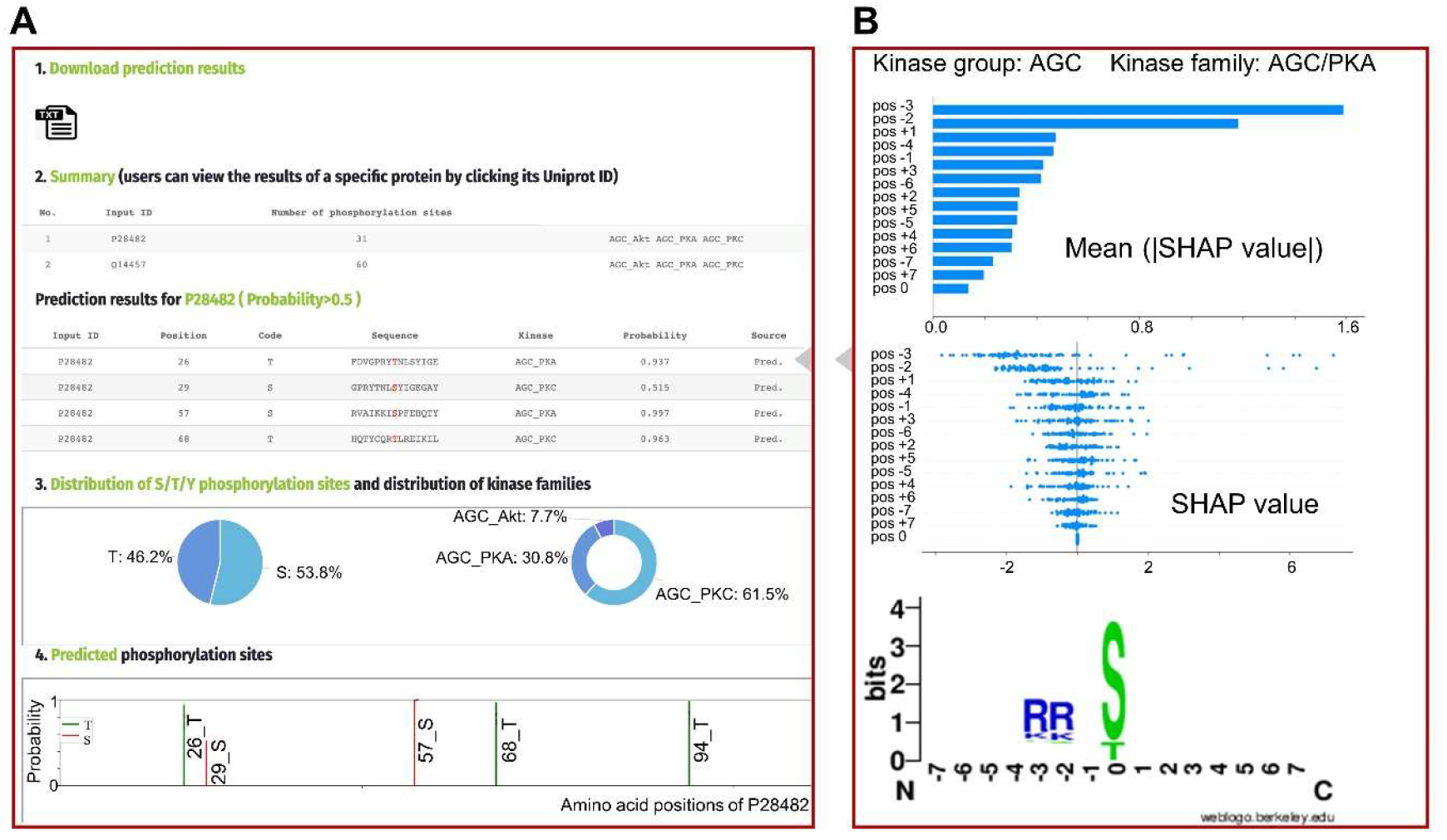
The web interface showing the results related to a specific protein. **A.** Detailed predictions for a specific protein, with predicted phosphorylation sites listed and depicted. **B.** Shapley additive explanations (SHAP) showing the impacts of feature groups on model output and sequence logo of the corresponding model.

Section 2 in Figure 7A lists the predicted sites for a particular protein (P28482 was clicked in this example). When users mouse over the rows of this table, the window shown by Figure 7B will be displayed on the right-hand side, showing the impact of feature groups on model output, (Figure 6A and C), and the sequence logo of the corresponding model. The distribution of S/T/Y phosphorylation sites and the distribution of the models are presented in the pie charts in Section 3. To provide a more intuitive view of the predicted phosphorylation sites, Section 4 displays predicted sites in a figure with probabilities, with S, T, and Y sites labeled in different colors. If users want to switch to predictions for another protein, they can click the protein’s ID in Section 2.

### Downloadable prediction tool

Considering the availability of large-scale phosphoproteomic data, a downloadable prediction tool (as shown in Figure S3) to predict all S/T and Y phosphorylation sites at kinase group, kinase family, and individual kinase levels is also provided at https://awi.cuhk.edu.cn/KinasePhos/download.html and https://github.com/tom-209/KinasePhos-3.0-executable-file. After downloading and starting KinasePhos3.exe, the “Browse” button is used to upload the data file, which should be a text file in FASTA format, as shown in the “Example Input.txt” file that is downloaded along with the tool. Users can then choose prediction models to test their data using checkboxes. If the “Kinase groups” are checked, all models at the group level will be executed. In addition, users can also choose group models separately by ticking the “AGC,” “Atypical,” “CAMK,” “CK1,” “CMGC,” “Other,” “PKL,” “STE,” “TK” or “TKL” checkboxes based on their requirements. Similarly, users can test their data using all models at the family and individual kinase levels by checking the “Kinase families” and the “Kinases” checkboxes, respectively. Additionally, users can choose specific family model or kinase model by clicking the corresponding checkboxes. It should be noted that these models at the kinase family level and individual kinase level are grouped into ten scroll areas corresponding to the ten kinase groups, while the models at the individual kinase level are further classified into human and other organisms for the convenience of testing data from humans and other species. With the models checked, users then click “Run and save” to run the prediction tool and save prediction results. It will take some time if users select many models and submit large-scale data before it produces a window that allows users to specify a location to save results as a CSV file. This downloadable prediction tool is recommended for users who want to test large-scale data using our predictive models.

## Discussion

Although advances in mass spectrometry and enrichment methods have led to a massive increase in high-throughput phosphoproteomic data, it is still difficult to determine the number of phosphorylation sites that can exist in a eukaryotic proteome [50] Vlastaridis et al. (2017) estimated that there are 230,000, 156,000, and 40,000 phosphorylation sites in human, mouse, and yeast, respectively [50]. However, as noted above, we only identified 41,421 experimentally verified, kinase-specific phosphorylation sites from 135 organisms, even with data that are already more comprehensive than those included in previous tools. The numbers of experimentally verified, kinase-specific phosphorylation sites in human, mouse, and yeast identified in this study were 19,123, 4,618, and 332, respectively. Therefore, for most phosphorylation sites, the kinases that phosphorylate them are yet to be identified. Computational methods are viable solutions for kinase-specific phosphorylation prediction, as empirical methods are more time-consuming and expensive. Kinase-specific phosphorylation sites in the kinase family and individual kinase levels are divided into S/T and Y, S/T, and Y site clusters, a total of 771 clusters, with a prediction model created for each.

The performance of KinasePhos 3.0 is competitive with other existing kinase-specific phosphorylation site prediction tools, such as GPS 5.0 and Scansite 4.0. It should be highlighted that the kinase-specific phosphorylation sites employed to develop KinasePhos 3.0 are more comprehensive than those employed with the existing tools, which is illustrated by the numbers of sites presented. In addition to collecting data from other existing tools, we text-mined experimentally verified kinase-specific phosphorylation sites from the UniProt database. Sample size is one of the most important parameters influencing model performance when developing machine learning-based classification models. We only used clusters with at least 15 experimentally verified phosphorylation sites when building models. This ensured that our sample size was comparable to those of some tools. For example, KinasePhos 2.0 used clusters with at least ten experimentally verified phosphorylation sites. In GPS 5.0, clusters with no less than three sites were considered, with 10-fold cross-validation and leave-one-out validation methods tested separately to evaluate the predictors’ performance with 245 kinase categories with no less than 30 sites and 372 kinase categories with 3 to 30 sites, respectively.

KinasePhos 3.0 also offers SHAP feature importance when performing prediction tasks. Since features were grouped based on their positions in the peptides containing 15 amino acids, that is, features related to a specific position were regarded as a feature group, feature group importance provides a more intuitive understanding of the implications of the surrounding residues on the phosphorylation of each peptide. The importance of the SHAP feature group is consistent with the sequence logo characteristics obtained from iceLogo, as illustrated above. The feature group importance is also consistent with the position weight computed using GPS. Instead of simply providing a prediction of whether a given residue can be phosphorylated by a specific kinase group, kinase family, or kinase, the inclusion of feature interpretation in the prediction models provides more insights into the potential roles of surrounding residues in phosphorylation.

Our study has several limitations. First, although we have collected a more comprehensive, experimentally verified kinase-specific phosphorylation site database than those used in other studies in this field, small numbers of these sites cannot be used to develop predictive models at the family or individual kinase level, as the number of sites is less than 15, below the threshold for creating a model, owing to data availability. However, these sites are included in the supplementary material for readers who might be interested in them. Second, the transfer learning technique adopted by Deznabi I et al. [51] might be employed to predict phosphorylation sites for kinases with less than 15 known phosphorylation sites. Moreover, considering protein-protein interactions and structural characteristics of proteins might improve predictions for kinases with few known phosphorylation sites. Third, we did not investigate deep learning methods, some of which have been described in the literature, such as DeepPhos [52] and MusiteDeep [53], and have demonstrated effectiveness in predicting kinase-specific phosphorylation sites. Leveraging the power of deep learning, along with more features, will be a good strategy to explore in the future to further increase the prediction performance. Fourth, our models do not distinguish among organisms, although the majority of phosphorylation sites are from humans, mice, and rats. Tools that can separate species may better satisfy some users’ requirements.

## Conclusion

In conclusion, our updated KinasePhos 3.0 represents a significant improvement over versions 1.0 and 2.0. Notably, more comprehensive experimentally verified kinase-specific phosphorylation site data have been collected, and prediction models have been increased, with the potential to meet more specific requirements of the users. The prediction performance of this version is competitive with that of other existing tools, such as GPS 5.0. Importantly, we provide users with both web-based and downloadable tools, making it more user-friendly. In the future, KinasePhos 3.0 will be valuable for predicting unknown sites, judging if these sites can be phosphorylated by a specific kinase group, kinase family, or kinase, based on user requirements. These predictions will aid in empirical kinase and substrate characterization, reducing costs and saving time.

## Code availability

The source code is available on Github: https://github.com/tom-209/KinasePhos-3.0-executable-file.

## CRediT author statement

**Renfei Ma**: Conceptualization; Data curation; Methodology; Formal analysis; Investigation; Software; Visualization; Writing - original draft. **Shangfu Li**: Conceptualization; Data curation; Formal analysis; Writing - original draft. **Wenshuo Li**: Software; Visualization. **Lantian Yao**: Software. **Hsien-Da Huang**: Project administration; Conceptualization; Supervision; Funding acquisition; Writing - review & editing. **Tzong-Yi Lee**: Project administration; Conceptualization; Supervision; Funding acquisition; Writing - review & editing.

## Competing interest

We declare no conflict of interests associated with this paper.

## Acknowledgments

The authors express their gratitude toward all database developers mentioned and quoted in this article for their important work and the data they shared. The author also would like to thank users for their comments and suggestions on the previous version of KinasePhos.

This work was supported by the National Natural Science Foundation of China [32070659], the Science, Technology and Innovation Commission of Shenzhen Municipality [JCYJ20200109150003938], Guangdong Province Basic and Applied Basic Research Fund [2021A1515012447], and Ganghong Young Scholar Development Fund [2021E007]. The authors sincerely appreciate the Warshel Institute for Computational Biology, the Chinese University of Hong Kong (Shenzhen) for financially supporting this research.

## Supplementary material

**Figure S1 The evolutionary tree for all the collected kinases**

**A.** The obtained evolutionary tree showed that homologous proteins or proteins with consistent domains clustered tightly in smaller branches. **B.** The kinome tree composed of several major groups. **C.** Analysis of the kinases of each group separately showed that the kinases in the same group contained similar domains.

**Figure S2 The web interface of KinasePhos 3.0**

**A.** The web interface for users to select model types, upload or enter their data and choose prediction models. **B.** The overview of predicted results.

**Figure S3 The downloadable prediction tool**

This is the standalone prediction tool of the KinasePhos3.0, which can predict S/T and Y phosphorylation sites at kinase group, kinase family, and in individual kinase levels.

**Table S1 The experimentally identified kinase-specific phosphorylation sites used in KinasePhos3.0**

**Table S2 The performance of the 771 models**

## Notes

### Competing Interest Statement

The authors have declared no competing interest.

## References

[1] Miller CJ, Turk BE. Homing in: mechanisms of substrate targeting by protein kinases. Trends Biochem Sci 2018;43:380–94.

[2] Delanghe T, Dondelinger Y, Bertrand MJ. RIPK1 kinase-dependent death: A symphony of phosphorylation events. Trends Cell Biol 2020;30:189–200.

[3] Taddei ML, Pardella E, Pranzini E, Raugei G, Paoli P. Role of tyrosine phosphorylation in modulating cancer cell metabolism. Biochim Biophys Acta Rev Cancer 2020;1874:188442.

[4] Kotrasová V, Keresztesová B, Ondrovičová G, Bauer JA, Havalová H, Pevala V, et al. Mitochondrial kinases and the role of mitochondrial protein phosphorylation in health and disease. Life 2021;11:82.

[5] Ge R, Shan W. Bacterial phosphoproteomic analysis reveals the correlation between protein phosphorylation and bacterial pathogenicity. Genomics Proteomics Bioinformatics, 2011;9:119–27.

[6] Jiang Y, Cong X, Jiang S, Dong Y, Zhao L, Zang Y, Tan M, Li J. Phosphoproteomics reveals AMPK substrate network in response to DNA damage and histone acetylation. Genomics Proteomics Bioinformatics 2021.

[7] Ji F, Zhou M, Zhu H, Jiang Z, Li Q, Ouyang X, Lv Y, Zhang S, Wu T, Li L. Integrative proteomic analysis of posttranslational modification in the inflammatory response. Genomics Proteomics Bioinformatics 2021.

[8] Ochoa D, Bradley D, Beltrao P. Evolution, dynamics and dysregulation of kinase signalling. Curr Opin Struct Biol 2018;48:133–40.

[9] Chen Y, Shao X, Cao J, Zhu H, Yang B, He Q, et al. Phosphorylation regulates cullin-based ubiquitination in tumorigenesis. Acta Pharm Sin B 2020; 11: 309–21.

[10] Gong T, Jiang W, Zhou R. Control of inflammasome activation by phosphorylation. Trends Biochem Sci 2018;43:685–99.

[11] Pandey V, Zhu T, Ma L, Lobie PE, et al. Bad phosphorylation as a target of inhibition in oncology. Cancer Lett 2018;415:177–86.

[12] Veerman GM, Hussaarts KG, Jansman FG, Koolen SW, van Leeuwen RW, Mathijssen RH. Clinical im-plications of food-drug interactions with small-molecule kinase inhibitors. Lancet Oncol 2020;21:e265–79.

[13] Abdeldayem A, Raouf YS, Constantinescu SN, Moriggl R, Gunning PT. Advances in covalent kinase inhibitors. Chem Soc Rev 2020;49:2617–87.

[14] Baltussen LL, Rosianu F, Ultanir SK. Kinases in synaptic development and neurological diseases. Prog Neuropsychopharmacol Biol Psychiatry 2018;84:343–52.

[15] Yang L, Zheng L, Chng WJ, Ding JL. Comprehensive analysis of ERK1/2 substrates for potential combination immunotherapies. Trends Pharmacol Sci 2019;40:897–910.

[16] Low TY, Mohtar MA, Lee PY, Omar N, Zhou H, Ye M. Widening the bottleneck of phosphoproteomics: Evolving strategies for phosphopeptide enrichment. Mass Spectrom Rev 2021;40:309–33.

[17] Hirst NL, Nebel JC, Lawton SP, Walker AJ. Deep phosphoproteome analysis of Schistosoma mansoni leads development of a kinomic array that highlights sex-biased differences in adult worm protein phosphorylation. PLoS Negl Trop Dis 2020;14:e0008115.

[18] Xue B, Jordan B, Rizvi S, Naegle KM. KinPred: A unified and sustainable approach for harnessing proteome-level human kinase-substrate predictions. PLoS Comput Biol 2021;17:e1008681.

[19] Song J, Wang H, Wang J, Leier A, Marquez-Lago T, Yang B, et al. PhosphoPredict: A bioinformatics tool for prediction of human kinase-specific phosphorylation substrates and sites by integrating heterogeneous feature selection. Sci Rep 2017;7:1—19.

[20] Blom N, Sicheritz-Pontén T, Gupta R, Gammeltoft S, Brunak S. Prediction of post-translational glycosylation and phosphorylation of proteins from the amino acid sequence. Proteomics 2004;4:1633—49.

[21] Li F, Li C, Marquez-Lago T, Leier A, Akutsu T, Purcell AW, Ian Smith A, Lithgow T, Daly, RJ, Song J and Chou KC. Quokka: a comprehensive tool for rapid and accurate prediction of kinase family-specific phosphorylation sites in the human proteome. Bioinformatics, 2018:34:4223–31.

[22] Gao J, Thelen JJ, Dunker AK, Xu D. Musite, a tool for global prediction of general and kinase-specific phosphorylation sites. Mol Cell Proteomics 2010;9:2586—600.

[23] Wang C, Xu H, Lin S, Deng W, Zhou J, Zhang Y, et al. GPS 5.0: an update on the prediction of kinase-specific phosphorylation sites in proteins. Genomics Proteomics Bioinformatics 2020;18:72—80.

[24] Dang TH, Van Leemput K, Verschoren A, Laukens K. Prediction of kinase-specific phosphorylation sites using conditional random fields. Bioinformatics 2008;24:2857—64.

[25] Kim JH, Lee J, Oh B, Kimm K, Koh I. Prediction of phosphorylation sites using SVMs. Bioinformatics 2004;20:3179–84.

[26] Huang HD, Lee TY, Tzeng SW, Horng JT. KinasePhos: a web tool for identifying protein kinase-specific phosphorylation sites. Nucleic Acids Res 2005;33:W226—9.

[27] Wong YH, Lee TY, Liang HK, Huang CM, Wang TY, Yang YH, et al. KinasePhos 2.0: a web server for identifying protein kinase-specific phosphorylation sites based on sequences and coupling patterns. Nucleic Acids Res 2007;35:W588—94.

[28] Hornbeck PV, Kornhauser JM, Latham V, Murray B, Nandhikonda V, Nord A, et al. 15 years of Phospho-SitePlus®: integrating post-translationally modified sites, disease variants and isoforms. Nucleic Acids Res 2019;47:D433—41.

[29] UniProt: The universal protein knowledgebase in 2021. Nucleic Acids Res 2021;49:D480—9.

[30] Dinkel H, Chica C, Via A, Gould CM, Jensen LJ, Gibson TJ, et al. Phospho. ELM: a database of phosphorylation sites—update 2011. Nucleic Acids Res 2010;39:D261–7.

[31] Manning G, Whyte DB, Martinez R, Hunter T, Sudarsanam S. The protein kinase complement of the human genome. Science 2002;298:1912–34.

[32] Rozewicki J, Li S, Amada KM, Standley DM, Katoh K. MAFFT-DASH: integrated protein sequence and structural alignment. Nucleic Acids Res 2019;47:W5—10.

[33] Price MN, Dehal PS, Arkin AP. FastTree 2—approximately maximum-likelihood trees for large alignments. PLoS One 2010;5:e9490.

[34] Stöver BC, Müller KF. TreeGraph 2: combining and visualizing evidence from different phylogenetic analyses. BMC Bioinformatics 2010;11:1–9.

[35] Mistry J, Chuguransky S, Williams L, Qureshi M, Salazar GA, Sonnhammer EL, et al. Pfam: The protein families database in 2021. Nucleic Acids Res 2021;49:D412–9.

[36] Letunic I, Khedkar S, Bork P. SMART: recent updates, new developments and status in 2020. Nucleic Acids Res 2021;49:D458–60.

[37] Letunic I, Bork P. Interactive Tree Of Life (iTOL): an online tool for phylogenetic tree display and annotation. Bioinformatics 2007;23:127–8.

[38] Xue Y, Ren J, Gao X, Jin C, Wen L, Yao X. GPS 2.0, a tool to predict kinase-specific phosphorylation sites in hierarchy. Mol Cell Proteomics 2008;7:1598–608.

[39] Jing X, Dong Q, Hong D, Lu R. Amino acid encoding methods for protein sequences: A comprehensive review and assessment. IEEE/ACM Trans Comput Biol Bioinform 2019;17:1918–31.

[40] Chen T, Guestrin C. Xgboost: A scalable tree boosting system. Proc ACM SIGKDD Int Conf Knowl Discov Data Min 2016; 22^nd^ (August): 785–94.

[41] Lundberg SM, Lee SI. A unified approach to interpreting model predictions. Adv Neural Inf Process Syst 2017;30:4765–74.

[42] Wang J, Gribskov M. IRESpy: an XGBoost model for prediction of internal ribosome entry sites. BMC Bioinformatics 2019;20:1–15.

[43] Bi Y, Xiang D, Ge Z, Li F, Jia C, Song J. An interpretable prediction model for identifying N7-methylguanosine sites based on XGBoost and SHAP. Mol Ther Nucleic Acids 2020;22:362–72.

[44] Lv Z, Cui F, Zou Q, Zhang L, Xu L. Anticancer peptides prediction with deep representation learning features. Brief Bioinform 2021.

[45] Henikoff S, Henikoff JG. Amino acid substitution matrices from protein blocks. Proc Natl Acad Sci USA 1992;89:10915–9.

[46] Huang Y, Niu B, Gao Y, Fu L, Li W. CD-HIT Suite: a web server for clustering and comparing biological sequences. Bioinformatics 2010;26:680–2.

[47] Kao HJ, Nguyen VN, Huang KY, Chang WC, Lee TY. SuccSite: incorporating amino acid composition and informative k-spaced amino acid pairs to identify protein succinylation sites. Genomics Proteomics Bioinformatics 2020; 18:208–19.

[48] Obenauer JC, Cantley LC, Yaffe MB. Scansite 2.0: Proteome-wide prediction of cell signaling interac-tions using short sequence motifs. Nucleic Acids Res 2003;31:3635–41.

[49] Colaert N, Helsens K, Martens L, Vandekerckhove J, Gevaert K. Improved visualization of protein consensus sequences by iceLogo. Nat Methods 2009;6:786–7.

[50] Vlastaridis P, Kyriakidou P, Chaliotis A, Van de Peer Y, Oliver SG, Amoutzias GD. Estimating the total number of phosphoproteins and phosphorylation sites in eukaryotic proteomes. Gigascience 2017;6:giw01

[51] Deznabi I, Arabaci B, Koyutürk M, Tastan O. DeepKinZero: zero-shot learning for predicting kinase-phosphosite associations involving understudied kinases. Bioinformatics 2020;36:3652–61.

[52] Luo F, Wang M, Liu Y, Zhao XM and Li A. DeepPhos: prediction of protein phosphorylation sites with deep learning. Bioinformatics 2019; 35:2766–73.

[53] Wang D, Zeng S, Xu C, Qiu W, Liang Y, Joshi T and Xu D. MusiteDeep: a deep-learning framework for general and kinase-specific phosphorylation site prediction. Bioinformatics 2017; 33:3909–16

