## Supplementary figures and images for "KinasePhos 3.0: Redesign and Expansion of the Prediction on Kinase-specific Phosphorylation Sites"

### Supplementary Figure S2

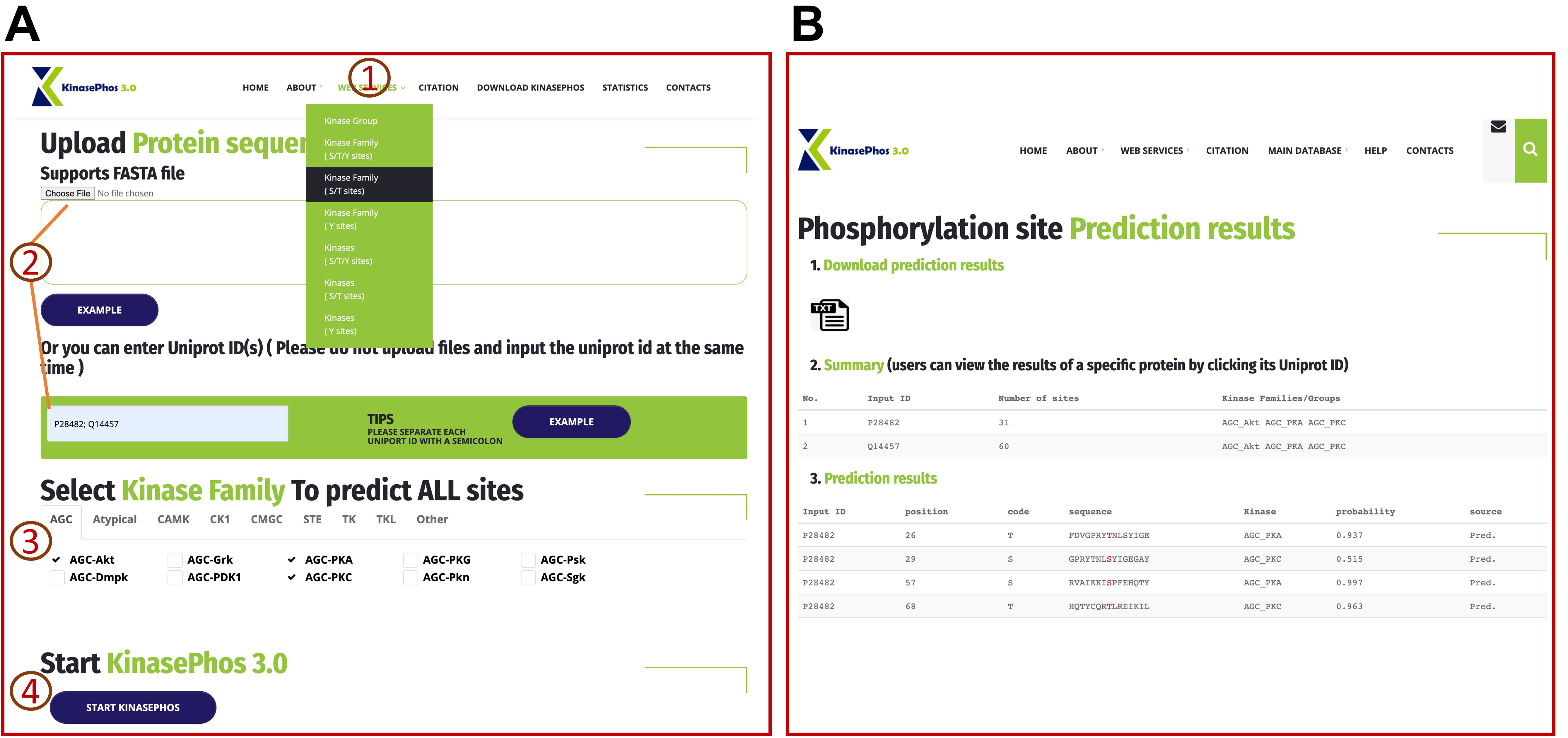

### Supplementary Figure S3

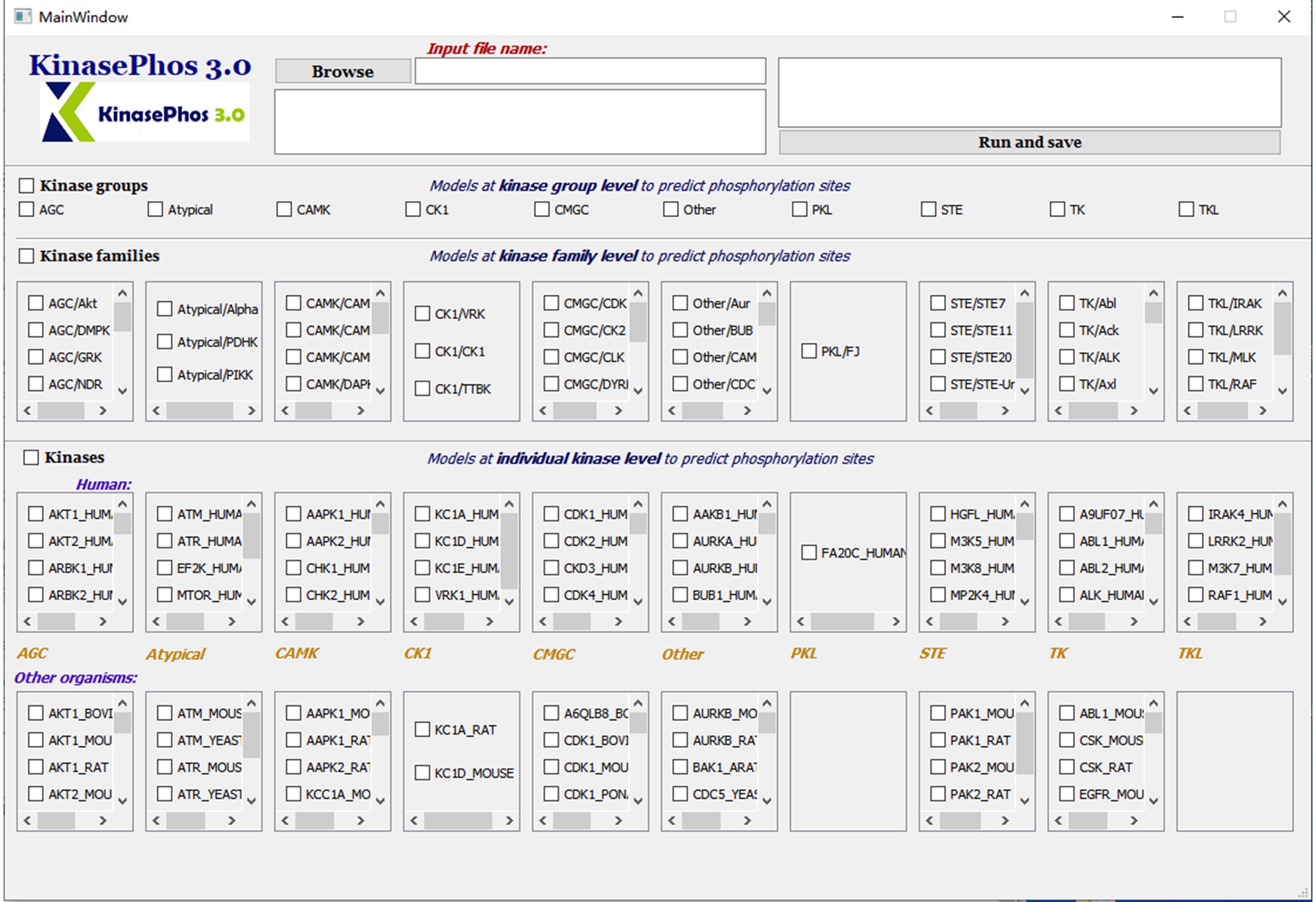
