## Supplementary Table S2 for "KinasePhos 3.0: Redesign and Expansion of the Prediction on Kinase-specific Phosphorylation Sites"

| **Clusters** | **No. of positive sites** | **Accuracy** | **Weighted f1 score** | **Weighted precision** | **Weighted recall** | **AUC** | **Logo** | **Mean \|SHAP value\|** | **SHAP value** |
| --- | --- | --- | --- | --- | --- | --- | --- | --- | --- |
| **AGC (STY)** | 4602 | 0.901 | 0.901 | 0.901 | 0.901 | 0.958 | 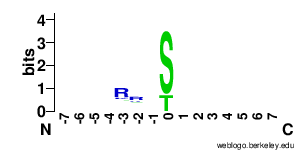 | 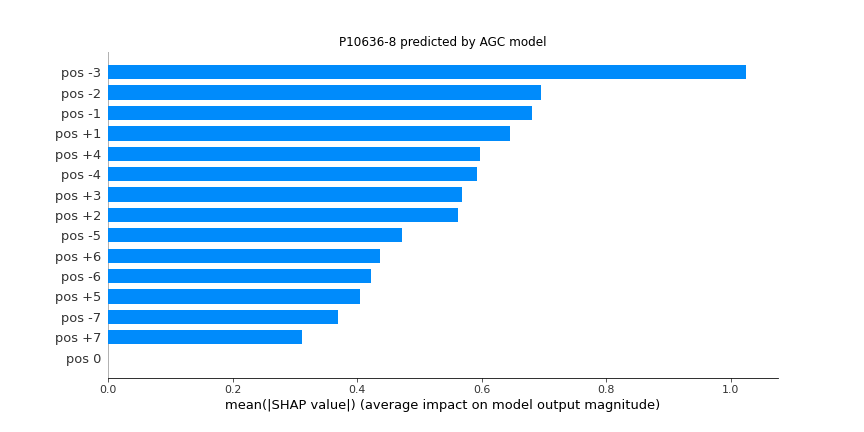 | 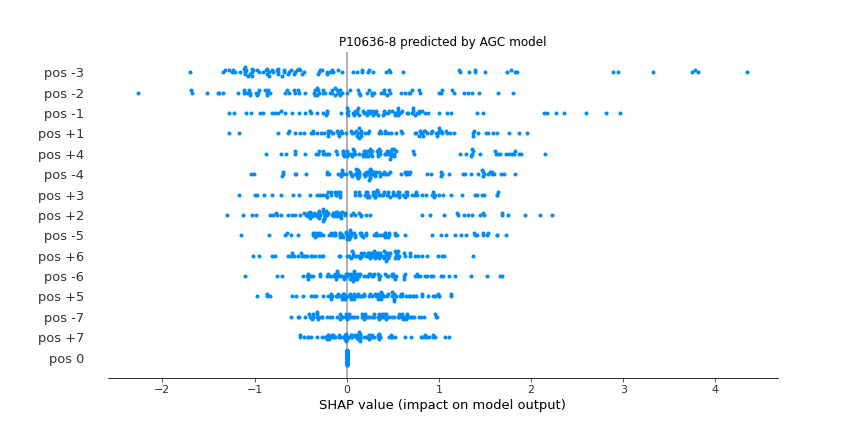 |
| **Atypical (STY)** | 1037 | 0.886 | 0.88 | 0.888 | 0.886 | 0.935 | 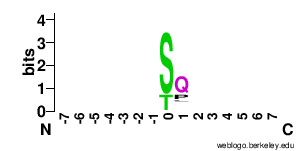 | 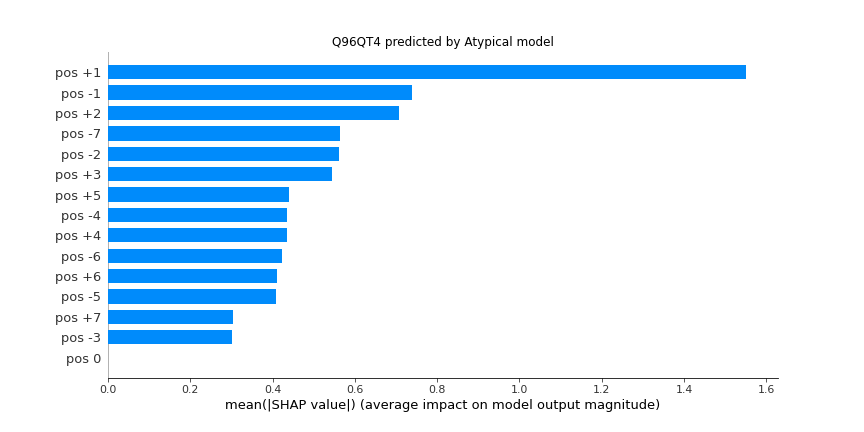 | 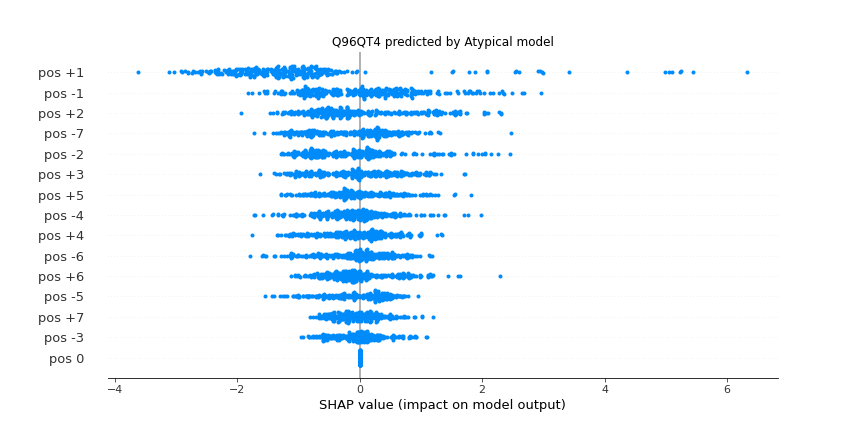 |
| **CAMK (STY)** | 1892 | 0.852 | 0.852 | 0.854 | 0.852 | 0.928 | 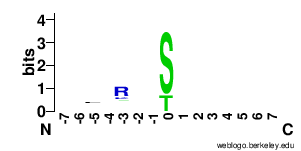 | 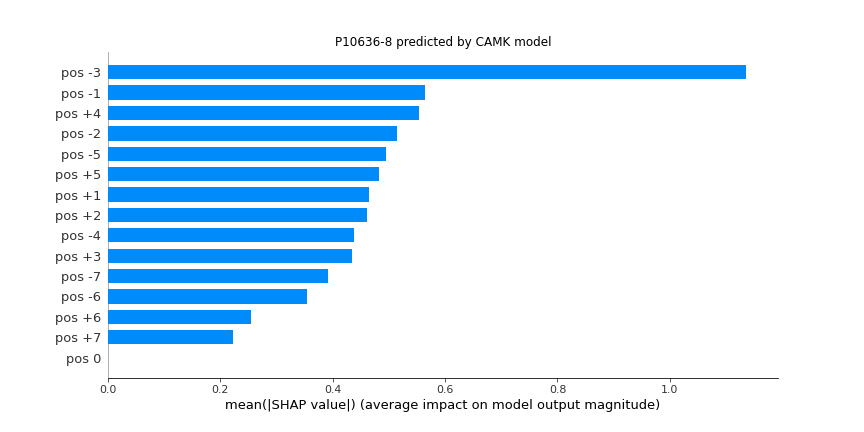 | 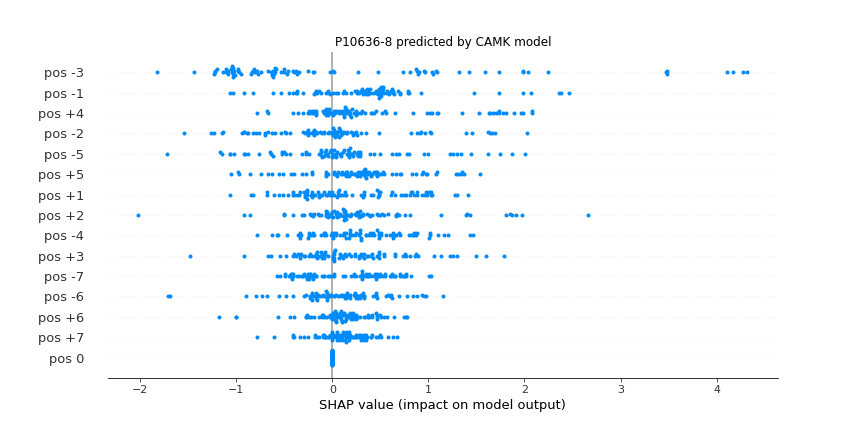 |
| **CK1 (STY)** | 508 | 0.857 | 0.849 | 0.854 | 0.857 | 0.888 | 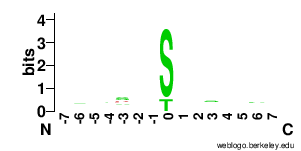 | 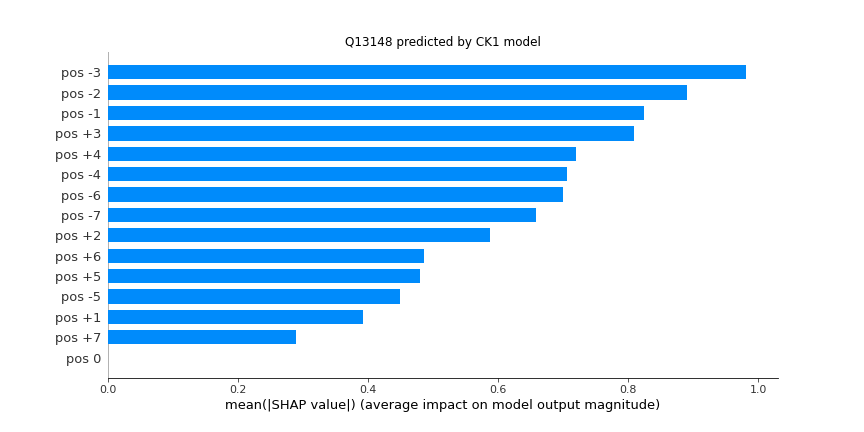 | 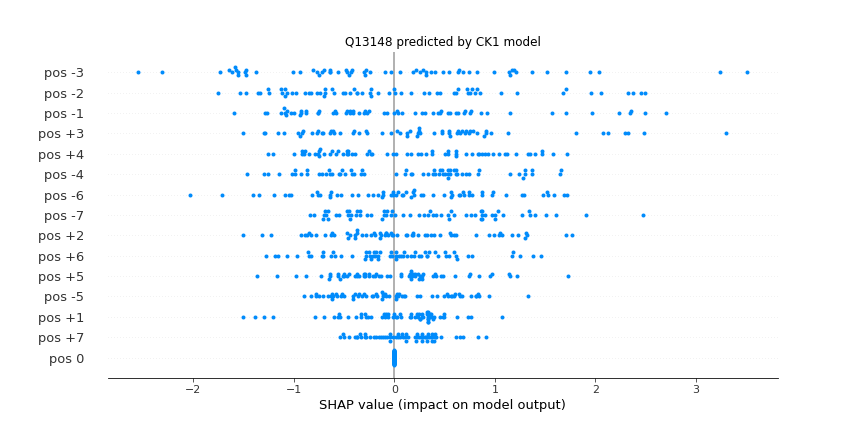 |
| **CMGC (STY)** | 5737 | 0.943 | 0.943 | 0.944 | 0.943 | 0.982 | 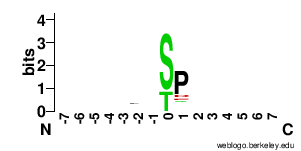 | 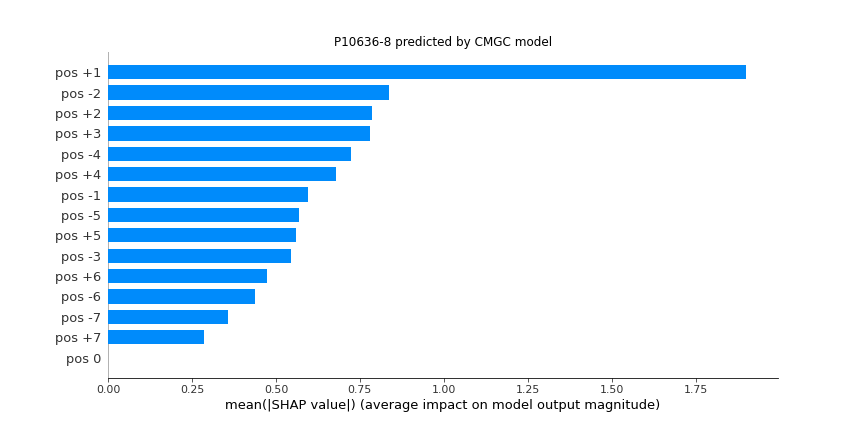 | 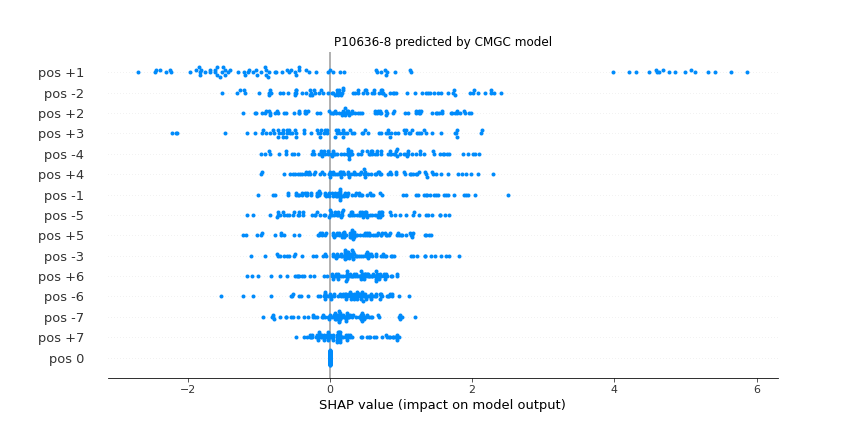 |
| **Other (STY)** | 2068 | 0.79 | 0.79 | 0.792 | 0.79 | 0.875 | 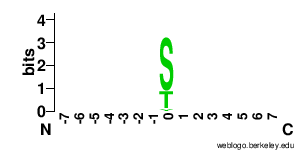 | 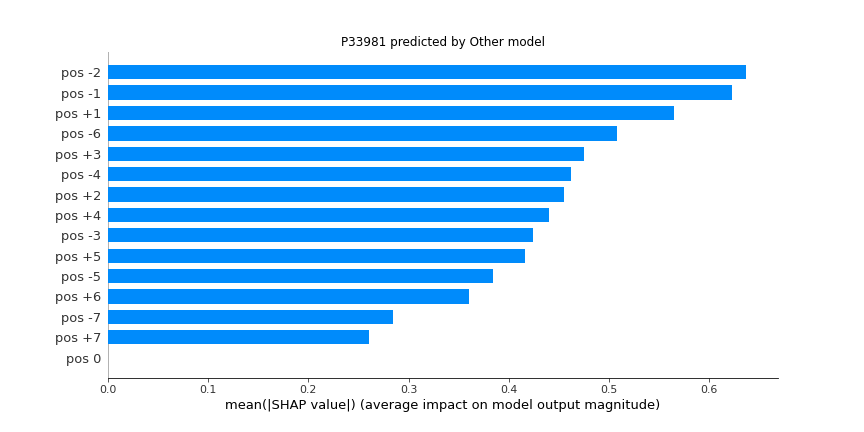 | 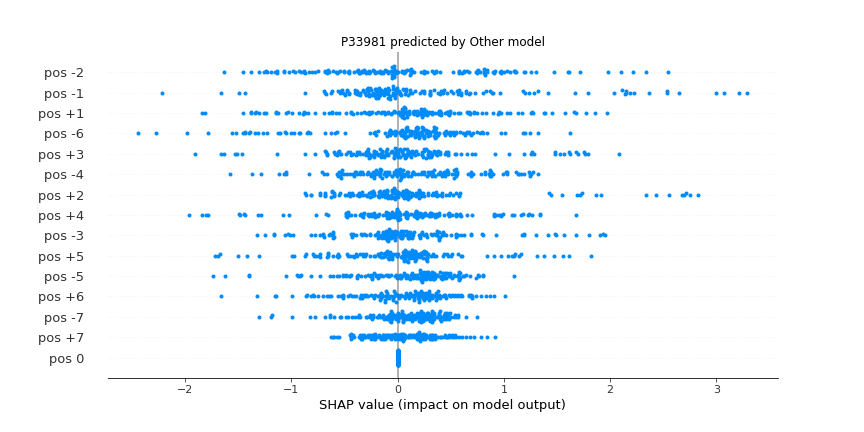 |
| **PKL (STY)** | 204 | 0.882 | 0.875 | 0.876 | 0.882 | 0.862 | 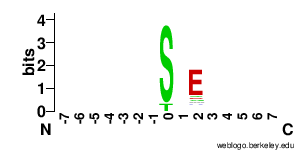 | 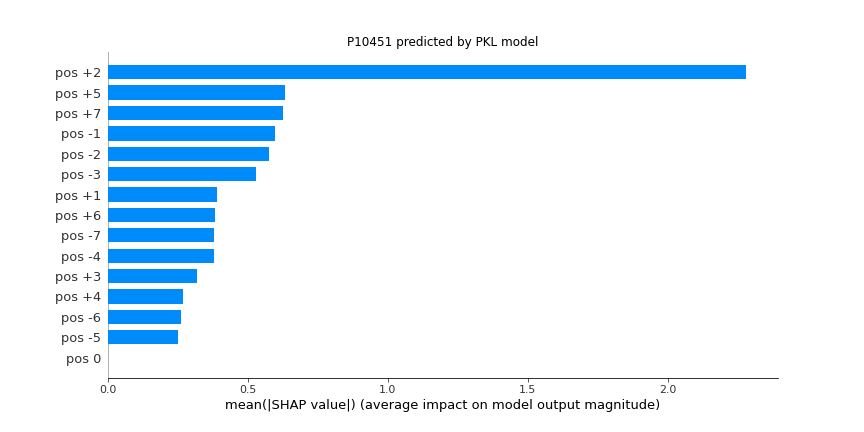 | 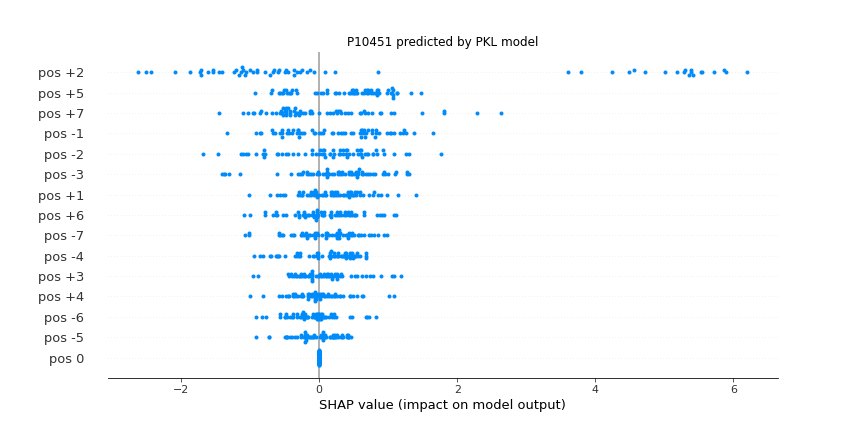 |
| **STE (STY)** | 625 | 0.837 | 0.826 | 0.833 | 0.837 | 0.851 | 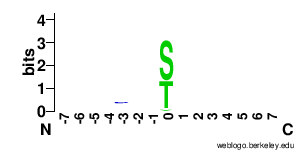 | 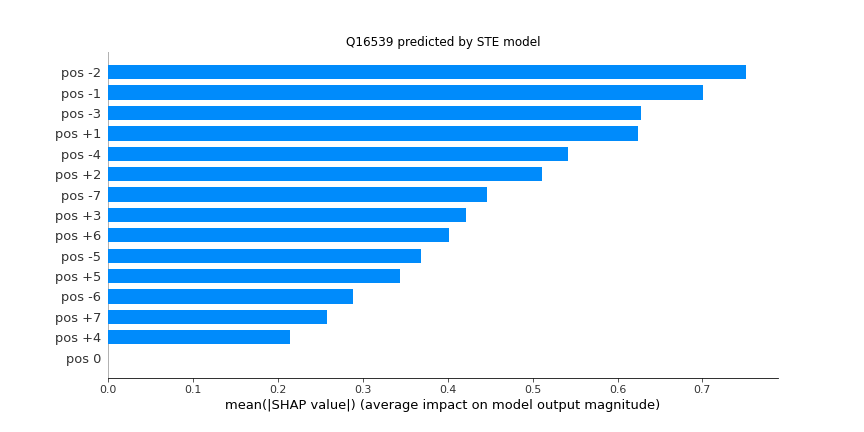 | 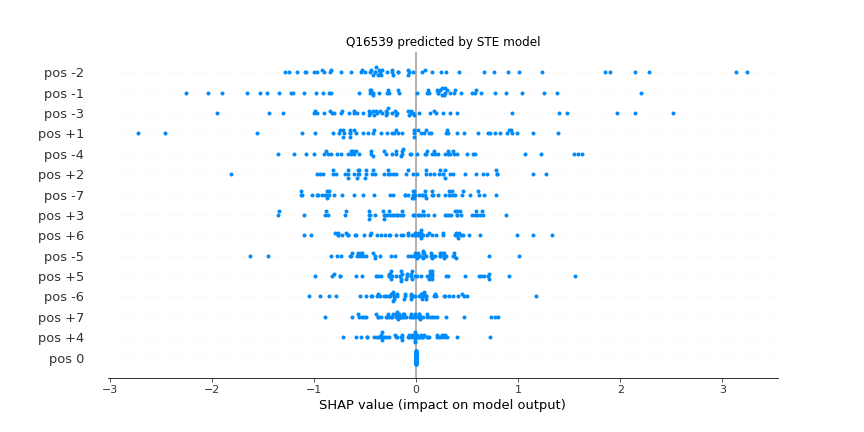 |
| **TK (STY)** | 2680 | 0.808 | 0.807 | 0.808 | 0.808 | 0.884 | 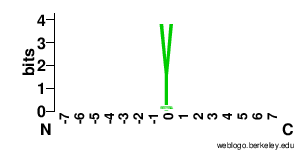 | 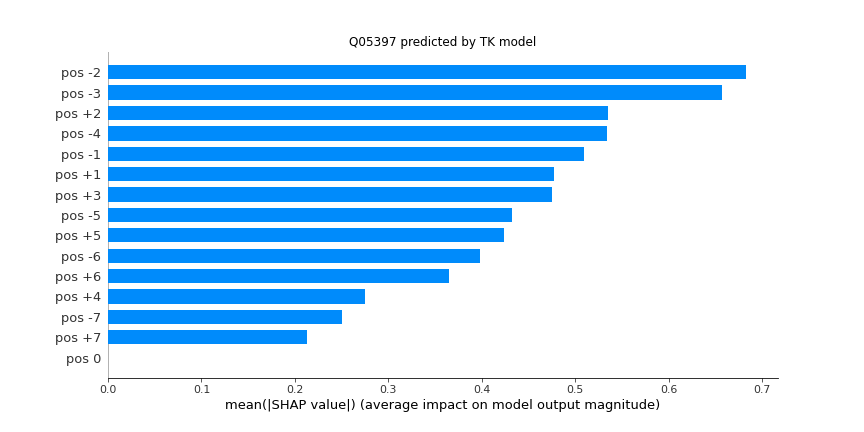 | 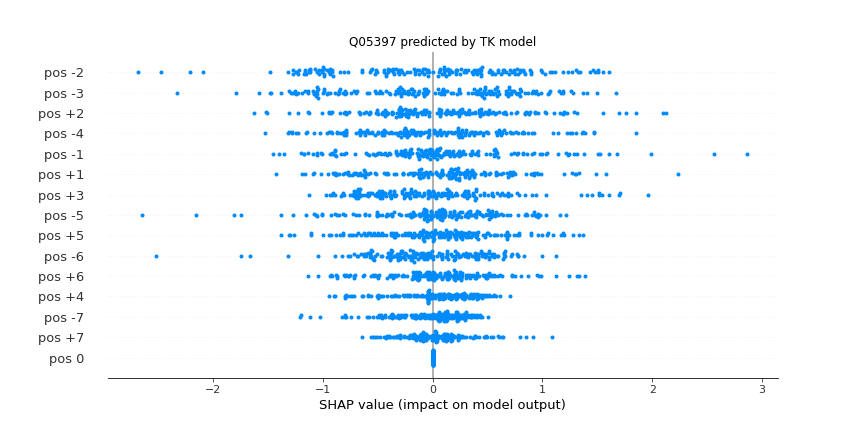 |
| **TKL (STY)** | 360 | 0.802 | 0.769 | 0.775 | 0.802 | 0.744 | 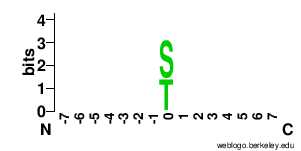 | 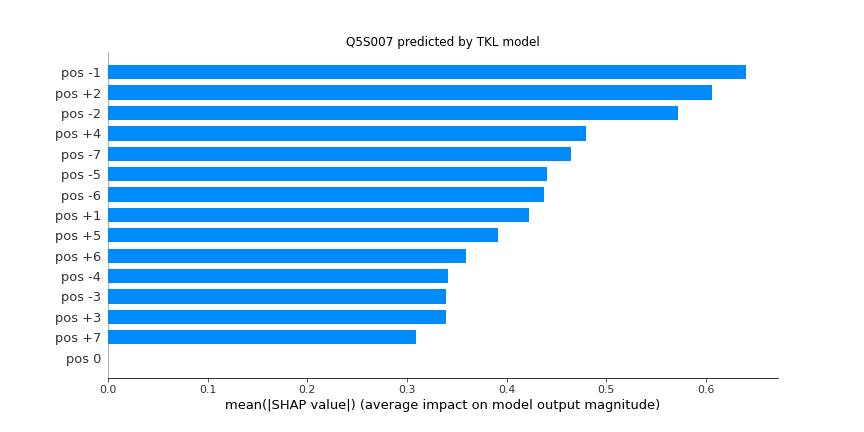 | 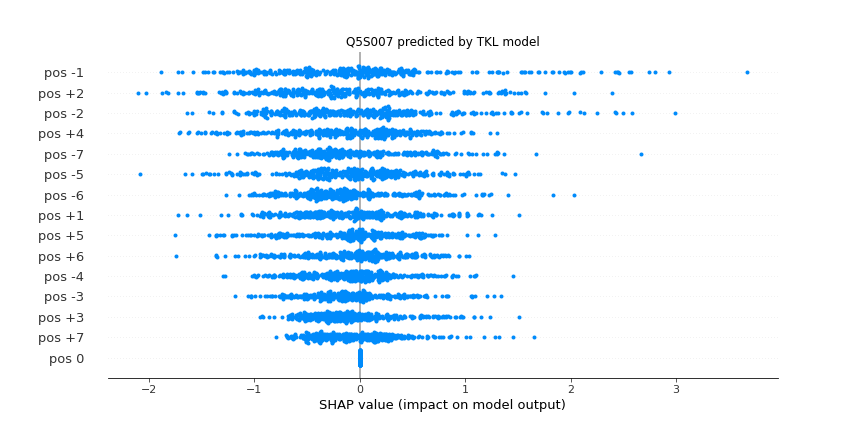 |
| **AGC_Akt (STY)** | 526 | 0.945 | 0.943 | 0.944 | 0.945 | 0.965 | 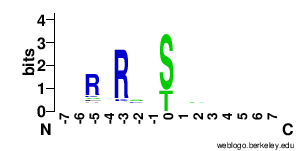 | 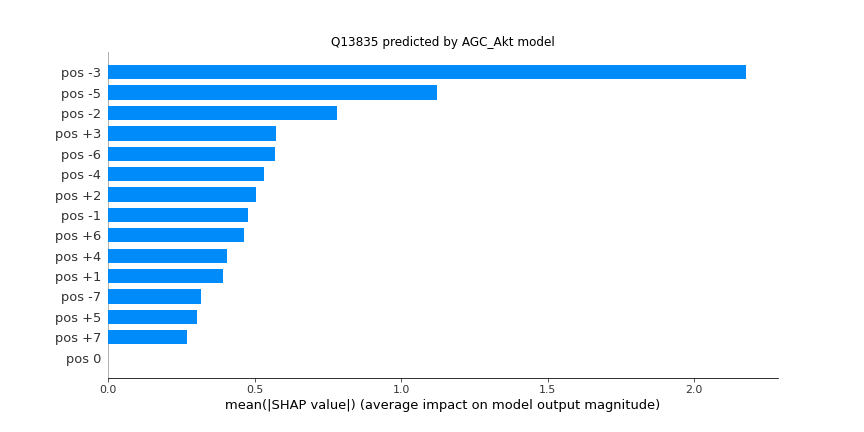 | 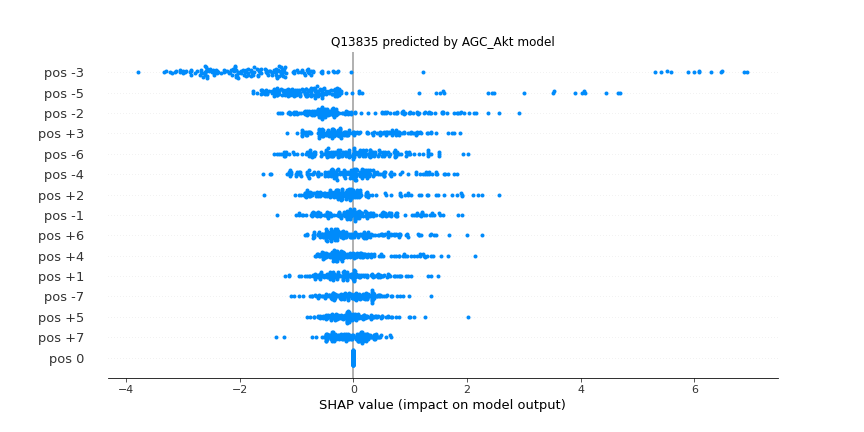 |
| **AGC_DMPK (STY)** | 154 | 0.899 | 0.889 | 0.895 | 0.899 | 0.87 | 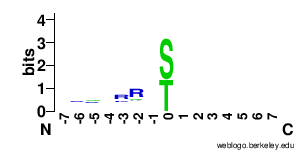 | 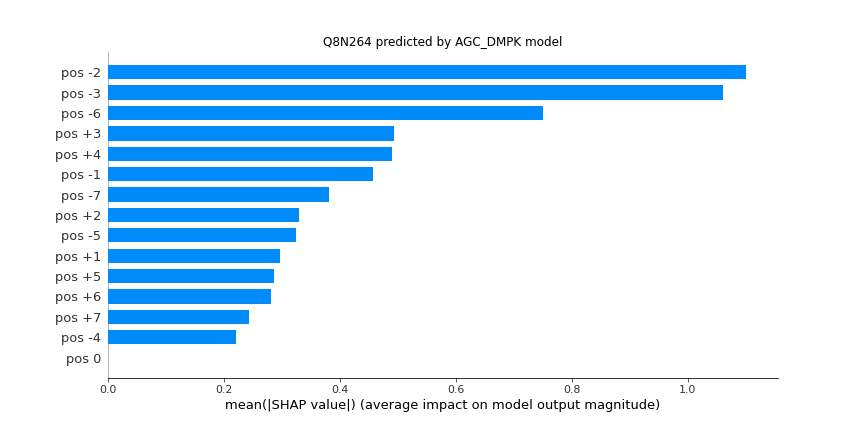 | 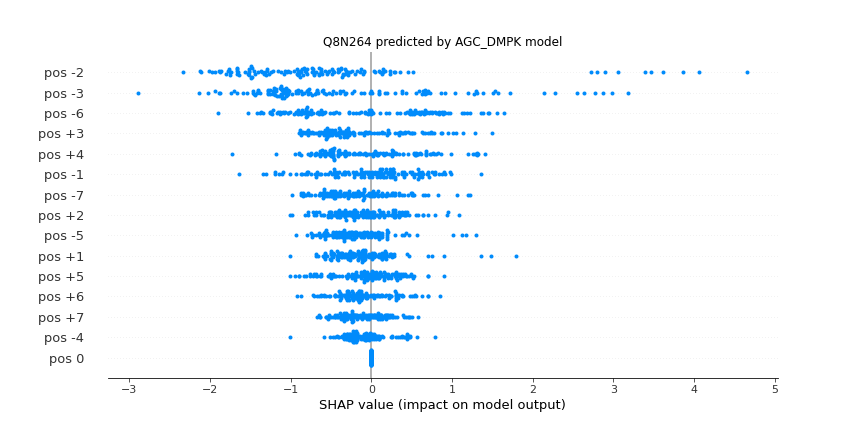 |
| **AGC_GRK (STY)** | 276 | 0.861 | 0.846 | 0.861 | 0.861 | 0.851 | 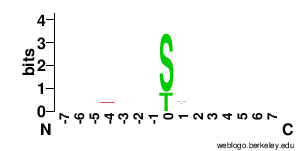 | 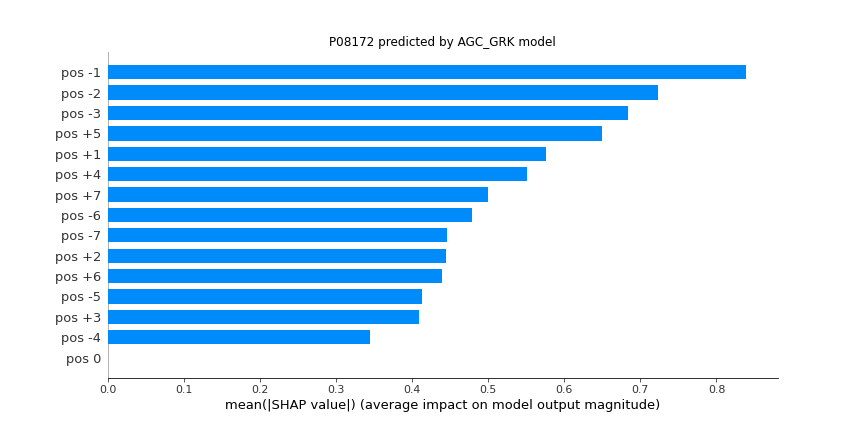 | 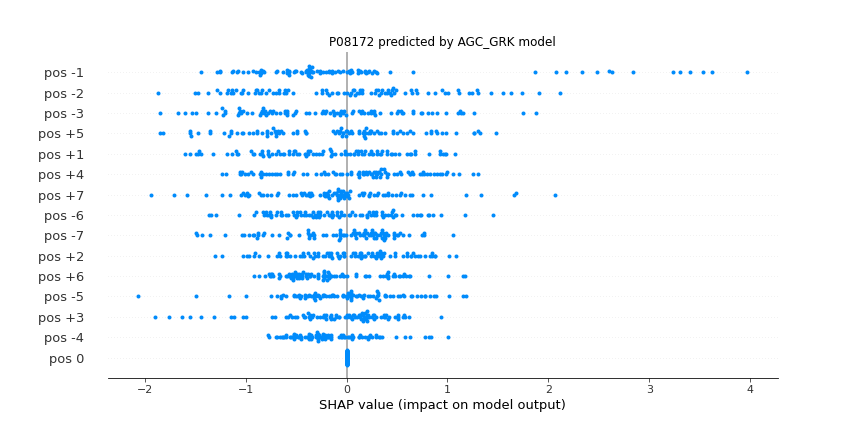 |
| **AGC_NDR (STY)** | 48 | 0.959 | 0.951 | 0.963 | 0.959 | 0.958 | 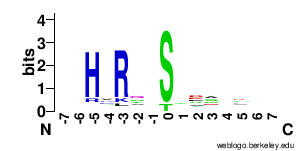 | 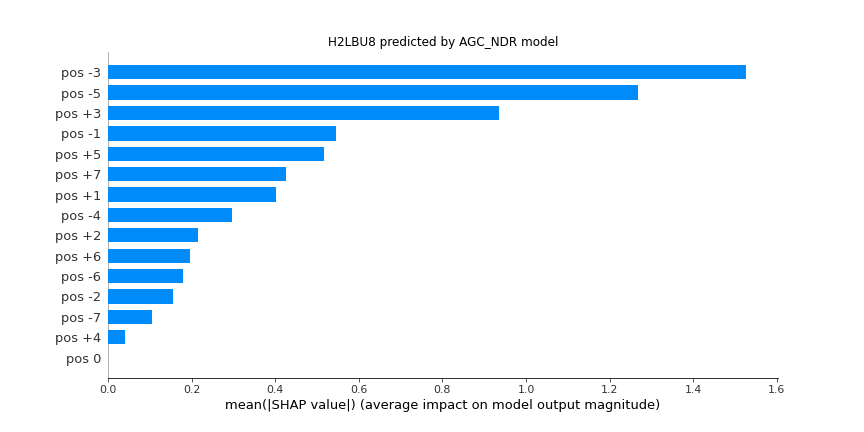 | 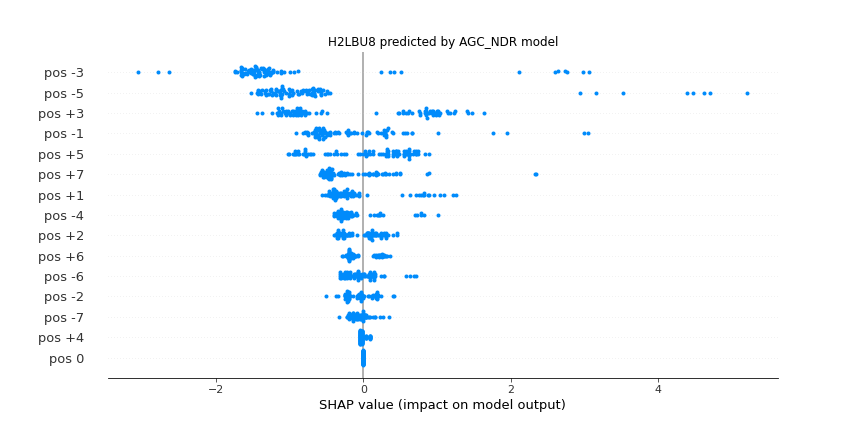 |
| **AGC_PDK1 (STY)** | 99 | 0.908 | 0.9 | 0.903 | 0.908 | 0.901 | 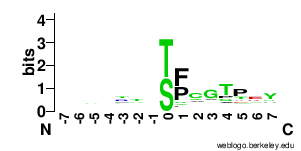 | 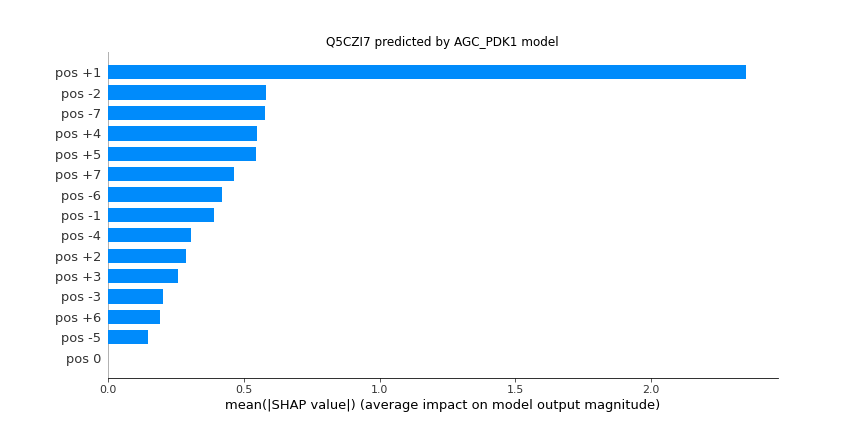 | 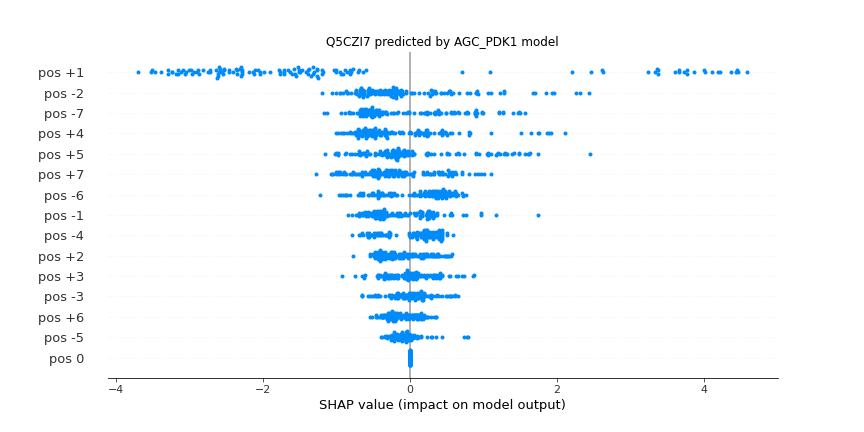 |
| **AGC_PKA (STY)** | 1769 | 0.909 | 0.908 | 0.911 | 0.909 | 0.961 | 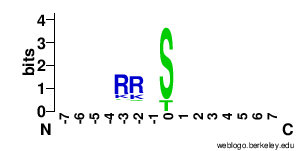 | 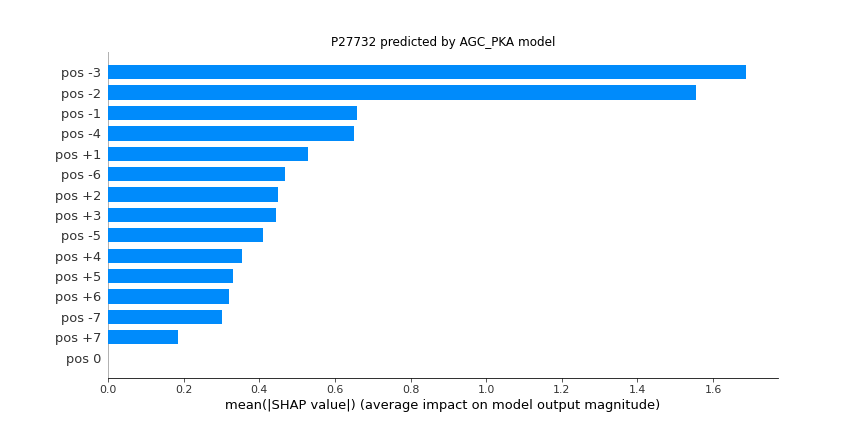 | 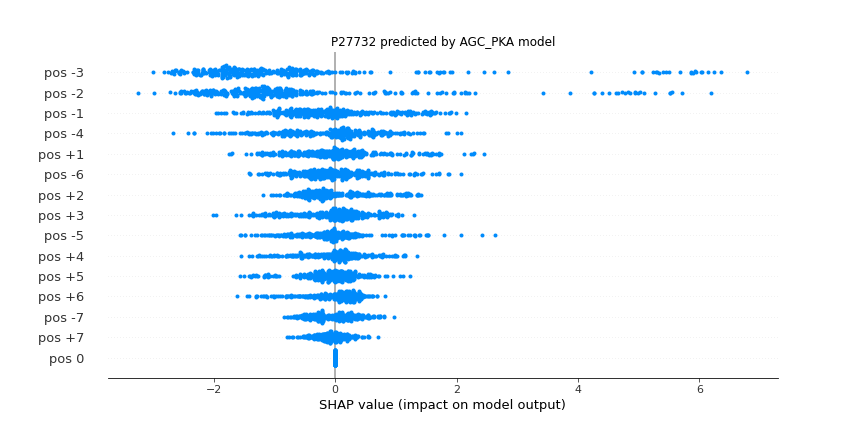 |
| **AGC_PKC (STY)** | 1641 | 0.852 | 0.851 | 0.853 | 0.852 | 0.92 | 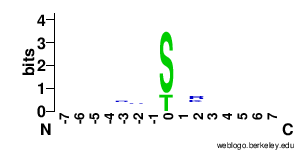 | 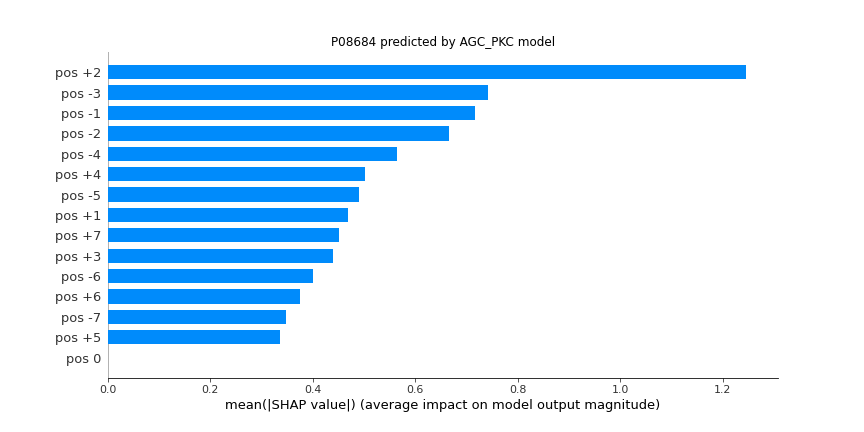 | 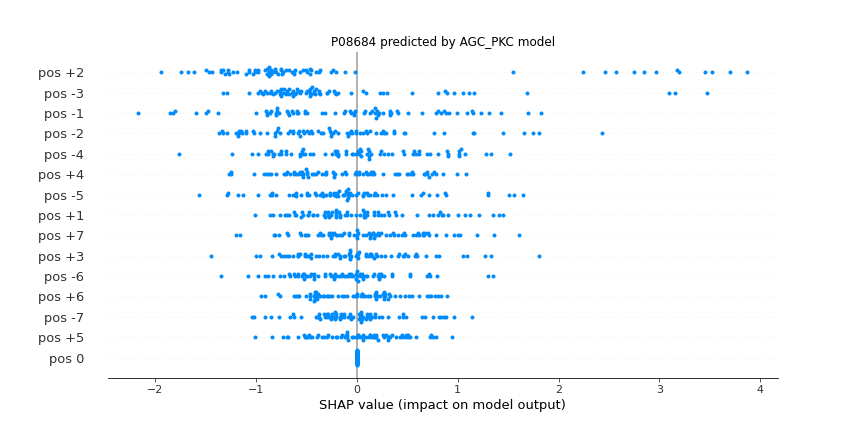 |
| **AGC_PKG (STY)** | 196 | 0.901 | 0.895 | 0.896 | 0.901 | 0.874 | 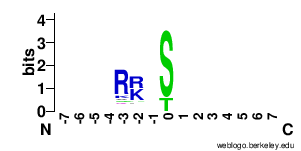 | 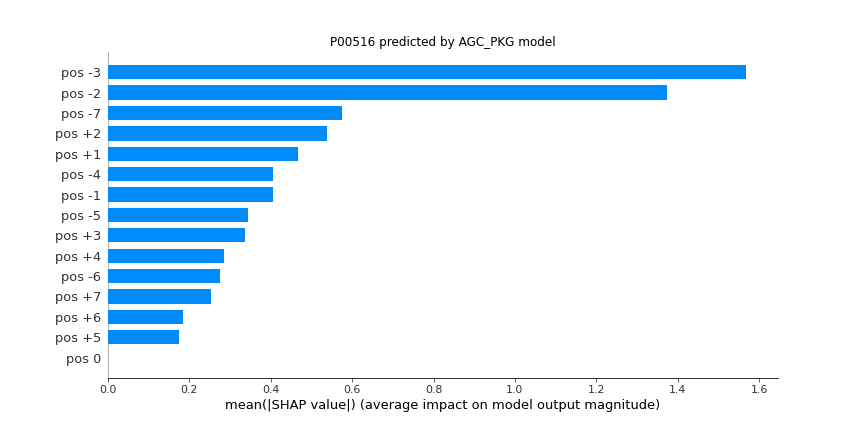 | 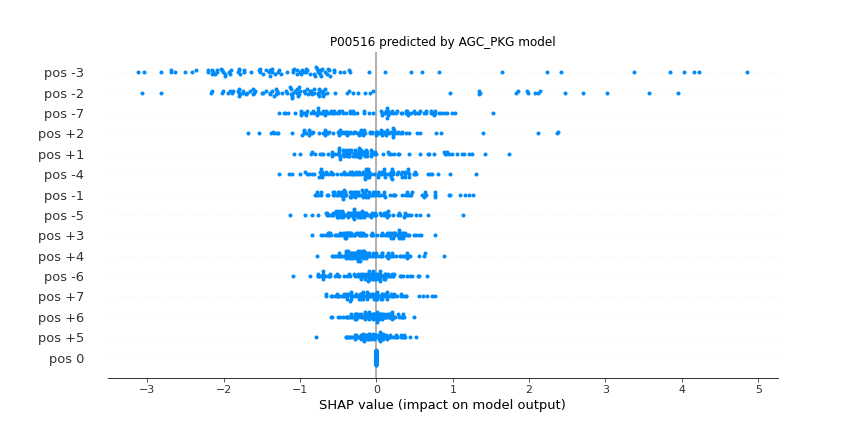 |
| **AGC_PKN (STY)** | 29 | 0.914 | 0.904 | 0.91 | 0.914 | 0.883 | 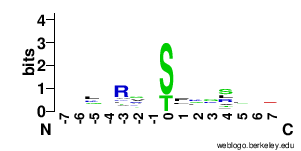 | 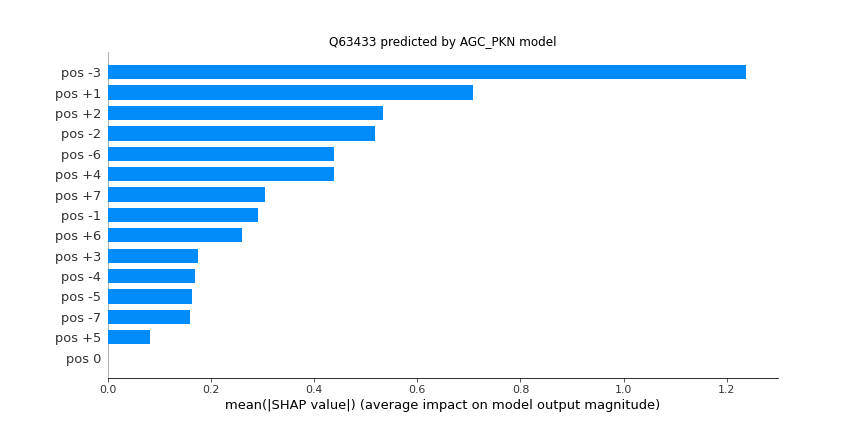 | 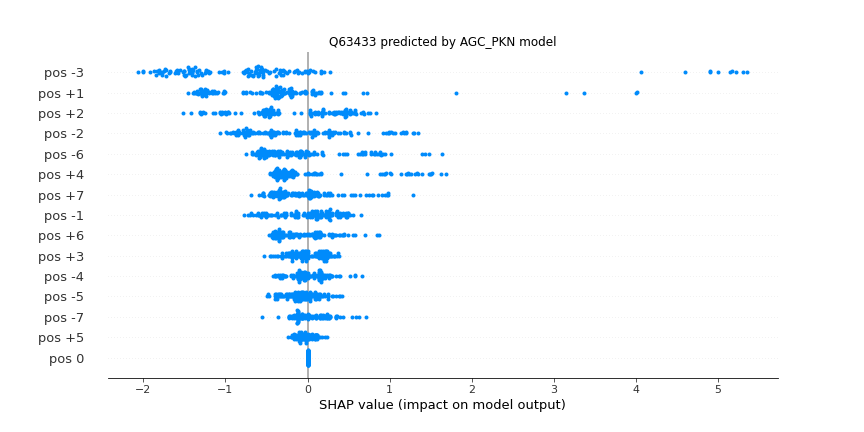 |
| **AGC_RSK (STY)** | 287 | 0.93 | 0.923 | 0.928 | 0.93 | 0.943 | 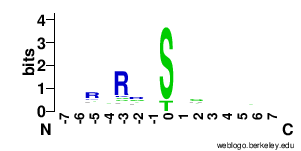 | 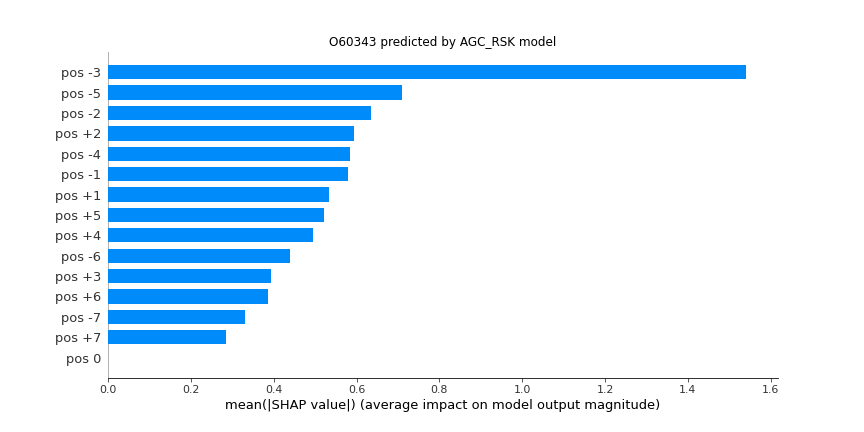 | 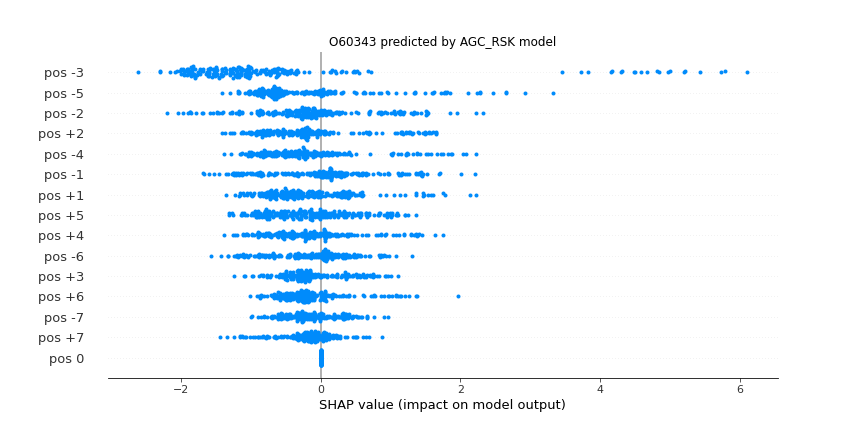 |
| **AGC_SGK (STY)** | 109 | 0.945 | 0.943 | 0.945 | 0.945 | 0.937 | 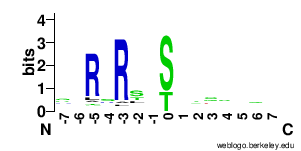 | 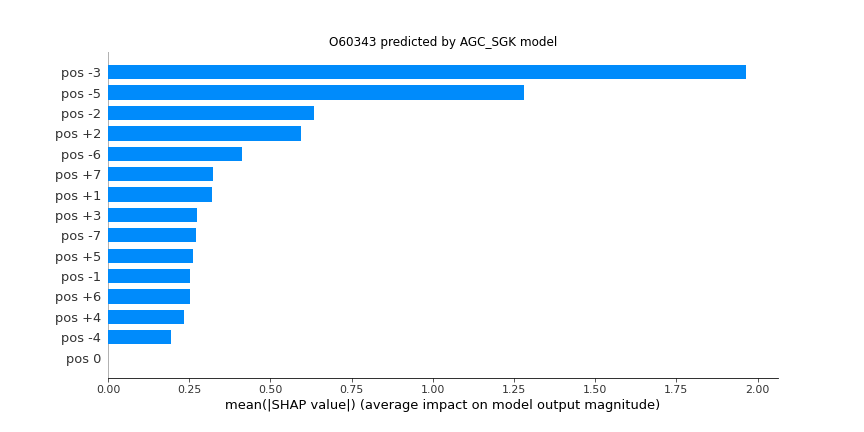 | 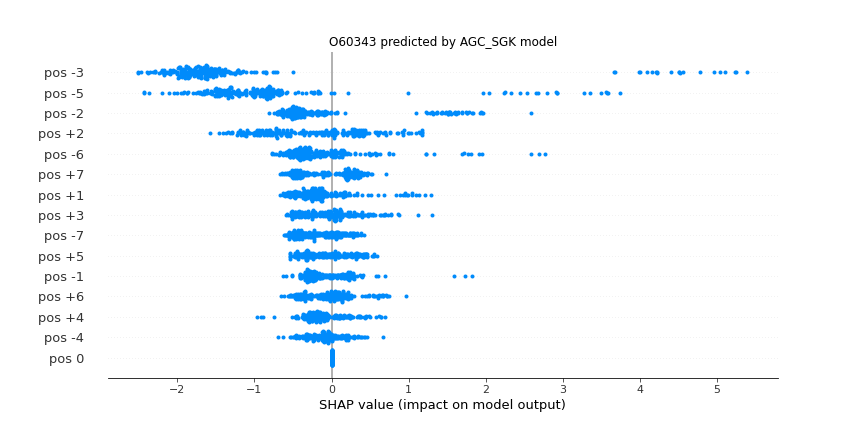 |
| **Atypical_Alpha (STY)** | 90 | 0.853 | 0.821 | 0.825 | 0.853 | 0.74 | 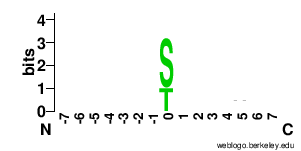 | 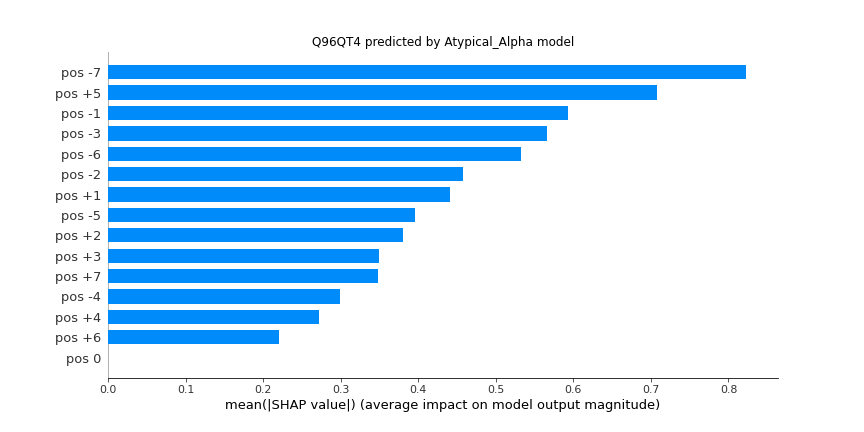 | 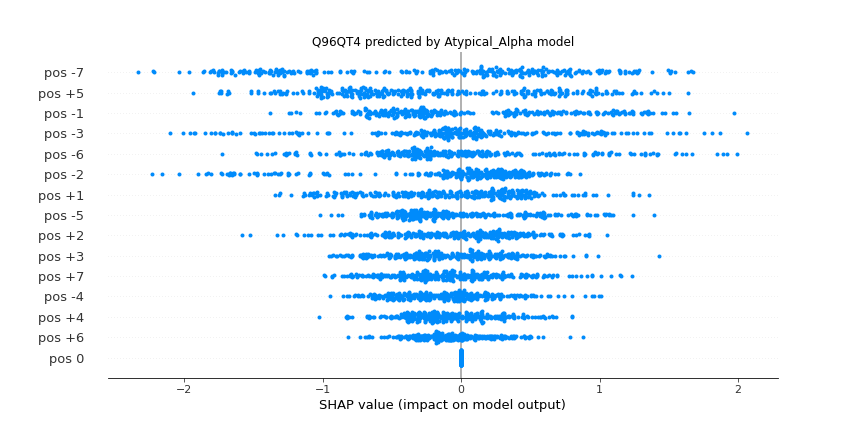 |
| **Atypical_PDHK (STY)** | 43 | 0.911 | 0.904 | 0.913 | 0.911 | 0.887 | 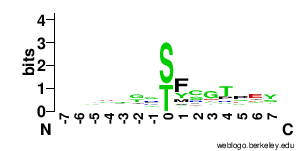 | 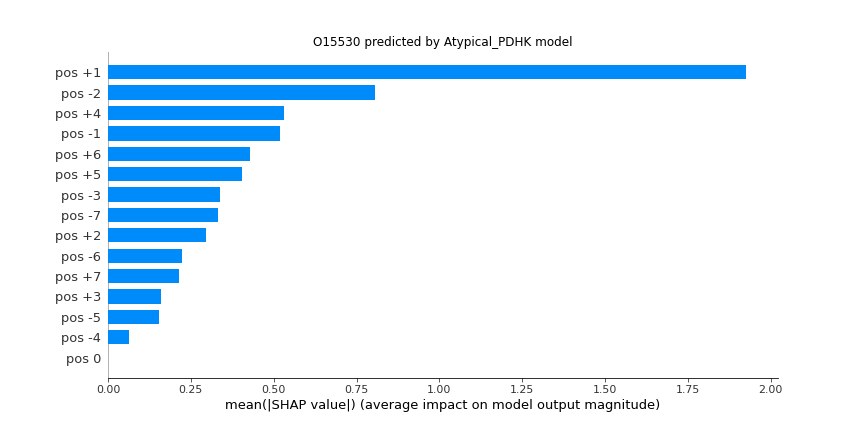 | 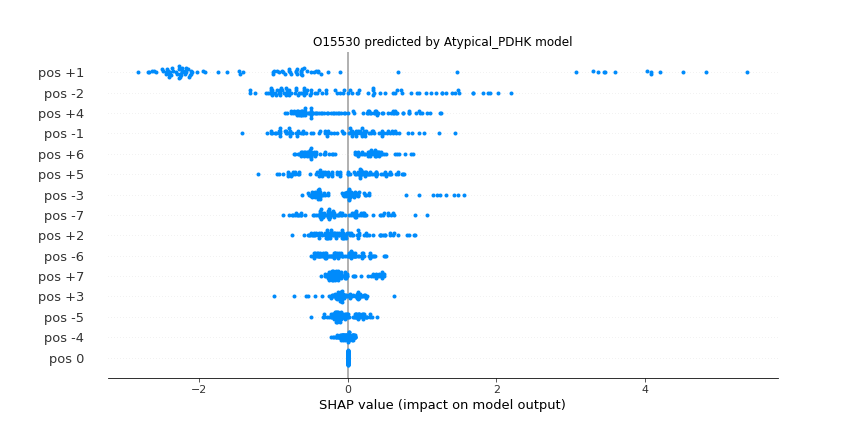 |
| **Atypical_PIKK (STY)** | 857 | 0.919 | 0.916 | 0.919 | 0.919 | 0.953 | 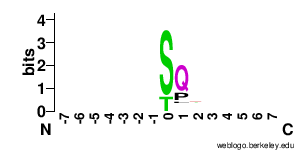 | 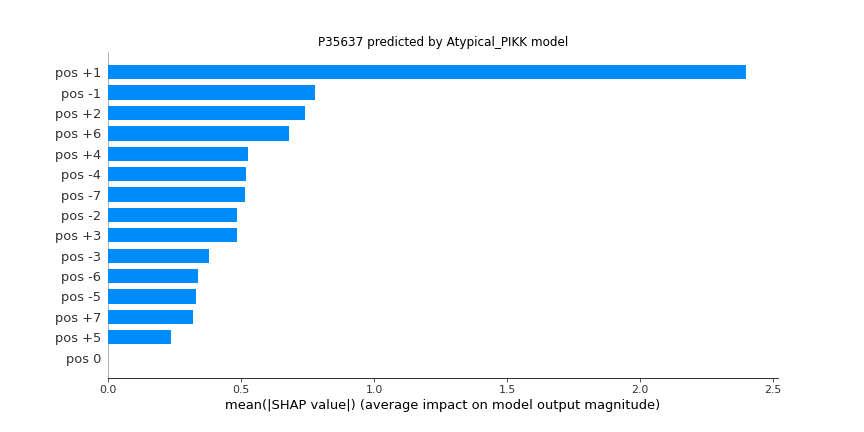 | 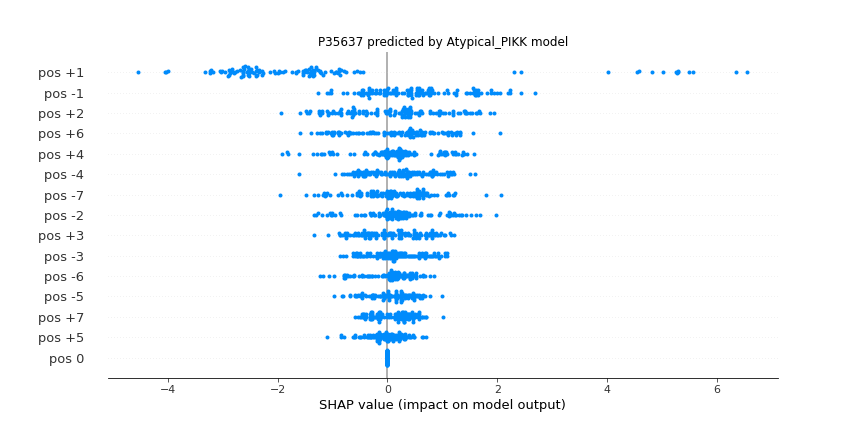 |
| **CAMK_CAMK1 (STY)** | 87 | 0.916 | 0.911 | 0.915 | 0.916 | 0.878 | 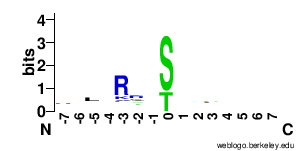 | 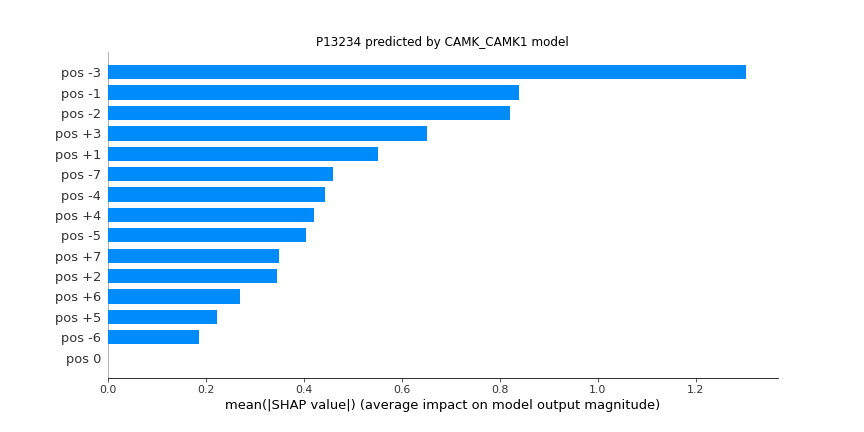 | 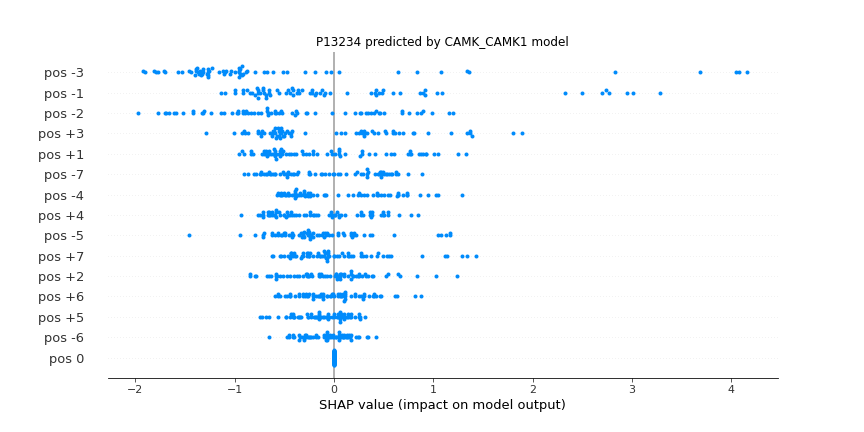 |
| **CAMK_CAMK2 (STY)** | 474 | 0.872 | 0.862 | 0.864 | 0.872 | 0.884 | 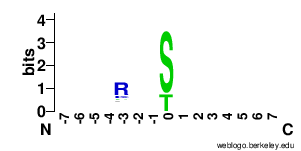 | 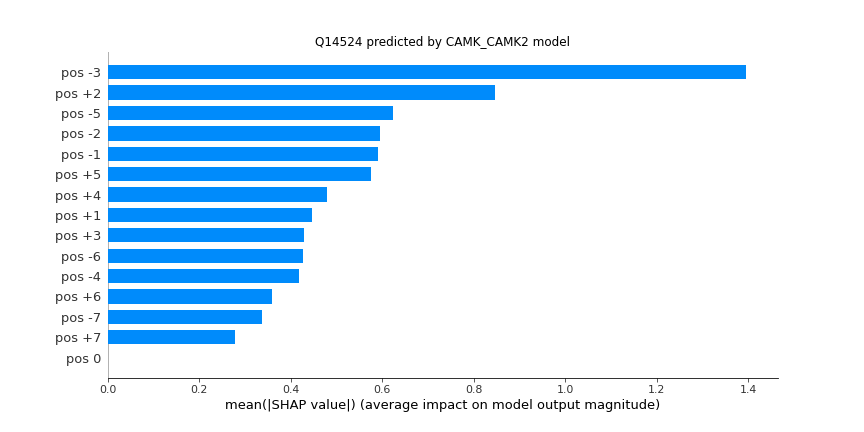 | 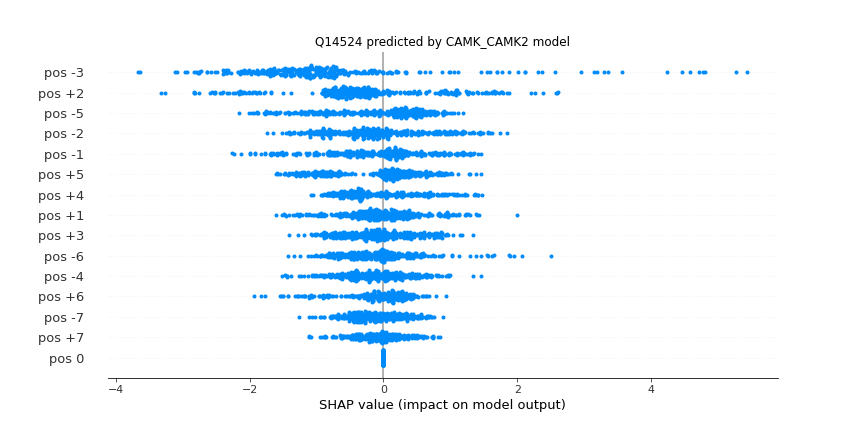 |
| **CAMK_CAMKL (STY)** | 754 | 0.876 | 0.87 | 0.875 | 0.876 | 0.893 | 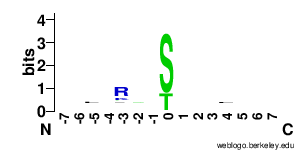 | 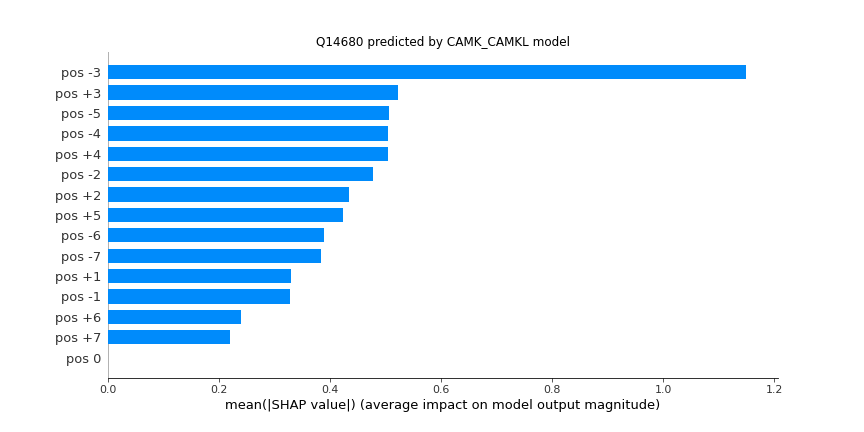 | 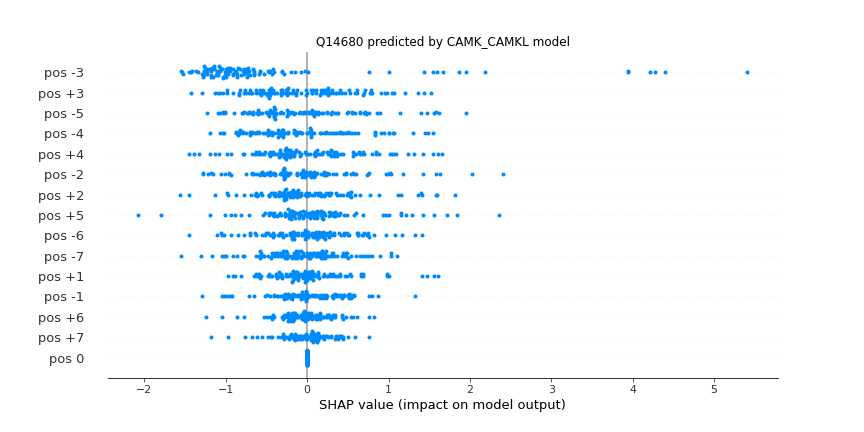 |
| **CAMK_DAPK (STY)** | 69 | 0.867 | 0.848 | 0.85 | 0.867 | 0.785 | 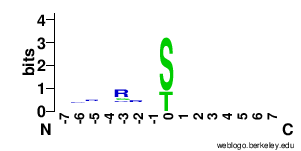 | 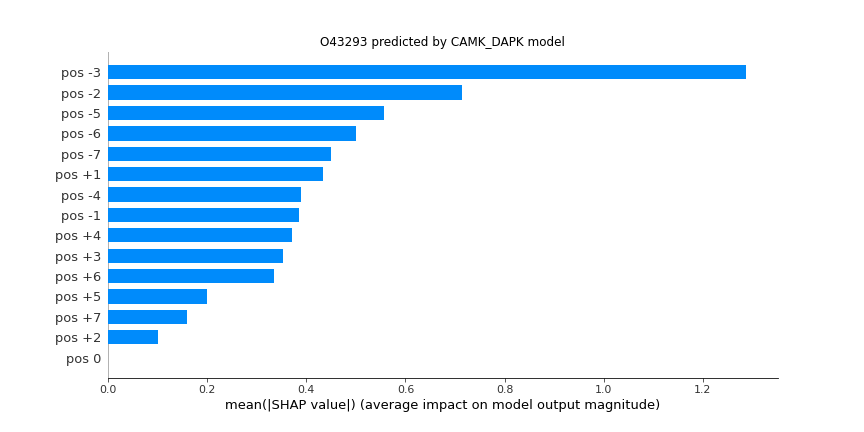 | 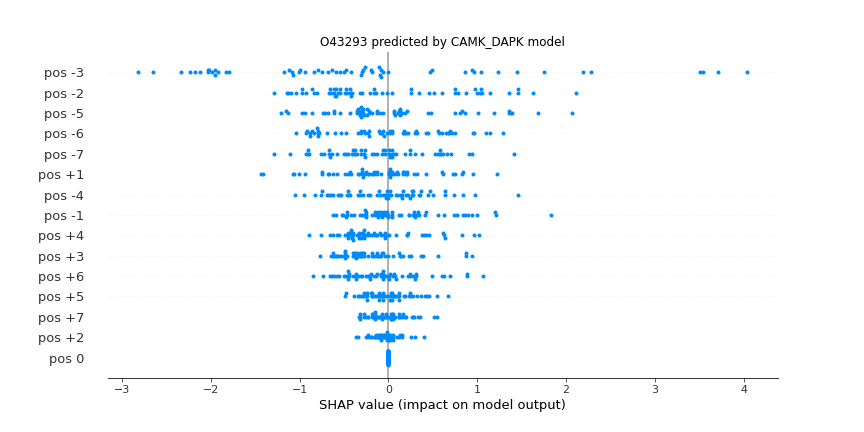 |
| **CAMK_MAPKAPK (STY)** | 184 | 0.902 | 0.895 | 0.897 | 0.902 | 0.895 | 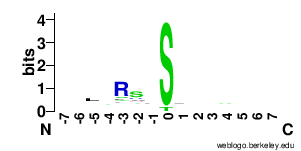 | 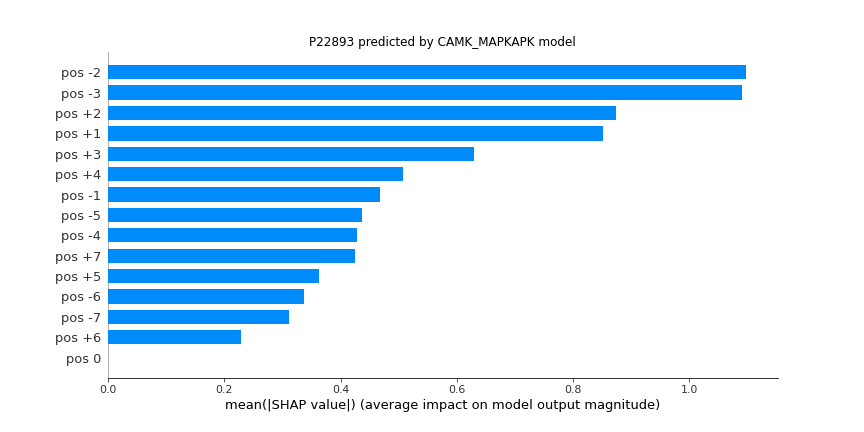 | 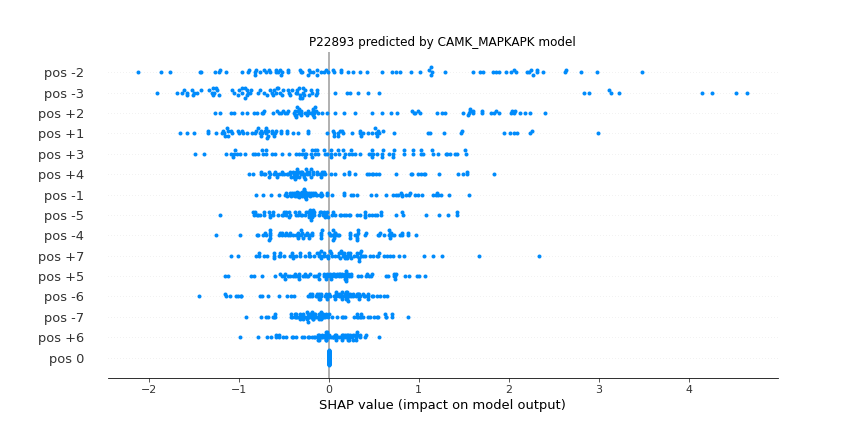 |
| **CAMK_MLCK (STY)** | 34 | 0.884 | 0.854 | 0.846 | 0.884 | 0.73 | 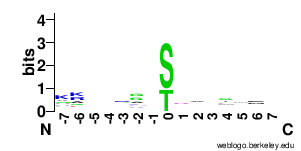 | 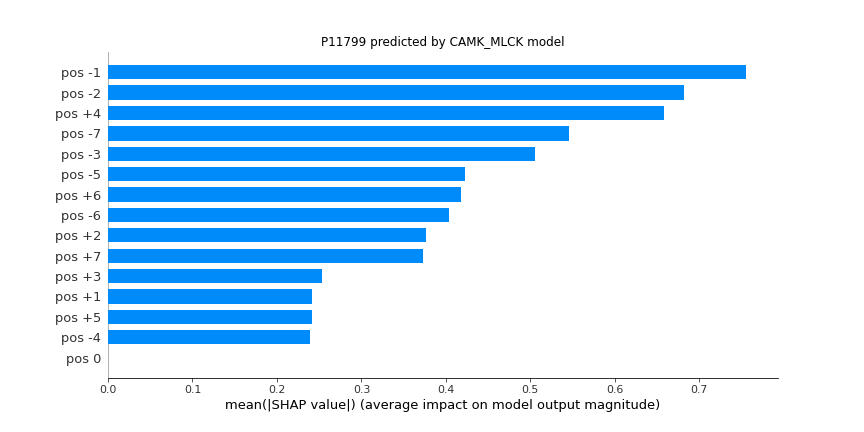 | 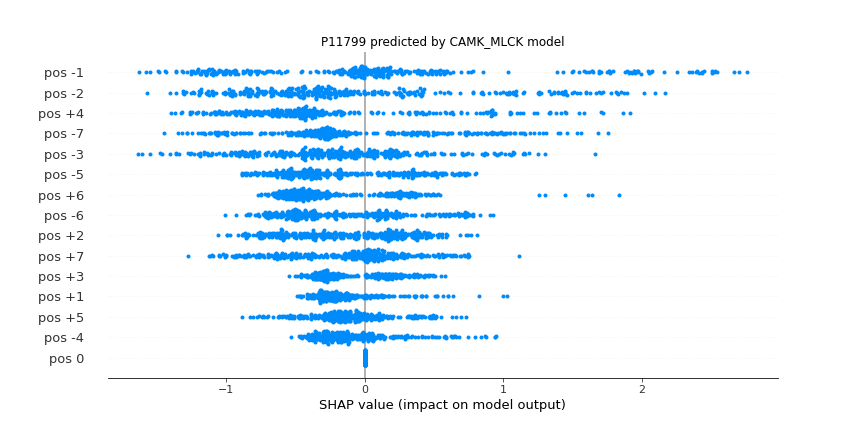 |
| **CAMK_PHK (STY)** | 30 | 0.85 | 0.825 | 0.823 | 0.85 | 0.647 | 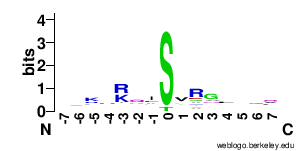 | 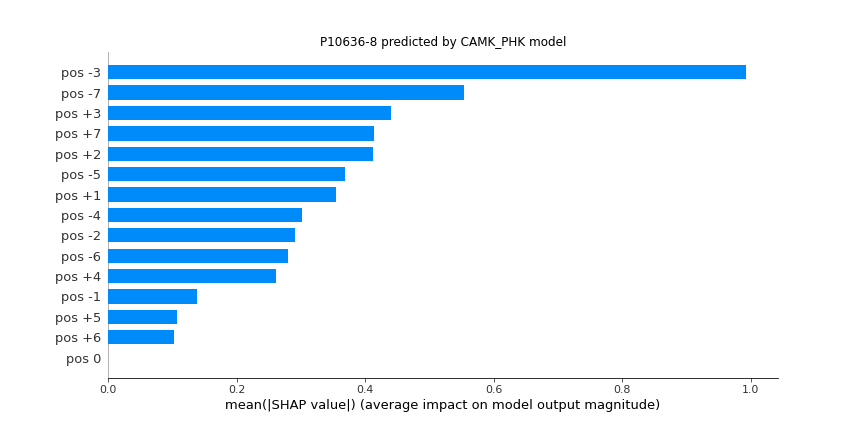 | 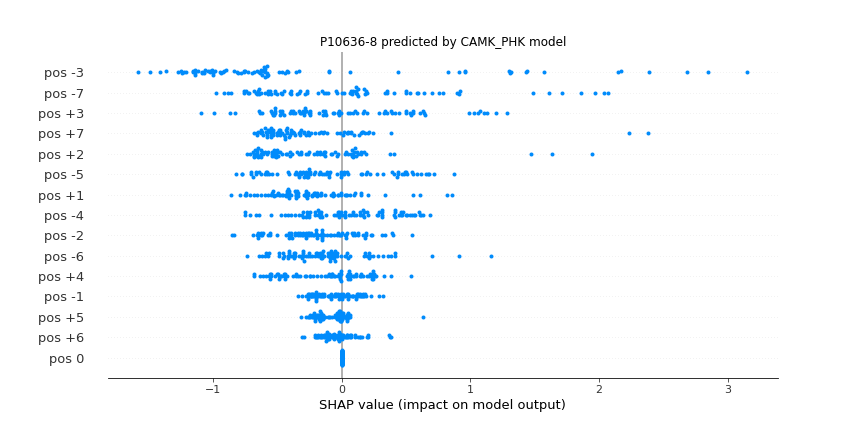 |
| **CAMK_PIM (STY)** | 84 | 0.893 | 0.879 | 0.889 | 0.893 | 0.84 | 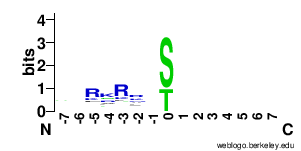 | 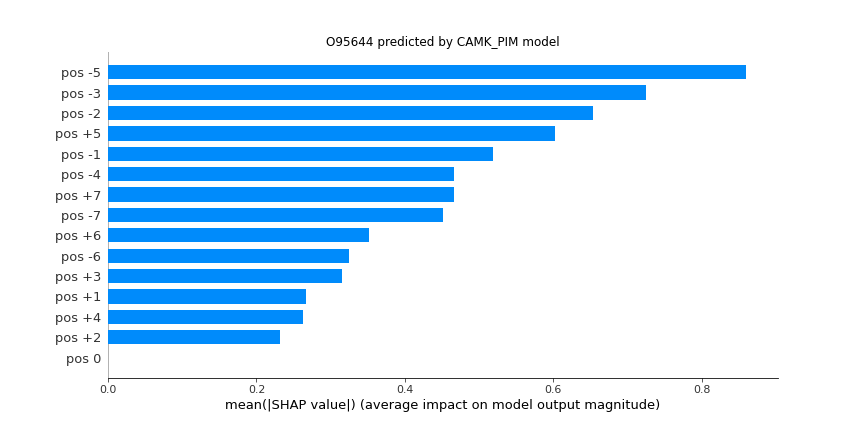 | 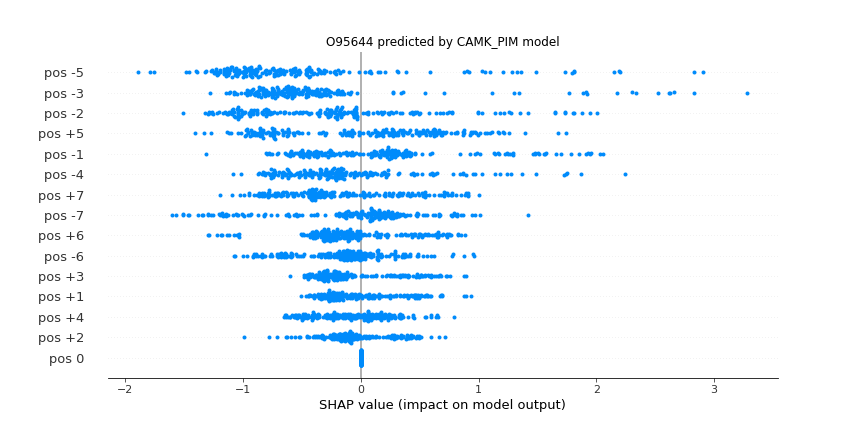 |
| **CAMK_PKD (STY)** | 111 | 0.933 | 0.926 | 0.932 | 0.933 | 0.908 | 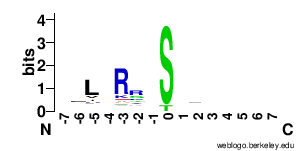 | 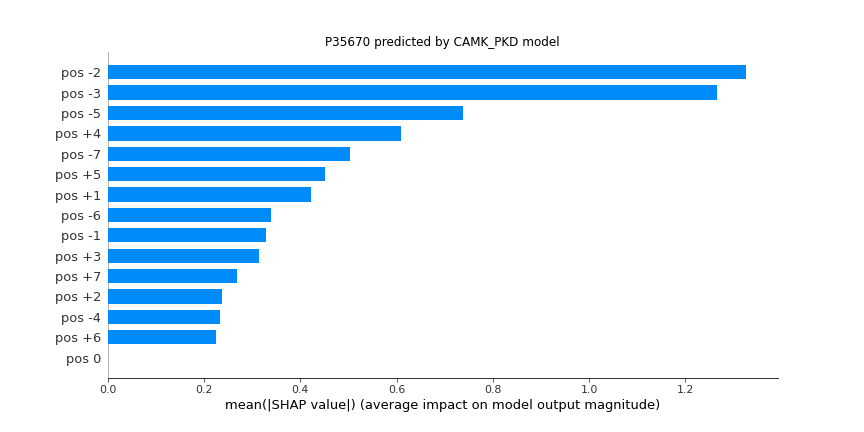 | 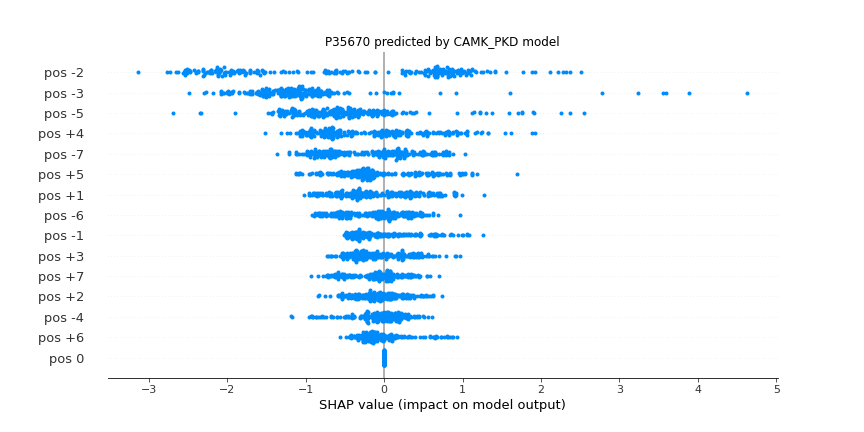 |
| **CAMK_RAD53 (STY)** | 111 | 0.889 | 0.875 | 0.884 | 0.889 | 0.842 | 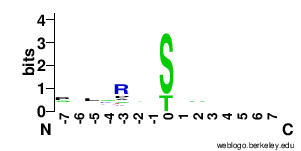 |  |  |
| **CK1_CK1 (STY)** | 464 | 0.855 | 0.847 | 0.851 | 0.855 | 0.873 |  |  |  |
| **CK1_TTBK (STY)** | 19 | 0.826 | 0.784 | 0.754 | 0.826 | 0.693 |  |  |  |
| **CK1_VRK (STY)** | 30 | 0.878 | 0.85 | 0.839 | 0.878 | 0.724 |  |  |  |
| **CMGC_CDK (STY)** | 2050 | 0.937 | 0.936 | 0.941 | 0.937 | 0.98 |  |  |  |
| **CMGC_CK2 (STY)** | 1337 | 0.925 | 0.924 | 0.926 | 0.925 | 0.971 |  |  |  |
| **CMGC_CLK (STY)** | 34 | 0.834 | 0.808 | 0.795 | 0.834 | 0.806 |  |  |  |
| **CMGC_DYRK (STY)** | 244 | 0.935 | 0.933 | 0.935 | 0.935 | 0.937 |  |  |  |
| **CMGC_GSK (STY)** | 572 | 0.923 | 0.921 | 0.922 | 0.923 | 0.952 |  |  |  |
| **CMGC_MAPK (STY)** | 1724 | 0.947 | 0.947 | 0.949 | 0.947 | 0.984 |  |  |  |
| **CMGC_RCK (STY)** | 21 | 0.856 | 0.818 | 0.793 | 0.856 | 0.831 |  |  |  |
| **CMGC_SRPK (STY)** | 34 | 0.951 | 0.947 | 0.951 | 0.951 | 0.944 |  |  |  |
| **Other_Aur (STY)** | 426 | 0.911 | 0.906 | 0.911 | 0.911 | 0.913 |  |  |  |
| **Other_BUB (STY)** | 28 | 0.75 | 0.713 | 0.69 | 0.75 | 0.694 |  |  |  |
| **Other_CAMKK (STY)** | 29 | 0.884 | 0.855 | 0.845 | 0.884 | 0.859 |  |  |  |
| **Other_CDC7 (STY)** | 50 | 0.853 | 0.821 | 0.83 | 0.853 | 0.752 |  |  |  |
| **Other_IKK (STY)** | 326 | 0.839 | 0.819 | 0.826 | 0.839 | 0.816 |  |  |  |
| **Other_KIS (STY)** | 15 | 0.911 | 0.898 | 0.896 | 0.911 | 0.895 |  |  |  |
| **Other_NAK (STY)** | 15 | 0.822 | 0.762 | 0.716 | 0.822 | 0.648 |  |  |  |
| **Other_NEK (STY)** | 103 | 0.861 | 0.833 | 0.837 | 0.861 | 0.799 |  |  |  |
| **Other_NKF2 (STY)** | 21 | 0.906 | 0.867 | 0.843 | 0.906 | 0.693 |  |  |  |
| **Other_PEK (STY)** | 37 | 0.833 | 0.794 | 0.773 | 0.833 | 0.717 |  |  |  |
| **Other_PLK (STY)** | 526 | 0.848 | 0.836 | 0.841 | 0.848 | 0.859 |  |  |  |
| **Other_TTK (STY)** | 87 | 0.824 | 0.772 | 0.763 | 0.824 | 0.613 |  |  |  |
| **Other_ULK (STY)** | 94 | 0.846 | 0.819 | 0.83 | 0.846 | 0.682 |  |  |  |
| **Other_WNK (STY)** | 26 | 0.841 | 0.816 | 0.813 | 0.841 | 0.64 |  |  |  |
| **PKL_FJ (STY)** | 204 | 0.897 | 0.89 | 0.892 | 0.897 | 0.876 |  |  |  |
| **STE_STE-Unique (STY)** | 38 | 0.846 | 0.823 | 0.828 | 0.846 | 0.607 |  |  |  |
| **STE_STE11 (STY)** | 71 | 0.857 | 0.829 | 0.852 | 0.857 | 0.783 |  |  |  |
| **STE_STE20 (STY)** | 457 | 0.854 | 0.839 | 0.851 | 0.854 | 0.838 |  |  |  |
| **STE_STE7 (STY)** | 66 | 0.895 | 0.869 | 0.855 | 0.895 | 0.836 |  |  |  |
| **TKL_IRAK (STY)** | 47 | 0.808 | 0.773 | 0.775 | 0.808 | 0.668 |  |  |  |
| **TKL_LRRK (STY)** | 108 | 0.858 | 0.817 | 0.808 | 0.858 | 0.723 |  |  |  |
| **TKL_MLK (STY)** | 92 | 0.864 | 0.838 | 0.84 | 0.864 | 0.732 |  |  |  |
| **TKL_RAF (STY)** | 34 | 0.809 | 0.766 | 0.732 | 0.809 | 0.623 |  |  |  |
| **TKL_RIPK (STY)** | 34 | 0.853 | 0.825 | 0.833 | 0.853 | 0.705 |  |  |  |
| **TKL_STKR (STY)** | 41 | 0.865 | 0.84 | 0.842 | 0.865 | 0.828 |  |  |  |
| **TK_ALK (STY)** | 15 | 0.8 | 0.741 | 0.691 | 0.8 | 0.523 |  |  |  |
| **TK_Abl (STY)** | 312 | 0.876 | 0.859 | 0.87 | 0.876 | 0.829 |  |  |  |
| **TK_Ack (STY)** | 22 | 0.941 | 0.932 | 0.95 | 0.941 | 0.959 |  |  |  |
| **TK_Axl (STY)** | 20 | 0.883 | 0.846 | 0.823 | 0.883 | 0.71 |  |  |  |
| **TK_Csk (STY)** | 91 | 0.952 | 0.948 | 0.954 | 0.952 | 0.918 |  |  |  |
| **TK_DDR (STY)** | 33 | 0.965 | 0.958 | 0.955 | 0.965 | 0.918 |  |  |  |
| **TK_EGFR (STY)** | 179 | 0.866 | 0.843 | 0.847 | 0.866 | 0.829 |  |  |  |
| **TK_Eph (STY)** | 56 | 0.89 | 0.863 | 0.864 | 0.89 | 0.722 |  |  |  |
| **TK_FAK (STY)** | 156 | 0.919 | 0.908 | 0.917 | 0.919 | 0.895 |  |  |  |
| **TK_FGFR (STY)** | 81 | 0.873 | 0.843 | 0.836 | 0.873 | 0.743 |  |  |  |
| **TK_Fer (STY)** | 42 | 0.878 | 0.859 | 0.847 | 0.878 | 0.807 |  |  |  |
| **TK_InsR (STY)** | 142 | 0.901 | 0.886 | 0.899 | 0.901 | 0.867 |  |  |  |
| **TK_Jak (STY)** | 174 | 0.898 | 0.881 | 0.893 | 0.898 | 0.784 |  |  |  |
| **TK_Met (STY)** | 47 | 0.89 | 0.865 | 0.864 | 0.89 | 0.798 |  |  |  |
| **TK_PDGFR (STY)** | 111 | 0.898 | 0.883 | 0.896 | 0.898 | 0.849 |  |  |  |
| **TK_Ret (STY)** | 36 | 0.867 | 0.843 | 0.844 | 0.867 | 0.775 |  |  |  |
| **TK_Src (STY)** | 1321 | 0.795 | 0.793 | 0.795 | 0.795 | 0.86 |  |  |  |
| **TK_Syk (STY)** | 159 | 0.902 | 0.89 | 0.898 | 0.902 | 0.877 |  |  |  |
| **TK_Tec (STY)** | 84 | 0.863 | 0.829 | 0.856 | 0.863 | 0.729 |  |  |  |
| **TK_Trk (STY)** | 32 | 0.87 | 0.841 | 0.833 | 0.87 | 0.83 |  |  |  |
| **TK_VEGFR (STY)** | 33 | 0.879 | 0.854 | 0.858 | 0.879 | 0.693 |  |  |  |
| **AGC_Akt (ST)** | 526 | 0.947 | 0.946 | 0.947 | 0.947 | 0.971 |  |  |  |
| **AGC_DMPK (ST)** | 154 | 0.905 | 0.893 | 0.901 | 0.905 | 0.878 |  |  |  |
| **AGC_GRK (ST)** | 276 | 0.85 | 0.837 | 0.849 | 0.85 | 0.859 |  |  |  |
| **AGC_NDR (ST)** | 48 | 0.952 | 0.945 | 0.957 | 0.952 | 0.939 |  |  |  |
| **AGC_PDK1 (ST)** | 99 | 0.934 | 0.928 | 0.936 | 0.934 | 0.931 |  |  |  |
| **AGC_PKA (ST)** | 1767 | 0.921 | 0.921 | 0.923 | 0.921 | 0.969 |  |  |  |
| **AGC_PKC (ST)** | 1635 | 0.862 | 0.862 | 0.865 | 0.862 | 0.935 |  |  |  |
| **AGC_PKG (ST)** | 196 | 0.929 | 0.924 | 0.928 | 0.929 | 0.895 |  |  |  |
| **AGC_PKN (ST)** | 29 | 0.885 | 0.87 | 0.863 | 0.885 | 0.843 |  |  |  |
| **AGC_RSK (ST)** | 286 | 0.928 | 0.922 | 0.925 | 0.928 | 0.936 |  |  |  |
| **AGC_SGK (ST)** | 109 | 0.951 | 0.948 | 0.952 | 0.951 | 0.954 |  |  |  |
| **Atypical_Alpha (ST)** | 90 | 0.821 | 0.795 | 0.801 | 0.821 | 0.702 |  |  |  |
| **Atypical_PDHK (ST)** | 43 | 0.915 | 0.908 | 0.915 | 0.915 | 0.88 |  |  |  |
| **Atypical_PIKK (ST)** | 855 | 0.927 | 0.926 | 0.929 | 0.927 | 0.962 |  |  |  |
| **CAMK_CAMK1 (ST)** | 87 | 0.923 | 0.917 | 0.925 | 0.923 | 0.882 |  |  |  |
| **CAMK_CAMK2 (ST)** | 474 | 0.851 | 0.844 | 0.846 | 0.851 | 0.882 |  |  |  |
| **CAMK_CAMKL (ST)** | 747 | 0.882 | 0.878 | 0.884 | 0.882 | 0.908 |  |  |  |
| **CAMK_DAPK (ST)** | 69 | 0.87 | 0.858 | 0.861 | 0.87 | 0.798 |  |  |  |
| **CAMK_MAPKAPK (ST)** | 184 | 0.91 | 0.902 | 0.904 | 0.91 | 0.916 |  |  |  |
| **CAMK_MLCK (ST)** | 34 | 0.864 | 0.831 | 0.813 | 0.864 | 0.756 |  |  |  |
| **CAMK_PHK (ST)** | 30 | 0.872 | 0.848 | 0.853 | 0.872 | 0.716 |  |  |  |
| **CAMK_PIM (ST)** | 84 | 0.887 | 0.876 | 0.884 | 0.887 | 0.833 |  |  |  |
| **CAMK_PKD (ST)** | 111 | 0.943 | 0.938 | 0.942 | 0.943 | 0.932 |  |  |  |
| **CAMK_RAD53 (ST)** | 111 | 0.889 | 0.873 | 0.881 | 0.889 | 0.846 |  |  |  |
| **CK1_CK1 (ST)** | 463 | 0.855 | 0.851 | 0.854 | 0.855 | 0.892 |  |  |  |
| **CK1_TTBK (ST)** | 18 | 0.835 | 0.792 | 0.762 | 0.835 | 0.689 |  |  |  |
| **CK1_VRK (ST)** | 30 | 0.889 | 0.864 | 0.87 | 0.889 | 0.691 |  |  |  |
| **CMGC_CDK (ST)** | 2046 | 0.946 | 0.946 | 0.95 | 0.946 | 0.985 |  |  |  |
| **CMGC_CK2 (ST)** | 1332 | 0.924 | 0.924 | 0.925 | 0.924 | 0.974 |  |  |  |
| **CMGC_CLK (ST)** | 32 | 0.863 | 0.829 | 0.822 | 0.863 | 0.896 |  |  |  |
| **CMGC_DYRK (ST)** | 234 | 0.943 | 0.941 | 0.943 | 0.943 | 0.953 |  |  |  |
| **CMGC_GSK (ST)** | 571 | 0.92 | 0.919 | 0.92 | 0.92 | 0.96 |  |  |  |
| **CMGC_MAPK (ST)** | 1715 | 0.953 | 0.953 | 0.956 | 0.953 | 0.988 |  |  |  |
| **CMGC_RCK (ST)** | 18 | 0.863 | 0.833 | 0.813 | 0.863 | 0.833 |  |  |  |
| **CMGC_SRPK (ST)** | 34 | 0.911 | 0.906 | 0.91 | 0.911 | 0.896 |  |  |  |
| **Other_Aur (ST)** | 424 | 0.907 | 0.902 | 0.91 | 0.907 | 0.925 |  |  |  |
| **Other_BUB (ST)** | 28 | 0.743 | 0.703 | 0.69 | 0.743 | 0.6 |  |  |  |
| **Other_CAMKK (ST)** | 29 | 0.884 | 0.859 | 0.845 | 0.884 | 0.835 |  |  |  |
| **Other_CDC7 (ST)** | 50 | 0.857 | 0.828 | 0.823 | 0.857 | 0.779 |  |  |  |
| **Other_IKK (ST)** | 326 | 0.841 | 0.829 | 0.834 | 0.841 | 0.851 |  |  |  |
| **Other_KIS (ST)** | 15 | 0.878 | 0.863 | 0.873 | 0.878 | 0.889 |  |  |  |
| **Other_NEK (ST)** | 100 | 0.845 | 0.821 | 0.823 | 0.845 | 0.777 |  |  |  |
| **Other_NKF2 (ST)** | 21 | 0.874 | 0.842 | 0.821 | 0.874 | 0.683 |  |  |  |
| **Other_PEK (ST)** | 30 | 0.839 | 0.789 | 0.76 | 0.839 | 0.613 |  |  |  |
| **Other_PLK (ST)** | 525 | 0.829 | 0.821 | 0.824 | 0.829 | 0.867 |  |  |  |
| **Other_TTK (ST)** | 86 | 0.829 | 0.787 | 0.791 | 0.829 | 0.7 |  |  |  |
| **Other_ULK (ST)** | 94 | 0.826 | 0.791 | 0.778 | 0.826 | 0.644 |  |  |  |
| **Other_WNK (ST)** | 26 | 0.855 | 0.826 | 0.818 | 0.855 | 0.691 |  |  |  |
| **PKL_FJ (ST)** | 204 | 0.893 | 0.886 | 0.888 | 0.893 | 0.884 |  |  |  |
| **STE_STE-Unique (ST)** | 37 | 0.874 | 0.852 | 0.858 | 0.874 | 0.715 |  |  |  |
| **STE_STE11 (ST)** | 69 | 0.848 | 0.819 | 0.83 | 0.848 | 0.785 |  |  |  |
| **STE_STE20 (ST)** | 457 | 0.833 | 0.822 | 0.828 | 0.833 | 0.829 |  |  |  |
| **STE_STE7 (ST)** | 46 | 0.9 | 0.876 | 0.887 | 0.9 | 0.834 |  |  |  |
| **TKL_IRAK (ST)** | 47 | 0.796 | 0.748 | 0.735 | 0.796 | 0.68 |  |  |  |
| **TKL_LRRK (ST)** | 105 | 0.845 | 0.811 | 0.814 | 0.845 | 0.737 |  |  |  |
| **TKL_MLK (ST)** | 92 | 0.862 | 0.838 | 0.852 | 0.862 | 0.727 |  |  |  |
| **TKL_RAF (ST)** | 34 | 0.839 | 0.797 | 0.782 | 0.839 | 0.612 |  |  |  |
| **TKL_RIPK (ST)** | 33 | 0.828 | 0.794 | 0.777 | 0.828 | 0.739 |  |  |  |
| **TKL_STKR (ST)** | 38 | 0.873 | 0.843 | 0.838 | 0.873 | 0.821 |  |  |  |
| **TK_DDR (ST)** | 30 | 0.978 | 0.976 | 0.976 | 0.978 | 0.969 |  |  |  |
| **TK_FAK (ST)** | 86 | 0.957 | 0.955 | 0.958 | 0.957 | 0.94 |  |  |  |
| **STE_STE7 (Y)** | 20 | 0.874 | 0.855 | 0.857 | 0.874 | 0.84 |  |  |  |
| **TK_Abl (Y)** | 311 | 0.767 | 0.764 | 0.768 | 0.767 | 0.827 |  |  |  |
| **TK_Ack (Y)** | 21 | 0.9 | 0.886 | 0.891 | 0.9 | 0.879 |  |  |  |
| **TK_Axl (Y)** | 18 | 0.861 | 0.837 | 0.824 | 0.861 | 0.706 |  |  |  |
| **TK_Csk (Y)** | 91 | 0.859 | 0.855 | 0.87 | 0.859 | 0.927 |  |  |  |
| **TK_EGFR (Y)** | 178 | 0.807 | 0.796 | 0.804 | 0.807 | 0.836 |  |  |  |
| **TK_Eph (Y)** | 56 | 0.784 | 0.761 | 0.758 | 0.784 | 0.735 |  |  |  |
| **TK_FAK (Y)** | 70 | 0.844 | 0.827 | 0.831 | 0.844 | 0.787 |  |  |  |
| **TK_FGFR (Y)** | 81 | 0.801 | 0.771 | 0.762 | 0.801 | 0.752 |  |  |  |
| **TK_Fer (Y)** | 42 | 0.861 | 0.844 | 0.853 | 0.861 | 0.838 |  |  |  |
| **TK_InsR (Y)** | 139 | 0.804 | 0.789 | 0.808 | 0.804 | 0.838 |  |  |  |
| **TK_Jak (Y)** | 172 | 0.788 | 0.778 | 0.792 | 0.788 | 0.802 |  |  |  |
| **TK_Met (Y)** | 47 | 0.829 | 0.814 | 0.83 | 0.829 | 0.78 |  |  |  |
| **TK_PDGFR (Y)** | 111 | 0.78 | 0.772 | 0.781 | 0.78 | 0.822 |  |  |  |
| **TK_Ret (Y)** | 32 | 0.843 | 0.798 | 0.779 | 0.843 | 0.772 |  |  |  |
| **TK_Src (Y)** | 1310 | 0.853 | 0.844 | 0.848 | 0.853 | 0.903 |  |  |  |
| **TK_Syk (Y)** | 154 | 0.838 | 0.833 | 0.846 | 0.838 | 0.877 |  |  |  |
| **TK_Tec (Y)** | 80 | 0.801 | 0.776 | 0.791 | 0.801 | 0.691 |  |  |  |
| **TK_Trk (Y)** | 32 | 0.876 | 0.851 | 0.853 | 0.876 | 0.814 |  |  |  |
| **TK_VEGFR (Y)** | 33 | 0.824 | 0.794 | 0.804 | 0.824 | 0.756 |  |  |  |
| **A6QLB8 (STY)** | 16 | 0.78 | 0.752 | 0.781 | 0.78 | 0.833 |  |  |  |
| **A9UF07 (STY)** | 18 | 0.797 | 0.751 | 0.715 | 0.797 | 0.578 |  |  |  |
| **G3N1T2 (STY)** | 24 | 0.855 | 0.82 | 0.809 | 0.855 | 0.828 |  |  |  |
| **O00141 (STY)** | 76 | 0.952 | 0.951 | 0.955 | 0.952 | 0.955 |  |  |  |
| **O00311 (STY)** | 48 | 0.837 | 0.809 | 0.797 | 0.837 | 0.71 |  |  |  |
| **O00418 (STY)** | 16 | 0.806 | 0.746 | 0.702 | 0.806 | 0.699 |  |  |  |
| **O00444 (STY)** | 15 | 0.767 | 0.723 | 0.686 | 0.767 | 0.614 |  |  |  |
| **O14757 (STY)** | 205 | 0.885 | 0.876 | 0.879 | 0.885 | 0.86 |  |  |  |
| **O14920 (STY)** | 96 | 0.832 | 0.792 | 0.792 | 0.832 | 0.74 |  |  |  |
| **O14965 (STY)** | 145 | 0.891 | 0.88 | 0.884 | 0.891 | 0.835 |  |  |  |
| **O15111 (STY)** | 56 | 0.824 | 0.782 | 0.77 | 0.824 | 0.683 |  |  |  |
| **O15264 (STY)** | 38 | 0.922 | 0.915 | 0.916 | 0.922 | 0.893 |  |  |  |
| **O15530 (STY)** | 76 | 0.917 | 0.91 | 0.913 | 0.917 | 0.918 |  |  |  |
| **O43293 (STY)** | 24 | 0.889 | 0.868 | 0.869 | 0.889 | 0.742 |  |  |  |
| **O43318 (STY)** | 43 | 0.861 | 0.832 | 0.835 | 0.861 | 0.747 |  |  |  |
| **O43683 (STY)** | 27 | 0.788 | 0.756 | 0.757 | 0.788 | 0.647 |  |  |  |
| **O55099 (STY)** | 18 | 0.87 | 0.861 | 0.876 | 0.87 | 0.811 |  |  |  |
| **O55173 (STY)** | 31 | 0.935 | 0.919 | 0.927 | 0.935 | 0.911 |  |  |  |
| **O60674 (STY)** | 61 | 0.82 | 0.769 | 0.743 | 0.82 | 0.65 |  |  |  |
| **O70126 (STY)** | 34 | 0.883 | 0.87 | 0.869 | 0.883 | 0.862 |  |  |  |
| **O70405 (STY)** | 17 | 0.795 | 0.75 | 0.717 | 0.795 | 0.495 |  |  |  |
| **O75116 (STY)** | 48 | 0.861 | 0.83 | 0.846 | 0.861 | 0.758 |  |  |  |
| **O75385 (STY)** | 80 | 0.812 | 0.778 | 0.762 | 0.812 | 0.739 |  |  |  |
| **O75582 (STY)** | 33 | 0.899 | 0.887 | 0.889 | 0.899 | 0.857 |  |  |  |
| **O88351 (STY)** | 39 | 0.811 | 0.775 | 0.76 | 0.811 | 0.638 |  |  |  |
| **O88643 (STY)** | 22 | 0.848 | 0.813 | 0.793 | 0.848 | 0.797 |  |  |  |
| **O95747 (STY)** | 15 | 0.933 | 0.917 | 0.908 | 0.933 | 0.879 |  |  |  |
| **O95835 (STY)** | 17 | 0.952 | 0.94 | 0.939 | 0.952 | 0.904 |  |  |  |
| **O96013 (STY)** | 21 | 0.849 | 0.818 | 0.806 | 0.849 | 0.806 |  |  |  |
| **O96017 (STY)** | 74 | 0.878 | 0.858 | 0.874 | 0.878 | 0.85 |  |  |  |
| **P00516 (STY)** | 54 | 0.919 | 0.907 | 0.929 | 0.919 | 0.873 |  |  |  |
| **P00517 (STY)** | 216 | 0.91 | 0.907 | 0.907 | 0.91 | 0.929 |  |  |  |
| **P00519 (STY)** | 226 | 0.86 | 0.841 | 0.846 | 0.86 | 0.782 |  |  |  |
| **P00520 (STY)** | 82 | 0.843 | 0.817 | 0.828 | 0.843 | 0.775 |  |  |  |
| **P00523 (STY)** | 47 | 0.848 | 0.825 | 0.814 | 0.848 | 0.785 |  |  |  |
| **P00533 (STY)** | 117 | 0.833 | 0.794 | 0.776 | 0.833 | 0.726 |  |  |  |
| **P00546 (STY)** | 93 | 0.871 | 0.854 | 0.862 | 0.871 | 0.779 |  |  |  |
| **P04049 (STY)** | 20 | 0.875 | 0.852 | 0.843 | 0.875 | 0.77 |  |  |  |
| **P04409 (STY)** | 59 | 0.878 | 0.864 | 0.883 | 0.878 | 0.892 |  |  |  |
| **P04551 (STY)** | 38 | 0.965 | 0.965 | 0.968 | 0.965 | 0.993 |  |  |  |
| **P04626 (STY)** | 17 | 0.853 | 0.798 | 0.763 | 0.853 | 0.708 |  |  |  |
| **P04629 (STY)** | 15 | 0.811 | 0.76 | 0.724 | 0.811 | 0.564 |  |  |  |
| **P05129 (STY)** | 56 | 0.828 | 0.797 | 0.774 | 0.828 | 0.773 |  |  |  |
| **P05132 (STY)** | 194 | 0.93 | 0.928 | 0.929 | 0.93 | 0.94 |  |  |  |
| **P05480 (STY)** | 159 | 0.868 | 0.85 | 0.855 | 0.868 | 0.855 |  |  |  |
| **P05696 (STY)** | 187 | 0.881 | 0.87 | 0.87 | 0.881 | 0.858 |  |  |  |
| **P05771 (STY)** | 101 | 0.866 | 0.843 | 0.852 | 0.866 | 0.764 |  |  |  |
| **P06213 (STY)** | 73 | 0.838 | 0.802 | 0.797 | 0.838 | 0.739 |  |  |  |
| **P06239 (STY)** | 134 | 0.866 | 0.842 | 0.857 | 0.866 | 0.811 |  |  |  |
| **P06240 (STY)** | 34 | 0.873 | 0.857 | 0.879 | 0.873 | 0.809 |  |  |  |
| **P06241 (STY)** | 207 | 0.86 | 0.839 | 0.843 | 0.86 | 0.808 |  |  |  |
| **P06493 (STY)** | 760 | 0.959 | 0.959 | 0.959 | 0.959 | 0.971 |  |  |  |
| **P07947 (STY)** | 25 | 0.813 | 0.764 | 0.73 | 0.813 | 0.611 |  |  |  |
| **P07948 (STY)** | 143 | 0.859 | 0.833 | 0.839 | 0.859 | 0.759 |  |  |  |
| **P07949 (STY)** | 31 | 0.844 | 0.801 | 0.772 | 0.844 | 0.762 |  |  |  |
| **P08069 (STY)** | 29 | 0.799 | 0.767 | 0.745 | 0.799 | 0.655 |  |  |  |
| **P08103 (STY)** | 15 | 0.811 | 0.754 | 0.71 | 0.811 | 0.804 |  |  |  |
| **P08413 (STY)** | 64 | 0.841 | 0.809 | 0.793 | 0.841 | 0.769 |  |  |  |
| **P08581 (STY)** | 36 | 0.857 | 0.835 | 0.835 | 0.857 | 0.808 |  |  |  |
| **P08631 (STY)** | 36 | 0.88 | 0.865 | 0.883 | 0.88 | 0.792 |  |  |  |
| **P09215 (STY)** | 23 | 0.855 | 0.824 | 0.816 | 0.855 | 0.715 |  |  |  |
| **P09216 (STY)** | 15 | 0.811 | 0.779 | 0.767 | 0.811 | 0.712 |  |  |  |
| **P09217 (STY)** | 22 | 0.826 | 0.777 | 0.736 | 0.826 | 0.78 |  |  |  |
| **P09619 (STY)** | 34 | 0.829 | 0.807 | 0.809 | 0.829 | 0.75 |  |  |  |
| **P09769 (STY)** | 19 | 0.823 | 0.786 | 0.763 | 0.823 | 0.75 |  |  |  |
| **P0C605 (STY)** | 26 | 0.886 | 0.874 | 0.882 | 0.886 | 0.922 |  |  |  |
| **P11275 (STY)** | 104 | 0.875 | 0.863 | 0.862 | 0.875 | 0.864 |  |  |  |
| **P11309 (STY)** | 64 | 0.888 | 0.874 | 0.882 | 0.888 | 0.812 |  |  |  |
| **P11362 (STY)** | 46 | 0.83 | 0.78 | 0.75 | 0.83 | 0.572 |  |  |  |
| **P11440 (STY)** | 136 | 0.957 | 0.957 | 0.96 | 0.957 | 0.954 |  |  |  |
| **P11798 (STY)** | 73 | 0.888 | 0.876 | 0.886 | 0.888 | 0.846 |  |  |  |
| **P11802 (STY)** | 78 | 0.94 | 0.942 | 0.949 | 0.94 | 0.959 |  |  |  |
| **P12931 (STY)** | 658 | 0.813 | 0.798 | 0.801 | 0.813 | 0.803 |  |  |  |
| **P13234 (STY)** | 17 | 0.834 | 0.814 | 0.809 | 0.834 | 0.922 |  |  |  |
| **P15127 (STY)** | 27 | 0.827 | 0.787 | 0.768 | 0.827 | 0.78 |  |  |  |
| **P15208 (STY)** | 26 | 0.897 | 0.868 | 0.856 | 0.897 | 0.814 |  |  |  |
| **P16054 (STY)** | 28 | 0.845 | 0.817 | 0.805 | 0.845 | 0.818 |  |  |  |
| **P16092 (STY)** | 18 | 0.835 | 0.796 | 0.771 | 0.835 | 0.667 |  |  |  |
| **P16591 (STY)** | 21 | 0.794 | 0.749 | 0.713 | 0.794 | 0.578 |  |  |  |
| **P17157 (STY)** | 20 | 0.958 | 0.96 | 0.973 | 0.958 | 0.97 |  |  |  |
| **P17252 (STY)** | 718 | 0.868 | 0.864 | 0.864 | 0.868 | 0.903 |  |  |  |
| **P17612 (STY)** | 930 | 0.909 | 0.907 | 0.908 | 0.909 | 0.949 |  |  |  |
| **P18265 (STY)** | 20 | 0.9 | 0.874 | 0.863 | 0.9 | 0.925 |  |  |  |
| **P18266 (STY)** | 62 | 0.911 | 0.901 | 0.898 | 0.911 | 0.896 |  |  |  |
| **P18653 (STY)** | 36 | 0.922 | 0.919 | 0.927 | 0.922 | 0.931 |  |  |  |
| **P18654 (STY)** | 25 | 0.86 | 0.844 | 0.835 | 0.86 | 0.84 |  |  |  |
| **P19139 (STY)** | 75 | 0.909 | 0.899 | 0.906 | 0.909 | 0.893 |  |  |  |
| **P19525 (STY)** | 25 | 0.82 | 0.767 | 0.725 | 0.82 | 0.61 |  |  |  |
| **P19784 (STY)** | 21 | 0.824 | 0.799 | 0.791 | 0.824 | 0.866 |  |  |  |
| **P20444 (STY)** | 103 | 0.871 | 0.853 | 0.865 | 0.871 | 0.857 |  |  |  |
| **P21708 (STY)** | 67 | 0.955 | 0.956 | 0.959 | 0.955 | 0.952 |  |  |  |
| **P23443 (STY)** | 59 | 0.912 | 0.906 | 0.91 | 0.912 | 0.934 |  |  |  |
| **P23458 (STY)** | 17 | 0.843 | 0.808 | 0.789 | 0.843 | 0.761 |  |  |  |
| **P24723 (STY)** | 18 | 0.835 | 0.804 | 0.787 | 0.835 | 0.683 |  |  |  |
| **P24941 (STY)** | 465 | 0.971 | 0.971 | 0.971 | 0.971 | 0.979 |  |  |  |
| **P25098 (STY)** | 118 | 0.863 | 0.843 | 0.856 | 0.863 | 0.837 |  |  |  |
| **P25911 (STY)** | 51 | 0.879 | 0.853 | 0.861 | 0.879 | 0.847 |  |  |  |
| **P26927 (STY)** | 15 | 0.833 | 0.786 | 0.757 | 0.833 | 0.557 |  |  |  |
| **P27361 (STY)** | 487 | 0.955 | 0.954 | 0.955 | 0.955 | 0.964 |  |  |  |
| **P27791 (STY)** | 143 | 0.909 | 0.903 | 0.904 | 0.909 | 0.912 |  |  |  |
| **P28482 (STY)** | 599 | 0.963 | 0.963 | 0.964 | 0.963 | 0.979 |  |  |  |
| **P28867 (STY)** | 29 | 0.88 | 0.858 | 0.871 | 0.88 | 0.822 |  |  |  |
| **P29597 (STY)** | 17 | 0.783 | 0.749 | 0.725 | 0.783 | 0.444 |  |  |  |
| **P31749 (STY)** | 411 | 0.956 | 0.955 | 0.956 | 0.956 | 0.971 |  |  |  |
| **P31750 (STY)** | 101 | 0.937 | 0.936 | 0.937 | 0.937 | 0.96 |  |  |  |
| **P31751 (STY)** | 79 | 0.875 | 0.862 | 0.873 | 0.875 | 0.876 |  |  |  |
| **P32562 (STY)** | 18 | 0.771 | 0.721 | 0.689 | 0.771 | 0.677 |  |  |  |
| **P32577 (STY)** | 22 | 0.796 | 0.746 | 0.703 | 0.796 | 0.495 |  |  |  |
| **P33674 (STY)** | 18 | 0.917 | 0.904 | 0.897 | 0.917 | 0.978 |  |  |  |
| **P33981 (STY)** | 66 | 0.823 | 0.778 | 0.76 | 0.823 | 0.701 |  |  |  |
| **P34947 (STY)** | 23 | 0.805 | 0.761 | 0.726 | 0.805 | 0.674 |  |  |  |
| **P35465 (STY)** | 25 | 0.86 | 0.839 | 0.825 | 0.86 | 0.796 |  |  |  |
| **P35626 (STY)** | 27 | 0.858 | 0.827 | 0.819 | 0.858 | 0.764 |  |  |  |
| **P36887 (STY)** | 20 | 0.858 | 0.841 | 0.845 | 0.858 | 0.825 |  |  |  |
| **P37173 (STY)** | 16 | 0.813 | 0.776 | 0.75 | 0.813 | 0.525 |  |  |  |
| **P38110 (STY)** | 19 | 0.955 | 0.955 | 0.962 | 0.955 | 0.978 |  |  |  |
| **P38111 (STY)** | 21 | 0.953 | 0.951 | 0.959 | 0.953 | 0.987 |  |  |  |
| **P39688 (STY)** | 65 | 0.833 | 0.794 | 0.768 | 0.833 | 0.723 |  |  |  |
| **P39951 (STY)** | 79 | 0.956 | 0.957 | 0.96 | 0.956 | 0.978 |  |  |  |
| **P41240 (STY)** | 28 | 0.881 | 0.837 | 0.82 | 0.881 | 0.604 |  |  |  |
| **P41241 (STY)** | 22 | 0.835 | 0.785 | 0.75 | 0.835 | 0.647 |  |  |  |
| **P41279 (STY)** | 25 | 0.88 | 0.852 | 0.842 | 0.88 | 0.711 |  |  |  |
| **P41743 (STY)** | 47 | 0.837 | 0.814 | 0.813 | 0.837 | 0.753 |  |  |  |
| **P42345 (STY)** | 142 | 0.885 | 0.873 | 0.882 | 0.885 | 0.827 |  |  |  |
| **P42346 (STY)** | 17 | 0.824 | 0.777 | 0.746 | 0.824 | 0.678 |  |  |  |
| **P42684 (STY)** | 33 | 0.823 | 0.808 | 0.81 | 0.823 | 0.658 |  |  |  |
| **P43250 (STY)** | 16 | 0.794 | 0.759 | 0.74 | 0.794 | 0.519 |  |  |  |
| **P43403 (STY)** | 39 | 0.863 | 0.833 | 0.854 | 0.863 | 0.778 |  |  |  |
| **P43405 (STY)** | 85 | 0.894 | 0.882 | 0.89 | 0.894 | 0.86 |  |  |  |
| **P45983 (STY)** | 229 | 0.964 | 0.964 | 0.965 | 0.964 | 0.971 |  |  |  |
| **P45984 (STY)** | 82 | 0.955 | 0.955 | 0.959 | 0.955 | 0.968 |  |  |  |
| **P45985 (STY)** | 17 | 0.92 | 0.899 | 0.884 | 0.92 | 0.835 |  |  |  |
| **P46196 (STY)** | 17 | 0.883 | 0.867 | 0.864 | 0.883 | 0.828 |  |  |  |
| **P47196 (STY)** | 49 | 0.952 | 0.952 | 0.955 | 0.952 | 0.98 |  |  |  |
| **P47811 (STY)** | 90 | 0.928 | 0.925 | 0.928 | 0.928 | 0.893 |  |  |  |
| **P48025 (STY)** | 38 | 0.873 | 0.848 | 0.862 | 0.873 | 0.803 |  |  |  |
| **P48729 (STY)** | 170 | 0.885 | 0.87 | 0.877 | 0.885 | 0.867 |  |  |  |
| **P48730 (STY)** | 105 | 0.871 | 0.858 | 0.864 | 0.871 | 0.851 |  |  |  |
| **P48734 (STY)** | 36 | 0.945 | 0.945 | 0.949 | 0.945 | 0.983 |  |  |  |
| **P49137 (STY)** | 101 | 0.926 | 0.921 | 0.926 | 0.926 | 0.915 |  |  |  |
| **P49138 (STY)** | 36 | 0.903 | 0.882 | 0.872 | 0.903 | 0.87 |  |  |  |
| **P49185 (STY)** | 42 | 0.92 | 0.92 | 0.927 | 0.92 | 0.953 |  |  |  |
| **P49186 (STY)** | 16 | 0.908 | 0.892 | 0.897 | 0.908 | 0.95 |  |  |  |
| **P49336 (STY)** | 18 | 0.945 | 0.932 | 0.928 | 0.945 | 0.889 |  |  |  |
| **P49615 (STY)** | 105 | 0.959 | 0.959 | 0.962 | 0.959 | 0.95 |  |  |  |
| **P49674 (STY)** | 60 | 0.847 | 0.819 | 0.818 | 0.847 | 0.781 |  |  |  |
| **P49760 (STY)** | 20 | 0.817 | 0.784 | 0.763 | 0.817 | 0.77 |  |  |  |
| **P49840 (STY)** | 110 | 0.938 | 0.937 | 0.94 | 0.938 | 0.959 |  |  |  |
| **P49841 (STY)** | 433 | 0.925 | 0.923 | 0.924 | 0.925 | 0.952 |  |  |  |
| **P50613 (STY)** | 60 | 0.867 | 0.85 | 0.858 | 0.867 | 0.786 |  |  |  |
| **P50750 (STY)** | 51 | 0.885 | 0.874 | 0.875 | 0.885 | 0.818 |  |  |  |
| **P51812 (STY)** | 72 | 0.891 | 0.881 | 0.888 | 0.891 | 0.861 |  |  |  |
| **P51813 (STY)** | 22 | 0.811 | 0.771 | 0.744 | 0.811 | 0.742 |  |  |  |
| **P51955 (STY)** | 35 | 0.81 | 0.765 | 0.737 | 0.81 | 0.637 |  |  |  |
| **P52333 (STY)** | 23 | 0.834 | 0.782 | 0.742 | 0.834 | 0.634 |  |  |  |
| **P52564 (STY)** | 15 | 0.922 | 0.914 | 0.922 | 0.922 | 0.886 |  |  |  |
| **P53350 (STY)** | 340 | 0.869 | 0.849 | 0.86 | 0.869 | 0.812 |  |  |  |
| **P53351 (STY)** | 24 | 0.82 | 0.776 | 0.749 | 0.82 | 0.611 |  |  |  |
| **P53355 (STY)** | 31 | 0.823 | 0.786 | 0.781 | 0.823 | 0.659 |  |  |  |
| **P53778 (STY)** | 41 | 0.947 | 0.947 | 0.954 | 0.947 | 0.987 |  |  |  |
| **P53779 (STY)** | 32 | 0.912 | 0.912 | 0.915 | 0.912 | 0.913 |  |  |  |
| **P54199 (STY)** | 18 | 0.751 | 0.689 | 0.639 | 0.751 | 0.436 |  |  |  |
| **P54645 (STY)** | 30 | 0.878 | 0.86 | 0.871 | 0.878 | 0.88 |  |  |  |
| **P54646 (STY)** | 47 | 0.869 | 0.85 | 0.864 | 0.869 | 0.841 |  |  |  |
| **P57059 (STY)** | 20 | 0.892 | 0.858 | 0.829 | 0.892 | 0.87 |  |  |  |
| **P63085 (STY)** | 267 | 0.966 | 0.966 | 0.967 | 0.966 | 0.977 |  |  |  |
| **P63086 (STY)** | 90 | 0.959 | 0.959 | 0.961 | 0.959 | 0.927 |  |  |  |
| **P67870 (STY)** | 20 | 0.917 | 0.911 | 0.923 | 0.917 | 0.98 |  |  |  |
| **P67999 (STY)** | 21 | 0.937 | 0.93 | 0.941 | 0.937 | 0.913 |  |  |  |
| **P68399 (STY)** | 16 | 0.94 | 0.937 | 0.942 | 0.94 | 0.95 |  |  |  |
| **P68400 (STY)** | 746 | 0.922 | 0.921 | 0.922 | 0.922 | 0.958 |  |  |  |
| **P68403 (STY)** | 25 | 0.82 | 0.784 | 0.76 | 0.82 | 0.776 |  |  |  |
| **P68404 (STY)** | 22 | 0.881 | 0.848 | 0.829 | 0.881 | 0.841 |  |  |  |
| **P70032 (STY)** | 15 | 0.811 | 0.753 | 0.707 | 0.811 | 0.664 |  |  |  |
| **P70335 (STY)** | 27 | 0.932 | 0.924 | 0.939 | 0.932 | 0.802 |  |  |  |
| **P70336 (STY)** | 27 | 0.87 | 0.841 | 0.83 | 0.87 | 0.751 |  |  |  |
| **P70618 (STY)** | 21 | 0.874 | 0.871 | 0.891 | 0.874 | 0.915 |  |  |  |
| **P78527 (STY)** | 136 | 0.914 | 0.907 | 0.912 | 0.914 | 0.887 |  |  |  |
| **P97313 (STY)** | 17 | 0.844 | 0.802 | 0.772 | 0.844 | 0.851 |  |  |  |
| **P97377 (STY)** | 72 | 0.926 | 0.925 | 0.93 | 0.926 | 0.951 |  |  |  |
| **P97633 (STY)** | 15 | 0.8 | 0.748 | 0.707 | 0.8 | 0.57 |  |  |  |
| **Q00526 (STY)** | 16 | 0.918 | 0.906 | 0.904 | 0.918 | 0.95 |  |  |  |
| **Q00534 (STY)** | 41 | 0.935 | 0.936 | 0.941 | 0.935 | 0.98 |  |  |  |
| **Q00535 (STY)** | 246 | 0.969 | 0.969 | 0.97 | 0.969 | 0.972 |  |  |  |
| **Q01279 (STY)** | 26 | 0.833 | 0.799 | 0.774 | 0.833 | 0.758 |  |  |  |
| **Q01314 (STY)** | 20 | 0.917 | 0.915 | 0.927 | 0.917 | 0.96 |  |  |  |
| **Q02111 (STY)** | 17 | 0.843 | 0.799 | 0.773 | 0.843 | 0.603 |  |  |  |
| **Q02156 (STY)** | 113 | 0.878 | 0.863 | 0.866 | 0.878 | 0.808 |  |  |  |
| **Q02399 (STY)** | 16 | 0.95 | 0.947 | 0.96 | 0.95 | 0.944 |  |  |  |
| **Q02956 (STY)** | 34 | 0.864 | 0.837 | 0.833 | 0.864 | 0.832 |  |  |  |
| **Q03114 (STY)** | 73 | 0.959 | 0.959 | 0.962 | 0.959 | 0.965 |  |  |  |
| **Q04759 (STY)** | 61 | 0.864 | 0.842 | 0.842 | 0.864 | 0.793 |  |  |  |
| **Q05397 (STY)** | 29 | 0.827 | 0.791 | 0.772 | 0.827 | 0.765 |  |  |  |
| **Q05513 (STY)** | 121 | 0.844 | 0.811 | 0.814 | 0.844 | 0.751 |  |  |  |
| **Q05655 (STY)** | 177 | 0.866 | 0.847 | 0.852 | 0.866 | 0.82 |  |  |  |
| **Q06187 (STY)** | 26 | 0.847 | 0.813 | 0.8 | 0.847 | 0.709 |  |  |  |
| **Q06226 (STY)** | 15 | 0.9 | 0.875 | 0.862 | 0.9 | 0.83 |  |  |  |
| **Q07014 (STY)** | 18 | 0.825 | 0.78 | 0.754 | 0.825 | 0.783 |  |  |  |
| **Q07832 (STY)** | 49 | 0.844 | 0.798 | 0.782 | 0.844 | 0.787 |  |  |  |
| **Q08881 (STY)** | 15 | 0.8 | 0.741 | 0.693 | 0.8 | 0.461 |  |  |  |
| **Q09137 (STY)** | 35 | 0.867 | 0.845 | 0.857 | 0.867 | 0.849 |  |  |  |
| **Q13043 (STY)** | 34 | 0.819 | 0.78 | 0.751 | 0.819 | 0.628 |  |  |  |
| **Q13131 (STY)** | 203 | 0.896 | 0.884 | 0.889 | 0.896 | 0.883 |  |  |  |
| **Q13153 (STY)** | 91 | 0.867 | 0.856 | 0.862 | 0.867 | 0.856 |  |  |  |
| **Q13164 (STY)** | 32 | 0.886 | 0.868 | 0.871 | 0.886 | 0.914 |  |  |  |
| **Q13177 (STY)** | 50 | 0.833 | 0.796 | 0.773 | 0.833 | 0.746 |  |  |  |
| **Q13188 (STY)** | 28 | 0.816 | 0.792 | 0.778 | 0.816 | 0.66 |  |  |  |
| **Q13237 (STY)** | 29 | 0.919 | 0.912 | 0.932 | 0.919 | 0.957 |  |  |  |
| **Q13315 (STY)** | 321 | 0.957 | 0.957 | 0.958 | 0.957 | 0.976 |  |  |  |
| **Q13464 (STY)** | 86 | 0.89 | 0.874 | 0.871 | 0.89 | 0.902 |  |  |  |
| **Q13535 (STY)** | 124 | 0.952 | 0.952 | 0.953 | 0.952 | 0.966 |  |  |  |
| **Q13554 (STY)** | 22 | 0.909 | 0.898 | 0.91 | 0.909 | 0.829 |  |  |  |
| **Q13557 (STY)** | 26 | 0.878 | 0.861 | 0.876 | 0.878 | 0.819 |  |  |  |
| **Q13627 (STY)** | 50 | 0.903 | 0.903 | 0.909 | 0.903 | 0.931 |  |  |  |
| **Q13882 (STY)** | 26 | 0.821 | 0.778 | 0.757 | 0.821 | 0.521 |  |  |  |
| **Q13976 (STY)** | 84 | 0.915 | 0.909 | 0.917 | 0.915 | 0.918 |  |  |  |
| **Q14012 (STY)** | 42 | 0.924 | 0.917 | 0.924 | 0.924 | 0.873 |  |  |  |
| **Q14164 (STY)** | 48 | 0.823 | 0.79 | 0.789 | 0.823 | 0.734 |  |  |  |
| **Q14289 (STY)** | 20 | 0.817 | 0.765 | 0.725 | 0.817 | 0.645 |  |  |  |
| **Q14680 (STY)** | 29 | 0.798 | 0.751 | 0.718 | 0.798 | 0.642 |  |  |  |
| **Q15046 (STY)** | 38 | 0.882 | 0.864 | 0.86 | 0.882 | 0.891 |  |  |  |
| **Q15118 (STY)** | 31 | 0.897 | 0.885 | 0.882 | 0.897 | 0.924 |  |  |  |
| **Q15139 (STY)** | 90 | 0.926 | 0.917 | 0.927 | 0.926 | 0.917 |  |  |  |
| **Q15418 (STY)** | 109 | 0.925 | 0.921 | 0.924 | 0.925 | 0.933 |  |  |  |
| **Q15759 (STY)** | 51 | 0.938 | 0.937 | 0.94 | 0.938 | 0.942 |  |  |  |
| **Q15831 (STY)** | 37 | 0.883 | 0.851 | 0.862 | 0.883 | 0.738 |  |  |  |
| **Q15835 (STY)** | 19 | 0.823 | 0.807 | 0.809 | 0.823 | 0.753 |  |  |  |
| **Q16512 (STY)** | 16 | 0.836 | 0.804 | 0.786 | 0.836 | 0.912 |  |  |  |
| **Q16539 (STY)** | 301 | 0.936 | 0.935 | 0.935 | 0.936 | 0.95 |  |  |  |
| **Q16566 (STY)** | 24 | 0.91 | 0.889 | 0.893 | 0.91 | 0.858 |  |  |  |
| **Q16620 (STY)** | 17 | 0.834 | 0.777 | 0.737 | 0.834 | 0.783 |  |  |  |
| **Q2MHE4 (STY)** | 40 | 0.636 | 0.622 | 0.66 | 0.636 | 0.729 |  |  |  |
| **Q2TA25 (STY)** | 16 | 0.814 | 0.75 | 0.697 | 0.814 | 0.481 |  |  |  |
| **Q39011 (STY)** | 36 | 0.545 | 0.523 | 0.537 | 0.545 | 0.469 |  |  |  |
| **Q3SYZ2 (STY)** | 16 | 0.827 | 0.789 | 0.762 | 0.827 | 0.694 |  |  |  |
| **Q5EG47 (STY)** | 36 | 0.884 | 0.872 | 0.884 | 0.884 | 0.864 |  |  |  |
| **Q5RCH1 (STY)** | 17 | 0.98 | 0.982 | 0.988 | 0.98 | 0.988 |  |  |  |
| **Q5S007 (STY)** | 94 | 0.847 | 0.808 | 0.819 | 0.847 | 0.696 |  |  |  |
| **Q60670 (STY)** | 15 | 0.9 | 0.879 | 0.881 | 0.9 | 0.952 |  |  |  |
| **Q60680 (STY)** | 18 | 0.833 | 0.8 | 0.786 | 0.833 | 0.706 |  |  |  |
| **Q60737 (STY)** | 68 | 0.892 | 0.883 | 0.884 | 0.892 | 0.915 |  |  |  |
| **Q60806 (STY)** | 16 | 0.834 | 0.782 | 0.742 | 0.834 | 0.619 |  |  |  |
| **Q60823 (STY)** | 21 | 0.889 | 0.879 | 0.886 | 0.889 | 0.935 |  |  |  |
| **Q61036 (STY)** | 17 | 0.884 | 0.836 | 0.798 | 0.884 | 0.648 |  |  |  |
| **Q61831 (STY)** | 20 | 0.925 | 0.922 | 0.934 | 0.925 | 0.965 |  |  |  |
| **Q62101 (STY)** | 19 | 0.93 | 0.918 | 0.919 | 0.93 | 0.903 |  |  |  |
| **Q62120 (STY)** | 48 | 0.792 | 0.75 | 0.724 | 0.792 | 0.524 |  |  |  |
| **Q62388 (STY)** | 49 | 0.932 | 0.933 | 0.94 | 0.932 | 0.921 |  |  |  |
| **Q62689 (STY)** | 22 | 0.842 | 0.798 | 0.774 | 0.842 | 0.712 |  |  |  |
| **Q62844 (STY)** | 22 | 0.842 | 0.789 | 0.751 | 0.842 | 0.673 |  |  |  |
| **Q62868 (STY)** | 20 | 0.833 | 0.791 | 0.758 | 0.833 | 0.655 |  |  |  |
| **Q63450 (STY)** | 18 | 0.9 | 0.88 | 0.873 | 0.9 | 0.872 |  |  |  |
| **Q63470 (STY)** | 23 | 0.833 | 0.804 | 0.785 | 0.833 | 0.677 |  |  |  |
| **Q63531 (STY)** | 27 | 0.908 | 0.894 | 0.901 | 0.908 | 0.9 |  |  |  |
| **Q63644 (STY)** | 17 | 0.845 | 0.821 | 0.811 | 0.845 | 0.779 |  |  |  |
| **Q63699 (STY)** | 20 | 0.942 | 0.943 | 0.954 | 0.942 | 0.9 |  |  |  |
| **Q63844 (STY)** | 107 | 0.952 | 0.952 | 0.953 | 0.952 | 0.939 |  |  |  |
| **Q64303 (STY)** | 15 | 0.844 | 0.797 | 0.762 | 0.844 | 0.53 |  |  |  |
| **Q64702 (STY)** | 16 | 0.867 | 0.817 | 0.78 | 0.867 | 0.75 |  |  |  |
| **Q7KZI7 (STY)** | 31 | 0.877 | 0.856 | 0.853 | 0.877 | 0.706 |  |  |  |
| **Q8BSK8 (STY)** | 29 | 0.919 | 0.91 | 0.924 | 0.919 | 0.895 |  |  |  |
| **Q8C050 (STY)** | 19 | 0.886 | 0.872 | 0.87 | 0.886 | 0.896 |  |  |  |
| **Q8CIN4 (STY)** | 17 | 0.844 | 0.796 | 0.763 | 0.844 | 0.637 |  |  |  |
| **Q8IW41 (STY)** | 18 | 0.882 | 0.865 | 0.863 | 0.882 | 0.85 |  |  |  |
| **Q8IXL6 (STY)** | 196 | 0.893 | 0.883 | 0.889 | 0.893 | 0.855 |  |  |  |
| **Q8N5S9 (STY)** | 15 | 0.9 | 0.869 | 0.849 | 0.9 | 0.771 |  |  |  |
| **Q91Y86 (STY)** | 77 | 0.933 | 0.933 | 0.936 | 0.933 | 0.928 |  |  |  |
| **Q91YS8 (STY)** | 16 | 0.886 | 0.853 | 0.837 | 0.886 | 0.862 |  |  |  |
| **Q92630 (STY)** | 37 | 0.946 | 0.943 | 0.95 | 0.946 | 0.924 |  |  |  |
| **Q94F62 (STY)** | 23 | 0.681 | 0.627 | 0.586 | 0.681 | 0.377 |  |  |  |
| **Q96GD4 (STY)** | 191 | 0.912 | 0.907 | 0.909 | 0.912 | 0.912 |  |  |  |
| **Q96QT4 (STY)** | 63 | 0.811 | 0.762 | 0.749 | 0.811 | 0.57 |  |  |  |
| **Q96SB4 (STY)** | 23 | 0.946 | 0.943 | 0.957 | 0.946 | 0.952 |  |  |  |
| **Q99683 (STY)** | 28 | 0.846 | 0.823 | 0.831 | 0.846 | 0.844 |  |  |  |
| **Q99986 (STY)** | 18 | 0.889 | 0.859 | 0.848 | 0.889 | 0.678 |  |  |  |
| **Q9BXM7 (STY)** | 17 | 0.803 | 0.76 | 0.73 | 0.803 | 0.533 |  |  |  |
| **Q9BZL6 (STY)** | 20 | 0.917 | 0.908 | 0.91 | 0.917 | 0.905 |  |  |  |
| **Q9DC28 (STY)** | 53 | 0.852 | 0.825 | 0.832 | 0.852 | 0.724 |  |  |  |
| **Q9H0K1 (STY)** | 17 | 0.912 | 0.889 | 0.882 | 0.912 | 0.85 |  |  |  |
| **Q9H2X6 (STY)** | 68 | 0.941 | 0.942 | 0.946 | 0.941 | 0.916 |  |  |  |
| **Q9H4B4 (STY)** | 34 | 0.824 | 0.786 | 0.761 | 0.824 | 0.732 |  |  |  |
| **Q9HC98 (STY)** | 23 | 0.827 | 0.805 | 0.79 | 0.827 | 0.806 |  |  |  |
| **Q9JKK8 (STY)** | 15 | 1.0 | 1.0 | 1.0 | 1.0 | 1.0 |  |  |  |
| **Q9JLN9 (STY)** | 104 | 0.899 | 0.892 | 0.893 | 0.899 | 0.876 |  |  |  |
| **Q9NRM7 (STY)** | 19 | 0.939 | 0.935 | 0.941 | 0.939 | 0.989 |  |  |  |
| **Q9NWZ3 (STY)** | 34 | 0.638 | 0.597 | 0.567 | 0.638 | 0.488 |  |  |  |
| **Q9NYY3 (STY)** | 33 | 0.813 | 0.755 | 0.71 | 0.813 | 0.596 |  |  |  |
| **Q9P1W9 (STY)** | 18 | 0.888 | 0.873 | 0.882 | 0.888 | 0.706 |  |  |  |
| **Q9QZR5 (STY)** | 23 | 0.92 | 0.917 | 0.94 | 0.92 | 0.898 |  |  |  |
| **Q9R012 (STY)** | 24 | 0.813 | 0.755 | 0.706 | 0.813 | 0.628 |  |  |  |
| **Q9UBE8 (STY)** | 16 | 0.853 | 0.834 | 0.826 | 0.853 | 0.919 |  |  |  |
| **Q9UEW8 (STY)** | 20 | 0.883 | 0.863 | 0.859 | 0.883 | 0.865 |  |  |  |
| **Q9UHD2 (STY)** | 88 | 0.843 | 0.808 | 0.815 | 0.843 | 0.697 |  |  |  |
| **Q9UM73 (STY)** | 15 | 0.778 | 0.727 | 0.69 | 0.778 | 0.686 |  |  |  |
| **Q9UQM7 (STY)** | 279 | 0.878 | 0.866 | 0.867 | 0.878 | 0.883 |  |  |  |
| **Q9WTK7 (STY)** | 17 | 0.931 | 0.91 | 0.902 | 0.931 | 0.801 |  |  |  |
| **Q9WTU6 (STY)** | 30 | 0.939 | 0.937 | 0.944 | 0.939 | 0.956 |  |  |  |
| **Q9WUD9 (STY)** | 76 | 0.838 | 0.806 | 0.802 | 0.838 | 0.72 |  |  |  |
| **Q9WV60 (STY)** | 104 | 0.901 | 0.897 | 0.9 | 0.901 | 0.906 |  |  |  |
| **Q9WVC6 (STY)** | 32 | 0.933 | 0.928 | 0.941 | 0.933 | 0.976 |  |  |  |
| **Q9Y478 (STY)** | 57 | 0.889 | 0.879 | 0.879 | 0.889 | 0.876 |  |  |  |
| **Q9Z2A0 (STY)** | 35 | 0.924 | 0.906 | 0.895 | 0.924 | 0.906 |  |  |  |
| **A6QLB8 (ST)** | 16 | 0.775 | 0.758 | 0.777 | 0.775 | 0.85 |  |  |  |
| **G3N1T2 (ST)** | 24 | 0.869 | 0.846 | 0.851 | 0.869 | 0.775 |  |  |  |
| **O00141 (ST)** | 76 | 0.954 | 0.953 | 0.955 | 0.954 | 0.967 |  |  |  |
| **O00311 (ST)** | 48 | 0.844 | 0.815 | 0.825 | 0.844 | 0.738 |  |  |  |
| **O00418 (ST)** | 16 | 0.843 | 0.824 | 0.826 | 0.843 | 0.797 |  |  |  |
| **O14757 (ST)** | 201 | 0.894 | 0.886 | 0.886 | 0.894 | 0.854 |  |  |  |
| **O14920 (ST)** | 96 | 0.826 | 0.789 | 0.797 | 0.826 | 0.724 |  |  |  |
| **O14965 (ST)** | 144 | 0.895 | 0.887 | 0.889 | 0.895 | 0.849 |  |  |  |
| **O15111 (ST)** | 56 | 0.822 | 0.777 | 0.756 | 0.822 | 0.665 |  |  |  |
| **O15264 (ST)** | 38 | 0.935 | 0.929 | 0.926 | 0.935 | 0.909 |  |  |  |
| **O15530 (ST)** | 76 | 0.908 | 0.898 | 0.901 | 0.908 | 0.906 |  |  |  |
| **O43293 (ST)** | 24 | 0.876 | 0.85 | 0.836 | 0.876 | 0.788 |  |  |  |
| **O43318 (ST)** | 43 | 0.872 | 0.848 | 0.846 | 0.872 | 0.796 |  |  |  |
| **O43683 (ST)** | 27 | 0.673 | 0.631 | 0.615 | 0.673 | 0.537 |  |  |  |
| **O55099 (ST)** | 18 | 0.86 | 0.858 | 0.884 | 0.86 | 0.822 |  |  |  |
| **O55173 (ST)** | 31 | 0.945 | 0.931 | 0.936 | 0.945 | 0.958 |  |  |  |
| **O70126 (ST)** | 34 | 0.887 | 0.877 | 0.894 | 0.887 | 0.901 |  |  |  |
| **O70405 (ST)** | 17 | 0.795 | 0.738 | 0.692 | 0.795 | 0.569 |  |  |  |
| **O75116 (ST)** | 48 | 0.882 | 0.86 | 0.858 | 0.882 | 0.803 |  |  |  |
| **O75385 (ST)** | 80 | 0.838 | 0.805 | 0.795 | 0.838 | 0.706 |  |  |  |
| **O75582 (ST)** | 33 | 0.894 | 0.883 | 0.879 | 0.894 | 0.877 |  |  |  |
| **O88351 (ST)** | 39 | 0.85 | 0.812 | 0.801 | 0.85 | 0.644 |  |  |  |
| **O88643 (ST)** | 22 | 0.84 | 0.798 | 0.763 | 0.84 | 0.677 |  |  |  |
| **O95747 (ST)** | 15 | 0.9 | 0.889 | 0.884 | 0.9 | 0.846 |  |  |  |
| **O95835 (ST)** | 17 | 0.931 | 0.926 | 0.926 | 0.931 | 0.761 |  |  |  |
| **O96013 (ST)** | 21 | 0.856 | 0.826 | 0.811 | 0.856 | 0.843 |  |  |  |
| **O96017 (ST)** | 74 | 0.872 | 0.851 | 0.861 | 0.872 | 0.839 |  |  |  |
| **P00516 (ST)** | 54 | 0.92 | 0.908 | 0.923 | 0.92 | 0.855 |  |  |  |
| **P00517 (ST)** | 216 | 0.92 | 0.917 | 0.916 | 0.92 | 0.943 |  |  |  |
| **P00546 (ST)** | 93 | 0.836 | 0.814 | 0.823 | 0.836 | 0.765 |  |  |  |
| **P04049 (ST)** | 20 | 0.85 | 0.815 | 0.797 | 0.85 | 0.745 |  |  |  |
| **P04409 (ST)** | 59 | 0.87 | 0.853 | 0.856 | 0.87 | 0.897 |  |  |  |
| **P04551 (ST)** | 38 | 0.965 | 0.966 | 0.97 | 0.965 | 0.997 |  |  |  |
| **P05129 (ST)** | 56 | 0.863 | 0.839 | 0.857 | 0.863 | 0.812 |  |  |  |
| **P05132 (ST)** | 193 | 0.938 | 0.935 | 0.937 | 0.938 | 0.942 |  |  |  |
| **P05696 (ST)** | 187 | 0.878 | 0.865 | 0.867 | 0.878 | 0.863 |  |  |  |
| **P05771 (ST)** | 101 | 0.851 | 0.832 | 0.834 | 0.851 | 0.769 |  |  |  |
| **P06493 (ST)** | 760 | 0.962 | 0.962 | 0.964 | 0.962 | 0.975 |  |  |  |
| **P08413 (ST)** | 64 | 0.828 | 0.794 | 0.778 | 0.828 | 0.748 |  |  |  |
| **P09215 (ST)** | 23 | 0.848 | 0.817 | 0.807 | 0.848 | 0.739 |  |  |  |
| **P09216 (ST)** | 15 | 0.778 | 0.737 | 0.72 | 0.778 | 0.648 |  |  |  |
| **P09217 (ST)** | 22 | 0.819 | 0.768 | 0.732 | 0.819 | 0.752 |  |  |  |
| **P0C605 (ST)** | 26 | 0.93 | 0.927 | 0.944 | 0.93 | 0.915 |  |  |  |
| **P11275 (ST)** | 104 | 0.875 | 0.864 | 0.865 | 0.875 | 0.856 |  |  |  |
| **P11309 (ST)** | 64 | 0.867 | 0.853 | 0.856 | 0.867 | 0.783 |  |  |  |
| **P11440 (ST)** | 135 | 0.968 | 0.968 | 0.969 | 0.968 | 0.966 |  |  |  |
| **P11798 (ST)** | 73 | 0.884 | 0.866 | 0.879 | 0.884 | 0.812 |  |  |  |
| **P11802 (ST)** | 77 | 0.946 | 0.949 | 0.958 | 0.946 | 0.964 |  |  |  |
| **P13234 (ST)** | 17 | 0.843 | 0.821 | 0.821 | 0.843 | 0.843 |  |  |  |
| **P16054 (ST)** | 28 | 0.803 | 0.77 | 0.752 | 0.803 | 0.613 |  |  |  |
| **P17157 (ST)** | 20 | 0.892 | 0.886 | 0.899 | 0.892 | 0.97 |  |  |  |
| **P17252 (ST)** | 718 | 0.862 | 0.86 | 0.86 | 0.862 | 0.908 |  |  |  |
| **P17612 (ST)** | 929 | 0.915 | 0.914 | 0.916 | 0.915 | 0.959 |  |  |  |
| **P18265 (ST)** | 20 | 0.875 | 0.853 | 0.849 | 0.875 | 0.86 |  |  |  |
| **P18266 (ST)** | 62 | 0.914 | 0.903 | 0.914 | 0.914 | 0.893 |  |  |  |
| **P18653 (ST)** | 36 | 0.926 | 0.921 | 0.93 | 0.926 | 0.907 |  |  |  |
| **P18654 (ST)** | 25 | 0.88 | 0.875 | 0.888 | 0.88 | 0.862 |  |  |  |
| **P19139 (ST)** | 75 | 0.922 | 0.916 | 0.923 | 0.922 | 0.909 |  |  |  |
| **P19525 (ST)** | 21 | 0.795 | 0.759 | 0.735 | 0.795 | 0.571 |  |  |  |
| **P19784 (ST)** | 21 | 0.898 | 0.88 | 0.872 | 0.898 | 0.914 |  |  |  |
| **P20444 (ST)** | 103 | 0.874 | 0.853 | 0.868 | 0.874 | 0.843 |  |  |  |
| **P21708 (ST)** | 67 | 0.948 | 0.948 | 0.953 | 0.948 | 0.959 |  |  |  |
| **P23443 (ST)** | 59 | 0.921 | 0.915 | 0.921 | 0.921 | 0.932 |  |  |  |
| **P24723 (ST)** | 18 | 0.825 | 0.78 | 0.748 | 0.825 | 0.644 |  |  |  |
| **P24941 (ST)** | 465 | 0.98 | 0.979 | 0.98 | 0.98 | 0.982 |  |  |  |
| **P25098 (ST)** | 118 | 0.857 | 0.839 | 0.84 | 0.857 | 0.841 |  |  |  |
| **P26927 (ST)** | 15 | 0.844 | 0.805 | 0.772 | 0.844 | 0.55 |  |  |  |
| **P27361 (ST)** | 484 | 0.963 | 0.962 | 0.963 | 0.963 | 0.975 |  |  |  |
| **P27791 (ST)** | 143 | 0.923 | 0.917 | 0.921 | 0.923 | 0.924 |  |  |  |
| **P28482 (ST)** | 597 | 0.968 | 0.968 | 0.968 | 0.968 | 0.98 |  |  |  |
| **P28867 (ST)** | 29 | 0.845 | 0.821 | 0.815 | 0.845 | 0.843 |  |  |  |
| **P31749 (ST)** | 411 | 0.946 | 0.945 | 0.945 | 0.946 | 0.974 |  |  |  |
| **P31750 (ST)** | 101 | 0.955 | 0.954 | 0.957 | 0.955 | 0.966 |  |  |  |
| **P31751 (ST)** | 79 | 0.903 | 0.894 | 0.902 | 0.903 | 0.907 |  |  |  |
| **P32562 (ST)** | 18 | 0.76 | 0.721 | 0.695 | 0.76 | 0.735 |  |  |  |
| **P33674 (ST)** | 18 | 0.955 | 0.953 | 0.959 | 0.955 | 0.989 |  |  |  |
| **P33981 (ST)** | 65 | 0.826 | 0.775 | 0.752 | 0.826 | 0.669 |  |  |  |
| **P34947 (ST)** | 23 | 0.813 | 0.784 | 0.773 | 0.813 | 0.686 |  |  |  |
| **P35465 (ST)** | 25 | 0.893 | 0.878 | 0.874 | 0.893 | 0.821 |  |  |  |
| **P35626 (ST)** | 27 | 0.846 | 0.817 | 0.81 | 0.846 | 0.753 |  |  |  |
| **P36887 (ST)** | 20 | 0.842 | 0.811 | 0.792 | 0.842 | 0.91 |  |  |  |
| **P38110 (ST)** | 19 | 0.982 | 0.983 | 0.988 | 0.982 | 0.989 |  |  |  |
| **P38111 (ST)** | 21 | 0.976 | 0.978 | 0.984 | 0.976 | 0.995 |  |  |  |
| **P39951 (ST)** | 79 | 0.947 | 0.947 | 0.949 | 0.947 | 0.983 |  |  |  |
| **P41279 (ST)** | 25 | 0.853 | 0.817 | 0.8 | 0.853 | 0.745 |  |  |  |
| **P41743 (ST)** | 47 | 0.837 | 0.805 | 0.814 | 0.837 | 0.759 |  |  |  |
| **P42345 (ST)** | 142 | 0.872 | 0.858 | 0.86 | 0.872 | 0.842 |  |  |  |
| **P42346 (ST)** | 17 | 0.832 | 0.793 | 0.772 | 0.832 | 0.7 |  |  |  |
| **P43250 (ST)** | 16 | 0.827 | 0.787 | 0.753 | 0.827 | 0.625 |  |  |  |
| **P45983 (ST)** | 229 | 0.974 | 0.974 | 0.975 | 0.974 | 0.977 |  |  |  |
| **P45984 (ST)** | 82 | 0.955 | 0.954 | 0.956 | 0.955 | 0.971 |  |  |  |
| **P46196 (ST)** | 17 | 0.795 | 0.783 | 0.785 | 0.795 | 0.797 |  |  |  |
| **P47196 (ST)** | 49 | 0.946 | 0.946 | 0.951 | 0.946 | 0.98 |  |  |  |
| **P47811 (ST)** | 89 | 0.914 | 0.91 | 0.914 | 0.914 | 0.927 |  |  |  |
| **P48729 (ST)** | 169 | 0.895 | 0.882 | 0.894 | 0.895 | 0.881 |  |  |  |
| **P48730 (ST)** | 105 | 0.879 | 0.862 | 0.882 | 0.879 | 0.848 |  |  |  |
| **P48734 (ST)** | 36 | 0.968 | 0.968 | 0.973 | 0.968 | 0.987 |  |  |  |
| **P49137 (ST)** | 101 | 0.926 | 0.921 | 0.929 | 0.926 | 0.91 |  |  |  |
| **P49138 (ST)** | 36 | 0.907 | 0.896 | 0.915 | 0.907 | 0.894 |  |  |  |
| **P49185 (ST)** | 42 | 0.917 | 0.914 | 0.921 | 0.917 | 0.957 |  |  |  |
| **P49186 (ST)** | 16 | 0.938 | 0.928 | 0.929 | 0.938 | 0.912 |  |  |  |
| **P49336 (ST)** | 18 | 0.899 | 0.892 | 0.9 | 0.899 | 0.867 |  |  |  |
| **P49615 (ST)** | 105 | 0.959 | 0.959 | 0.961 | 0.959 | 0.948 |  |  |  |
| **P49674 (ST)** | 60 | 0.856 | 0.833 | 0.839 | 0.856 | 0.856 |  |  |  |
| **P49760 (ST)** | 20 | 0.782 | 0.742 | 0.715 | 0.782 | 0.876 |  |  |  |
| **P49840 (ST)** | 109 | 0.959 | 0.958 | 0.959 | 0.959 | 0.963 |  |  |  |
| **P49841 (ST)** | 432 | 0.931 | 0.929 | 0.93 | 0.931 | 0.963 |  |  |  |
| **P50613 (ST)** | 59 | 0.864 | 0.849 | 0.851 | 0.864 | 0.751 |  |  |  |
| **P50750 (ST)** | 51 | 0.905 | 0.893 | 0.891 | 0.905 | 0.853 |  |  |  |
| **P51812 (ST)** | 72 | 0.914 | 0.904 | 0.919 | 0.914 | 0.882 |  |  |  |
| **P51955 (ST)** | 35 | 0.771 | 0.735 | 0.704 | 0.771 | 0.572 |  |  |  |
| **P53350 (ST)** | 340 | 0.851 | 0.835 | 0.844 | 0.851 | 0.836 |  |  |  |
| **P53351 (ST)** | 24 | 0.786 | 0.741 | 0.701 | 0.786 | 0.474 |  |  |  |
| **P53355 (ST)** | 31 | 0.844 | 0.819 | 0.804 | 0.844 | 0.695 |  |  |  |
| **P53778 (ST)** | 41 | 0.931 | 0.928 | 0.935 | 0.931 | 0.961 |  |  |  |
| **P53779 (ST)** | 32 | 0.917 | 0.913 | 0.923 | 0.917 | 0.894 |  |  |  |
| **P54199 (ST)** | 18 | 0.762 | 0.695 | 0.647 | 0.762 | 0.504 |  |  |  |
| **P54645 (ST)** | 30 | 0.861 | 0.852 | 0.862 | 0.861 | 0.882 |  |  |  |
| **P54646 (ST)** | 47 | 0.876 | 0.854 | 0.866 | 0.876 | 0.837 |  |  |  |
| **P57059 (ST)** | 20 | 0.883 | 0.861 | 0.851 | 0.883 | 0.795 |  |  |  |
| **P63085 (ST)** | 265 | 0.97 | 0.97 | 0.971 | 0.97 | 0.981 |  |  |  |
| **P63086 (ST)** | 89 | 0.949 | 0.95 | 0.951 | 0.949 | 0.95 |  |  |  |
| **P67870 (ST)** | 20 | 0.908 | 0.9 | 0.904 | 0.908 | 0.895 |  |  |  |
| **P67999 (ST)** | 21 | 0.937 | 0.928 | 0.945 | 0.937 | 0.92 |  |  |  |
| **P68399 (ST)** | 16 | 0.94 | 0.937 | 0.942 | 0.94 | 0.969 |  |  |  |
| **P68400 (ST)** | 742 | 0.921 | 0.919 | 0.921 | 0.921 | 0.961 |  |  |  |
| **P68403 (ST)** | 25 | 0.807 | 0.774 | 0.752 | 0.807 | 0.691 |  |  |  |
| **P68404 (ST)** | 22 | 0.858 | 0.825 | 0.808 | 0.858 | 0.847 |  |  |  |
| **P70032 (ST)** | 15 | 0.811 | 0.757 | 0.712 | 0.811 | 0.529 |  |  |  |
| **P70335 (ST)** | 27 | 0.845 | 0.826 | 0.819 | 0.845 | 0.738 |  |  |  |
| **P70336 (ST)** | 27 | 0.865 | 0.828 | 0.816 | 0.865 | 0.742 |  |  |  |
| **P70618 (ST)** | 21 | 0.906 | 0.901 | 0.915 | 0.906 | 0.954 |  |  |  |
| **P78527 (ST)** | 136 | 0.911 | 0.904 | 0.909 | 0.911 | 0.899 |  |  |  |
| **P97313 (ST)** | 17 | 0.833 | 0.782 | 0.747 | 0.833 | 0.794 |  |  |  |
| **P97377 (ST)** | 72 | 0.935 | 0.936 | 0.939 | 0.935 | 0.956 |  |  |  |
| **P97633 (ST)** | 15 | 0.8 | 0.751 | 0.713 | 0.8 | 0.648 |  |  |  |
| **Q00526 (ST)** | 16 | 0.917 | 0.916 | 0.928 | 0.917 | 0.944 |  |  |  |
| **Q00534 (ST)** | 41 | 0.942 | 0.944 | 0.955 | 0.942 | 0.967 |  |  |  |
| **Q00535 (ST)** | 246 | 0.97 | 0.97 | 0.971 | 0.97 | 0.969 |  |  |  |
| **Q01314 (ST)** | 20 | 0.925 | 0.918 | 0.923 | 0.925 | 0.975 |  |  |  |
| **Q02111 (ST)** | 17 | 0.854 | 0.805 | 0.767 | 0.854 | 0.591 |  |  |  |
| **Q02156 (ST)** | 113 | 0.875 | 0.86 | 0.866 | 0.875 | 0.809 |  |  |  |
| **Q02399 (ST)** | 16 | 0.896 | 0.892 | 0.913 | 0.896 | 0.931 |  |  |  |
| **Q02956 (ST)** | 34 | 0.868 | 0.843 | 0.86 | 0.868 | 0.8 |  |  |  |
| **Q03114 (ST)** | 73 | 0.97 | 0.971 | 0.976 | 0.97 | 0.979 |  |  |  |
| **Q04759 (ST)** | 57 | 0.903 | 0.892 | 0.899 | 0.903 | 0.817 |  |  |  |
| **Q05513 (ST)** | 121 | 0.842 | 0.806 | 0.816 | 0.842 | 0.713 |  |  |  |
| **Q05655 (ST)** | 175 | 0.867 | 0.849 | 0.859 | 0.867 | 0.834 |  |  |  |
| **Q06226 (ST)** | 15 | 0.889 | 0.852 | 0.826 | 0.889 | 0.777 |  |  |  |
| **Q07832 (ST)** | 49 | 0.847 | 0.808 | 0.79 | 0.847 | 0.744 |  |  |  |
| **Q09137 (ST)** | 35 | 0.876 | 0.857 | 0.883 | 0.876 | 0.884 |  |  |  |
| **Q13043 (ST)** | 34 | 0.844 | 0.811 | 0.797 | 0.844 | 0.754 |  |  |  |
| **Q13131 (ST)** | 203 | 0.904 | 0.896 | 0.898 | 0.904 | 0.896 |  |  |  |
| **Q13153 (ST)** | 91 | 0.861 | 0.85 | 0.849 | 0.861 | 0.855 |  |  |  |
| **Q13164 (ST)** | 32 | 0.87 | 0.855 | 0.87 | 0.87 | 0.894 |  |  |  |
| **Q13177 (ST)** | 50 | 0.843 | 0.816 | 0.821 | 0.843 | 0.711 |  |  |  |
| **Q13188 (ST)** | 28 | 0.834 | 0.8 | 0.779 | 0.834 | 0.67 |  |  |  |
| **Q13237 (ST)** | 29 | 0.886 | 0.876 | 0.89 | 0.886 | 0.943 |  |  |  |
| **Q13315 (ST)** | 321 | 0.968 | 0.968 | 0.968 | 0.968 | 0.978 |  |  |  |
| **Q13464 (ST)** | 86 | 0.888 | 0.879 | 0.89 | 0.888 | 0.881 |  |  |  |
| **Q13535 (ST)** | 124 | 0.965 | 0.965 | 0.967 | 0.965 | 0.965 |  |  |  |
| **Q13554 (ST)** | 22 | 0.857 | 0.844 | 0.843 | 0.857 | 0.821 |  |  |  |
| **Q13557 (ST)** | 26 | 0.891 | 0.873 | 0.876 | 0.891 | 0.774 |  |  |  |
| **Q13627 (ST)** | 49 | 0.928 | 0.929 | 0.935 | 0.928 | 0.949 |  |  |  |
| **Q13976 (ST)** | 84 | 0.941 | 0.934 | 0.944 | 0.941 | 0.937 |  |  |  |
| **Q14012 (ST)** | 42 | 0.904 | 0.899 | 0.906 | 0.904 | 0.877 |  |  |  |
| **Q14164 (ST)** | 48 | 0.84 | 0.815 | 0.818 | 0.84 | 0.788 |  |  |  |
| **Q14680 (ST)** | 26 | 0.815 | 0.758 | 0.716 | 0.815 | 0.506 |  |  |  |
| **Q15118 (ST)** | 31 | 0.898 | 0.886 | 0.898 | 0.898 | 0.932 |  |  |  |
| **Q15139 (ST)** | 90 | 0.93 | 0.923 | 0.933 | 0.93 | 0.905 |  |  |  |
| **Q15418 (ST)** | 109 | 0.919 | 0.915 | 0.919 | 0.919 | 0.914 |  |  |  |
| **Q15759 (ST)** | 51 | 0.935 | 0.931 | 0.94 | 0.935 | 0.945 |  |  |  |
| **Q15831 (ST)** | 37 | 0.87 | 0.845 | 0.861 | 0.87 | 0.694 |  |  |  |
| **Q15835 (ST)** | 19 | 0.843 | 0.802 | 0.775 | 0.843 | 0.79 |  |  |  |
| **Q16512 (ST)** | 16 | 0.814 | 0.791 | 0.782 | 0.814 | 0.825 |  |  |  |
| **Q16539 (ST)** | 300 | 0.935 | 0.934 | 0.934 | 0.935 | 0.959 |  |  |  |
| **Q16566 (ST)** | 24 | 0.881 | 0.857 | 0.848 | 0.881 | 0.844 |  |  |  |
| **Q2MHE4 (ST)** | 38 | 0.624 | 0.599 | 0.62 | 0.624 | 0.772 |  |  |  |
| **Q2TA25 (ST)** | 16 | 0.791 | 0.735 | 0.694 | 0.791 | 0.425 |  |  |  |
| **Q39011 (ST)** | 36 | 0.568 | 0.539 | 0.544 | 0.568 | 0.534 |  |  |  |
| **Q3SYZ2 (ST)** | 16 | 0.762 | 0.722 | 0.689 | 0.762 | 0.606 |  |  |  |
| **Q5EG47 (ST)** | 36 | 0.912 | 0.906 | 0.91 | 0.912 | 0.912 |  |  |  |
| **Q5RCH1 (ST)** | 17 | 0.97 | 0.973 | 0.983 | 0.97 | 0.989 |  |  |  |
| **Q5S007 (ST)** | 91 | 0.846 | 0.809 | 0.808 | 0.846 | 0.696 |  |  |  |
| **Q60670 (ST)** | 15 | 0.9 | 0.875 | 0.862 | 0.9 | 0.907 |  |  |  |
| **Q60680 (ST)** | 18 | 0.835 | 0.8 | 0.777 | 0.835 | 0.672 |  |  |  |
| **Q60737 (ST)** | 68 | 0.88 | 0.871 | 0.876 | 0.88 | 0.902 |  |  |  |
| **Q60806 (ST)** | 16 | 0.803 | 0.744 | 0.696 | 0.803 | 0.506 |  |  |  |
| **Q60823 (ST)** | 21 | 0.976 | 0.974 | 0.98 | 0.976 | 0.976 |  |  |  |
| **Q61036 (ST)** | 17 | 0.885 | 0.849 | 0.824 | 0.885 | 0.642 |  |  |  |
| **Q61831 (ST)** | 20 | 0.917 | 0.913 | 0.916 | 0.917 | 0.96 |  |  |  |
| **Q62101 (ST)** | 19 | 0.938 | 0.928 | 0.937 | 0.938 | 0.838 |  |  |  |
| **Q62388 (ST)** | 49 | 0.959 | 0.96 | 0.963 | 0.959 | 0.944 |  |  |  |
| **Q62868 (ST)** | 20 | 0.85 | 0.799 | 0.764 | 0.85 | 0.535 |  |  |  |
| **Q63450 (ST)** | 18 | 0.899 | 0.879 | 0.871 | 0.899 | 0.822 |  |  |  |
| **Q63470 (ST)** | 19 | 0.859 | 0.836 | 0.82 | 0.859 | 0.834 |  |  |  |
| **Q63531 (ST)** | 27 | 0.913 | 0.903 | 0.917 | 0.913 | 0.908 |  |  |  |
| **Q63644 (ST)** | 17 | 0.882 | 0.856 | 0.84 | 0.882 | 0.815 |  |  |  |
| **Q63699 (ST)** | 20 | 0.942 | 0.942 | 0.953 | 0.942 | 0.925 |  |  |  |
| **Q63844 (ST)** | 106 | 0.958 | 0.957 | 0.958 | 0.958 | 0.952 |  |  |  |
| **Q64303 (ST)** | 15 | 0.811 | 0.78 | 0.764 | 0.811 | 0.491 |  |  |  |
| **Q64702 (ST)** | 16 | 0.874 | 0.84 | 0.817 | 0.874 | 0.881 |  |  |  |
| **Q7KZI7 (ST)** | 31 | 0.871 | 0.856 | 0.86 | 0.871 | 0.707 |  |  |  |
| **Q8BSK8 (ST)** | 29 | 0.919 | 0.911 | 0.924 | 0.919 | 0.859 |  |  |  |
| **Q8C050 (ST)** | 19 | 0.902 | 0.887 | 0.89 | 0.902 | 0.843 |  |  |  |
| **Q8CIN4 (ST)** | 17 | 0.823 | 0.794 | 0.776 | 0.823 | 0.636 |  |  |  |
| **Q8IW41 (ST)** | 18 | 0.909 | 0.885 | 0.885 | 0.909 | 0.828 |  |  |  |
| **Q8IXL6 (ST)** | 196 | 0.896 | 0.89 | 0.89 | 0.896 | 0.878 |  |  |  |
| **Q8N5S9 (ST)** | 15 | 0.9 | 0.872 | 0.861 | 0.9 | 0.938 |  |  |  |
| **Q91Y86 (ST)** | 76 | 0.958 | 0.959 | 0.962 | 0.958 | 0.961 |  |  |  |
| **Q91YS8 (ST)** | 16 | 0.909 | 0.883 | 0.869 | 0.909 | 0.85 |  |  |  |
| **Q92630 (ST)** | 37 | 0.91 | 0.908 | 0.914 | 0.91 | 0.91 |  |  |  |
| **Q94F62 (ST)** | 23 | 0.738 | 0.691 | 0.684 | 0.738 | 0.542 |  |  |  |
| **Q96GD4 (ST)** | 190 | 0.918 | 0.914 | 0.916 | 0.918 | 0.914 |  |  |  |
| **Q96QT4 (ST)** | 63 | 0.749 | 0.72 | 0.704 | 0.749 | 0.523 |  |  |  |
| **Q96SB4 (ST)** | 23 | 0.922 | 0.911 | 0.912 | 0.922 | 0.944 |  |  |  |
| **Q99683 (ST)** | 26 | 0.835 | 0.799 | 0.78 | 0.835 | 0.723 |  |  |  |
| **Q99986 (ST)** | 18 | 0.899 | 0.865 | 0.849 | 0.899 | 0.744 |  |  |  |
| **Q9BXM7 (ST)** | 17 | 0.862 | 0.818 | 0.792 | 0.862 | 0.588 |  |  |  |
| **Q9BZL6 (ST)** | 20 | 0.933 | 0.926 | 0.925 | 0.933 | 0.945 |  |  |  |
| **Q9DC28 (ST)** | 53 | 0.819 | 0.784 | 0.796 | 0.819 | 0.75 |  |  |  |
| **Q9H0K1 (ST)** | 17 | 0.912 | 0.893 | 0.886 | 0.912 | 0.835 |  |  |  |
| **Q9H2X6 (ST)** | 67 | 0.925 | 0.924 | 0.929 | 0.925 | 0.926 |  |  |  |
| **Q9H4B4 (ST)** | 34 | 0.83 | 0.793 | 0.765 | 0.83 | 0.79 |  |  |  |
| **Q9HC98 (ST)** | 22 | 0.894 | 0.88 | 0.886 | 0.894 | 0.915 |  |  |  |
| **Q9JKK8 (ST)** | 15 | 0.933 | 0.931 | 0.943 | 0.933 | 0.966 |  |  |  |
| **Q9JLN9 (ST)** | 104 | 0.907 | 0.901 | 0.902 | 0.907 | 0.897 |  |  |  |
| **Q9NRM7 (ST)** | 19 | 0.964 | 0.954 | 0.951 | 0.964 | 0.995 |  |  |  |
| **Q9NWZ3 (ST)** | 34 | 0.651 | 0.618 | 0.609 | 0.651 | 0.557 |  |  |  |
| **Q9NYY3 (ST)** | 33 | 0.818 | 0.761 | 0.715 | 0.818 | 0.654 |  |  |  |
| **Q9P1W9 (ST)** | 18 | 0.898 | 0.878 | 0.871 | 0.898 | 0.7 |  |  |  |
| **Q9QZR5 (ST)** | 22 | 0.88 | 0.873 | 0.892 | 0.88 | 0.93 |  |  |  |
| **Q9R012 (ST)** | 24 | 0.813 | 0.764 | 0.727 | 0.813 | 0.533 |  |  |  |
| **Q9UBE8 (ST)** | 16 | 0.866 | 0.847 | 0.845 | 0.866 | 0.938 |  |  |  |
| **Q9UEW8 (ST)** | 20 | 0.875 | 0.846 | 0.832 | 0.875 | 0.855 |  |  |  |
| **Q9UHD2 (ST)** | 88 | 0.856 | 0.821 | 0.832 | 0.856 | 0.737 |  |  |  |
| **Q9UQM7 (ST)** | 279 | 0.873 | 0.863 | 0.865 | 0.873 | 0.884 |  |  |  |
| **Q9WTK7 (ST)** | 17 | 0.921 | 0.907 | 0.903 | 0.921 | 0.762 |  |  |  |
| **Q9WTU6 (ST)** | 30 | 0.9 | 0.894 | 0.908 | 0.9 | 0.938 |  |  |  |
| **Q9WV60 (ST)** | 103 | 0.909 | 0.905 | 0.906 | 0.909 | 0.893 |  |  |  |
| **Q9WVC6 (ST)** | 32 | 0.948 | 0.942 | 0.954 | 0.948 | 0.981 |  |  |  |
| **Q9Y478 (ST)** | 57 | 0.918 | 0.905 | 0.906 | 0.918 | 0.891 |  |  |  |
| **Q9Z2A0 (ST)** | 35 | 0.929 | 0.912 | 0.905 | 0.929 | 0.951 |  |  |  |
| **A9UF07 (Y)** | 18 | 0.78 | 0.736 | 0.7 | 0.78 | 0.683 |  |  |  |
| **O60674 (Y)** | 59 | 0.815 | 0.773 | 0.773 | 0.815 | 0.706 |  |  |  |
| **P00519 (Y)** | 225 | 0.766 | 0.758 | 0.764 | 0.766 | 0.804 |  |  |  |
| **P00520 (Y)** | 82 | 0.802 | 0.779 | 0.784 | 0.802 | 0.735 |  |  |  |
| **P00523 (Y)** | 47 | 0.867 | 0.849 | 0.872 | 0.867 | 0.844 |  |  |  |
| **P00533 (Y)** | 116 | 0.796 | 0.772 | 0.782 | 0.796 | 0.736 |  |  |  |
| **P04626 (Y)** | 17 | 0.834 | 0.765 | 0.711 | 0.834 | 0.691 |  |  |  |
| **P04629 (Y)** | 15 | 0.78 | 0.75 | 0.767 | 0.78 | 0.733 |  |  |  |
| **P05480 (Y)** | 157 | 0.813 | 0.802 | 0.805 | 0.813 | 0.841 |  |  |  |
| **P06213 (Y)** | 71 | 0.806 | 0.77 | 0.772 | 0.806 | 0.763 |  |  |  |
| **P06239 (Y)** | 134 | 0.8 | 0.792 | 0.797 | 0.8 | 0.814 |  |  |  |
| **P06240 (Y)** | 34 | 0.881 | 0.859 | 0.873 | 0.881 | 0.861 |  |  |  |
| **P06241 (Y)** | 206 | 0.786 | 0.778 | 0.78 | 0.786 | 0.833 |  |  |  |
| **P07947 (Y)** | 25 | 0.773 | 0.736 | 0.707 | 0.773 | 0.593 |  |  |  |
| **P07948 (Y)** | 143 | 0.749 | 0.732 | 0.744 | 0.749 | 0.73 |  |  |  |
| P07949 (Y) | 27 | 0.814 | 0.772 | 0.745 | 0.814 | 0.682 |  |  |  |
| P08069 (Y) | 28 | 0.786 | 0.756 | 0.748 | 0.786 | 0.612 |  |  |  |
| P08103 (Y) | 15 | 0.8 | 0.751 | 0.71 | 0.8 | 0.709 |  |  |  |
| P08581 (Y) | 36 | 0.799 | 0.774 | 0.779 | 0.799 | 0.839 |  |  |  |
| P08631 (Y) | 36 | 0.81 | 0.796 | 0.811 | 0.81 | 0.713 |  |  |  |
| P09619 (Y) | 34 | 0.834 | 0.807 | 0.795 | 0.834 | 0.74 |  |  |  |
| P09769 (Y) | 19 | 0.8 | 0.754 | 0.718 | 0.8 | 0.724 |  |  |  |
| P11362 (Y) | 46 | 0.777 | 0.722 | 0.693 | 0.777 | 0.577 |  |  |  |
| P12931 (Y) | 652 | 0.762 | 0.761 | 0.765 | 0.762 | 0.838 |  |  |  |
| P15127 (Y) | 27 | 0.83 | 0.779 | 0.759 | 0.83 | 0.733 |  |  |  |
| P15208 (Y) | 26 | 0.842 | 0.821 | 0.827 | 0.842 | 0.793 |  |  |  |
| P16092 (Y) | 18 | 0.757 | 0.708 | 0.677 | 0.757 | 0.611 |  |  |  |
| P16591 (Y) | 21 | 0.801 | 0.759 | 0.729 | 0.801 | 0.553 |  |  |  |
| P23458 (Y) | 17 | 0.755 | 0.724 | 0.698 | 0.755 | 0.585 |  |  |  |
| P25911 (Y) | 51 | 0.831 | 0.815 | 0.827 | 0.831 | 0.856 |  |  |  |
| P29597 (Y) | 17 | 0.744 | 0.691 | 0.65 | 0.744 | 0.504 |  |  |  |
| P32577 (Y) | 22 | 0.823 | 0.769 | 0.736 | 0.823 | 0.72 |  |  |  |
| P39688 (Y) | 65 | 0.815 | 0.788 | 0.783 | 0.815 | 0.764 |  |  |  |
| P41240 (Y) | 28 | 0.85 | 0.827 | 0.839 | 0.85 | 0.662 |  |  |  |
| P41241 (Y) | 22 | 0.816 | 0.776 | 0.751 | 0.816 | 0.635 |  |  |  |
| P42684 (Y) | 33 | 0.851 | 0.835 | 0.831 | 0.851 | 0.739 |  |  |  |
| P43403 (Y) | 39 | 0.77 | 0.744 | 0.767 | 0.77 | 0.715 |  |  |  |
| P43405 (Y) | 82 | 0.858 | 0.848 | 0.859 | 0.858 | 0.882 |  |  |  |
| P48025 (Y) | 36 | 0.846 | 0.814 | 0.809 | 0.846 | 0.841 |  |  |  |
| P51813 (Y) | 22 | 0.857 | 0.814 | 0.791 | 0.857 | 0.752 |  |  |  |
| P52333 (Y) | 23 | 0.805 | 0.76 | 0.724 | 0.805 | 0.558 |  |  |  |
| Q01279 (Y) | 26 | 0.834 | 0.784 | 0.745 | 0.834 | 0.719 |  |  |  |
| Q05397 (Y) | 29 | 0.868 | 0.828 | 0.814 | 0.868 | 0.765 |  |  |  |
| Q06187 (Y) | 22 | 0.847 | 0.805 | 0.78 | 0.847 | 0.63 |  |  |  |
| Q07014 (Y) | 18 | 0.788 | 0.746 | 0.714 | 0.788 | 0.706 |  |  |  |
| Q08881 (Y) | 15 | 0.8 | 0.741 | 0.693 | 0.8 | 0.58 |  |  |  |
| Q13882 (Y) | 26 | 0.808 | 0.767 | 0.739 | 0.808 | 0.523 |  |  |  |
| Q14289 (Y) | 18 | 0.825 | 0.772 | 0.732 | 0.825 | 0.622 |  |  |  |
| Q15046 (Y) | 38 | 0.908 | 0.895 | 0.895 | 0.908 | 0.938 |  |  |  |
| Q16620 (Y) | 17 | 0.789 | 0.726 | 0.675 | 0.789 | 0.821 |  |  |  |
| Q62120 (Y) | 48 | 0.716 | 0.676 | 0.654 | 0.716 | 0.66 |  |  |  |
| Q62689 (Y) | 22 | 0.773 | 0.731 | 0.713 | 0.773 | 0.618 |  |  |  |
| Q62844 (Y) | 22 | 0.849 | 0.812 | 0.797 | 0.849 | 0.677 |  |  |  |
| Q9WUD9 (Y) | 75 | 0.818 | 0.782 | 0.782 | 0.818 | 0.721 |  |  |  |
